# Aptamer-Enabled Discovery and Clinical Analysis of Exosomal Surface Biomarkers in Hepatocellular Carcinoma

**DOI:** 10.1101/2025.05.28.655095

**Authors:** Haiyan Cao, Da Xu, Yuming Su, Chenshuo Zhang, Siqi Bian, Xueling Yan, Shaojin Li, Hong Xuan, Jinchen Long, Panzhu Yao, Xiyang Liu, Xuan Cheng, Huang Su, Yifan Chen, Yike Li, Jun Li, Wanlu Shi, Kun Wang, Baocai Xing, Liqin Zhang

## Abstract

Exosomal membrane proteins hold promise as cancer biomarkers but face detection challenges due to low abundance and conformational sensitivity. We developed AptEx-ID, a microbead-displayed aptamer platform enabling high-throughput screening of hepatocellular carcinoma (HCC)-specific exosomal surface proteins. Aptamers, selected via competitive binding to tumor-derived versus healthy exosomes, demonstrated high affinity and specificity. Clinical validation using 50 HCC and 30 control plasma samples identified a 7-aptamer panel via machine learning, achieving 93.75% accuracy and 0.95 ROC-AUC. These aptamers further revealed HCC-associated biomarkers (e.g., IMPDH1, ASCC3, GTSE1), validated by siRNA knockdown and biophysical assays. Post-treatment monitoring showed aptamer signal reduction in responsive patients, highlighting their prognostic utility. This platform integrates biomarker discovery, diagnostics, and therapeutic monitoring, offering a robust approach for HCC management through exosomal surface proteome profiling.

## Introduction

Exosomes are a promising source of disease biomarkers, reflecting the physiological and pathological states of their parent cells and offering insights into both cellular function and systemic conditions^1–4^. While exosomal RNAs are widely studied, membrane proteins, which play key roles in signal transduction and immune recognition, provide more direct and reliable insights into disease^5–7^. These proteins are accessible on the exosomal surface but are challenging to isolate and study due to their hydrophobic nature, low abundance, and dependence on membrane integrity^8^. Traditional proteomic tools, such as mass spectrometry, often lack the sensitivity to detect these proteins, and antibodies developed against recombinant proteins may fail to recognize them in their native state^9,10^. This has hindered large-scale discovery and validation of exosomal membrane protein biomarkers.

Although surface proteins such as GPC-1, HER2, and EGFR have shown promise, relying on single biomarkers is insufficient due to the complexity and heterogeneity of diseases^11–13^. There is a clear need for new analytical strategies capable of systematically identifying and profiling multiple exosomal membrane proteins to improve diagnostic accuracy, prognostic value, and treatment monitoring^14–17^.

Aptamer technology offers a solution, particularly for targeting cell surface markers. In situ selection methods enable aptamers to bind directly to live cell surfaces, facilitating the discovery of aptamers that can distinguish between cell types and identify novel markers^18–25^. These aptamers serve as molecular tools for biomarker discovery, enabling more precise disease classification.

Aptamers offer several advantages: they bind membrane proteins in their native conformation, avoiding issues that arise with antibodies developed against recombinant proteins, such as altered binding due to post-translational modifications^26–29^. Their synthetic nature allows for chemical modifications to improve affinity, specificity, and stability^30–32^, and they can be easily integrated into various reporting systems for sensitive detection, making them versatile tools for diagnostics and high-throughput screening^33–36^.

In our study, we developed AptEx-ID (microbead-displayed Aptamer screening for Exosomal biomarker IDentification), a platform that screens aptamer libraries bound to intact exosomes to identify tumor biomarkers. We focused on hepatocellular carcinoma (HCC), a highly lethal cancer with no effective early diagnostic tools. Existing biomarkers, such as alpha-fetoprotein (AFP), have limited sensitivity, and despite progress in immunotherapy, many patients exhibit poor response and rapid progression, compounded by HCC’s tumor heterogeneity. AptEx-ID enabled the identification of HCC-specific exosome-binding aptamers, followed by target deconvolution to reveal tumor-specific surface biomarkers (**Fig. 1a**). These aptamers were then used to create a sensitive analytical system, validated with blood samples from HCC patients, leading to the discovery of a series of exosomal biomarkers specific to HCC.

**Fig. 1.**
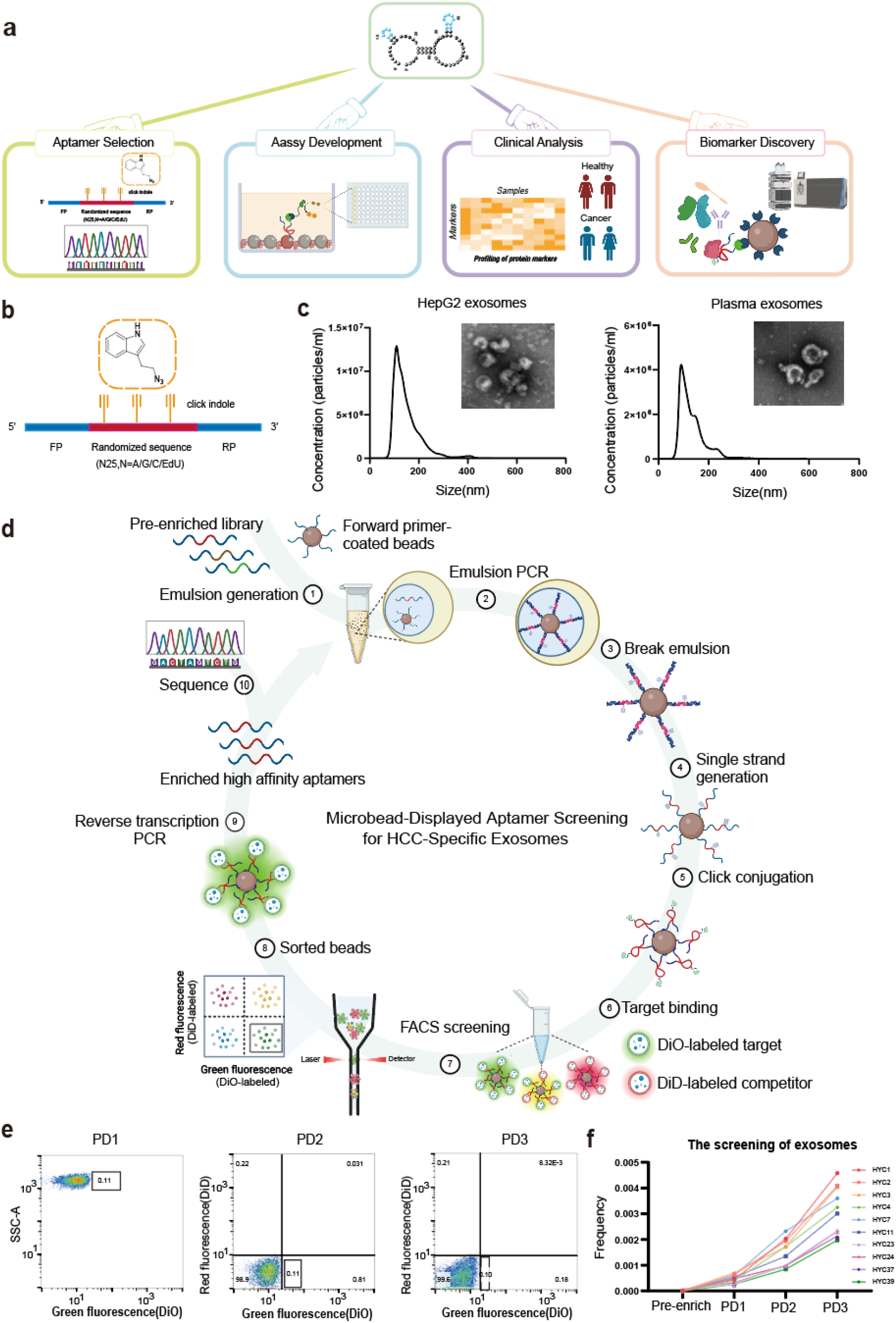
Schematic of the microbead-displayed aptamer screening strategy for targeting HCC-specific exosomes. **a** Manuscript Summary: Aptamers simultaneously perform four key functions: selection, assay development, biomarker discovery, and clinical analysis, showcasing their broad potential in research and clinical applications. **b** Initial Aptamer Library Construction: Creation of the library for in vitro selection. **c** Characterization of Exosomes: Analysis of HepG2 and healthy volunteer plasma-derived exosomes using NTA and cryo-TEM imaging. **d** Microbead-Displayed Aptamer Screening: ① Preparation of the pre-enriched library and forward primer-coated microbeads for emulsion generation. ② Emulsion PCR generates aptamer particles, each displaying multiple copies of a single sequence. ③ & ④ Emulsions are broken, and aptamer particles are converted to single-stranded DNA via NaOH treatment. ⑤ Indole modifications are introduced into the library using click chemistry. ⑥ Indole-modified magnetic microbeads are incubated with fluorescently labeled exosomes: tumor-derived exosomes are labeled with DiO and healthy exosomes with DiD. ⑦& ⑧ FACS sorts aptamer particles with strong DiO fluorescence (quadrant IV). ⑨ & ⑩ Aptamers are reversely converted to natural DNA for further screening or high-throughput sequencing analysis. **e** FACS Sorting: Three rounds of sorting green fluorescent microbeads during aptamer screening. **f** Frequency of Enriched Species: Sequencing data showing the frequency of enriched species from the three rounds of exosome screening.

## Results

### Development of microbead-displayed aptamer screening strategy targeting HCC-specific exosomes

We first leveraged the unique advantage of aptamer screening to directly target the intact surface of exosomes and distinguish tumor-derived vesicles. Although traditional SELEX methods have been applied to exosome targets, their low specificity and complex procedures have limited their broader utility^37,38^. To address this, we adapted the high-throughput efficiency of microbead-display aptamer screening ^39,40^. specifically for exosome surface recognition.

The initial DNA library was designed with a 25-nt randomized region containing an equimolar distribution of four nucleotides, where thymidine (dT) was replaced by 5-ethynyl-2′-deoxyuridine (EdU). This region was flanked by two 21-nt fixed sequences composed of standard nucleotides (**Fig. 1b**) ^41,42^. Taking advantage of aptamer chemical flexibility, we synthesized an azido-indole derivative and employed click chemistry to convert EdU into indole-modified uridine (IndU), which mimics the tryptophan side chain and enhances the potential for high-affinity aptamer binding (**Supplementary Fig. S1a–d**) ^43–45^. Tumor-derived exosomes and healthy plasma-derived exosomes were isolated by ultracentrifugation for screening (**Fig. 1c, Supplementary Fig. S1e**).

Microbead-displayed aptamer libraries were generated via emulsion PCR (**Supplementary Fig. S2a–b**). The optimal template concentration was determined using an Alexa Fluor 633-labeled complementary strand, allowing real-time monitoring of microbead-based library formation. Flow cytometry analysis revealed a 15–30% shift in fluorescence intensity, confirming a high degree of monoclonality (**Supplementary Fig. S2b**). Indole modifications were introduced via click chemistry to complete the library design (**Supplementary Fig. S2c**).

Selection began with two rounds of pre-enrichment against exosomes derived from HepG2 cells. The main screening workflow included: (1) emulsion PCR to create microbeads displaying multiple copies of individual aptamer sequences, (2) emulsion breaking and recovery of single-stranded DNA, (3) click-mediated indole modification, (4) incubation with fluorescently labeled exosomes, (5) FACS sorting of green-fluorescent microbeads, (6) “reverse” PCR amplification of the selected sequences, and (7) repetition of this process for three total rounds followed by deep sequencing (**Fig. 1d, Supplementary Fig. S3a–d**).

A critical step in ensuring the specificity of the selected aptamers was the implementation of a dual-labeling strategy—HepG2 exosomes with DiO (green) and healthy plasma exosomes with DiD (red). Only DiO-positive microbeads were sorted during each FACS round (**Fig. 1e**). Roughly 0.1% of microbeads were selected per round, and “reverse” PCR confirmed successful enrichment (**Supplementary Fig. S3c–d**). Deep sequencing revealed a set of enriched, high-affinity aptamers after three selection cycles (**Fig. 1f, Supplementary Fig. S3e**).

### Characterization of aptamers selected against HepG2 cell-derived exosomes

Sequencing analysis identified 20 top-ranked aptamer candidate clusters, which were synthesized and evaluated for binding to HepG2 exosomes versus healthy plasma-derived exosomes (**Supplementary Fig. S4a–b**). Candidates were PCR-amplified onto microbeads and modified with indole (**Fig. 2a**). Upon incubation with DiO-labeled HepG2 exosomes, most showed a clear fluorescence shift in flow cytometry, indicating strong binding. In contrast, no significant signal was observed with healthy plasma exosomes, comparable to the random sequence control (**Fig. 2b, Supplementary Fig. S4d**). The top 10 aptamers with selective binding to HepG2 exosomes are shown in **Fig. 2c**.

**Fig. 2.**
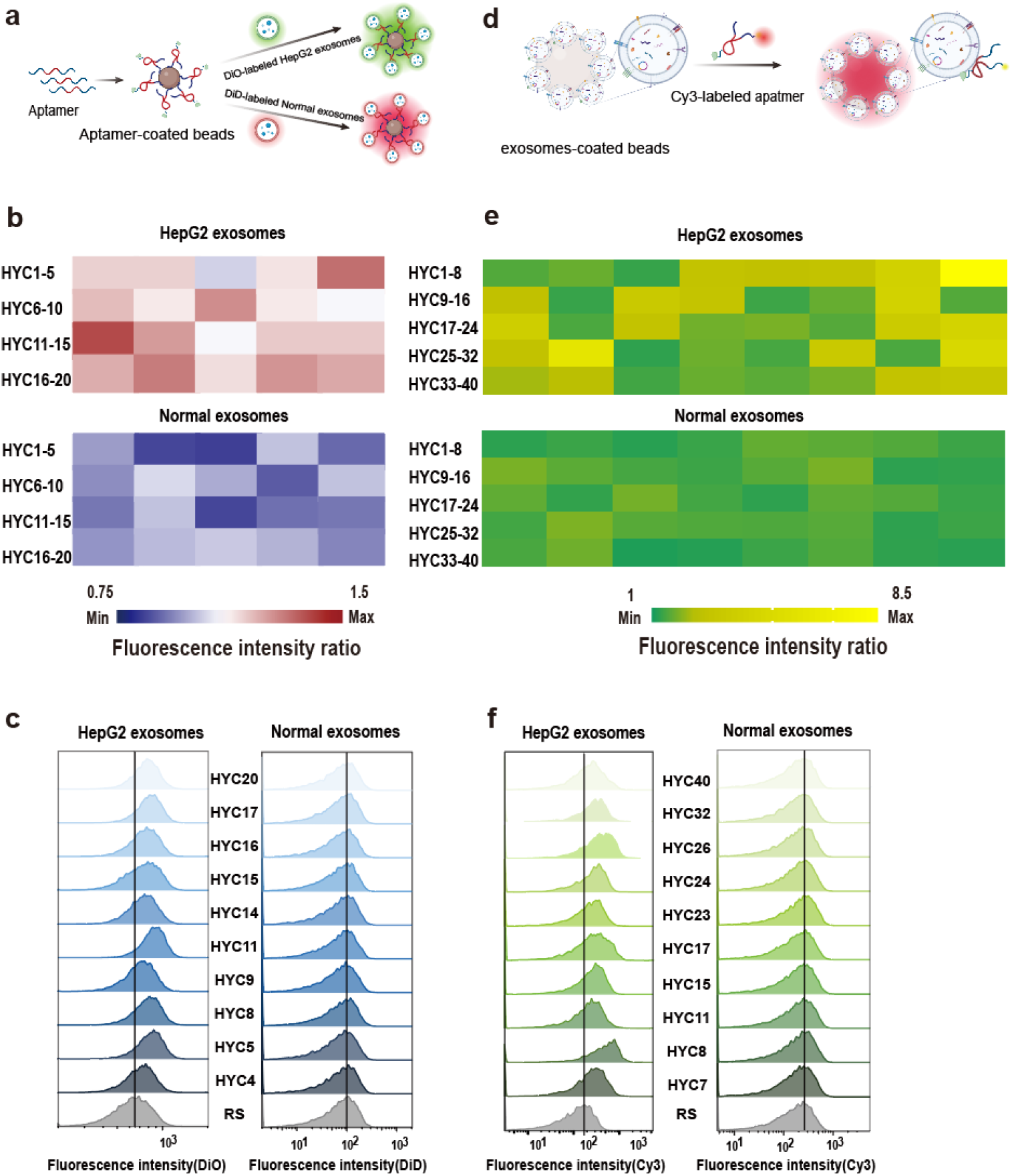
Characterization of Aptamer Binding to Exosomes. **a** Binding Validation Overview: Schematic showing the incubation of aptamer-coated microbeads with exosomes for binding validation. **b** Fluorescence Intensity Heatmaps: Heatmaps illustrating the fluorescence intensity ratios of aptamers compared to random sequences on normal human and HepG2 exosomes, assessed by magnetic microbead validation. **c** Flow Cytometry Analysis: Flow cytometry confirms specific binding of aptamers to HepG2 exosomes, with minimal binding to normal human plasma-derived exosomes, as evaluated using magnetic microbeads. **d** Exosome-Coated Latex Microbeads Binding Validation: Schematic of binding validation using latex microbeads coated with exosomes and Cy3-labeled aptamers. **e** Fluorescence Intensity Heatmaps (Latex Microbeads): Heatmaps showing fluorescence intensity ratios of aptamers versus random sequences on normal human and HepG2 exosomes, validated with latex microbeads. **f** Flow Cytometry with Latex Microbeads: Flow cytometry analysis demonstrates specific binding of aptamers to HepG2 exosomes over normal human plasma-derived exosomes, as assessed using latex microbead validation.

In an expanded screening, the top 40 sequences were tested using Cy3-labeled aptamers incubated with exosome-coated latex microbeads (**Fig. 2d, Supplementary Fig. S4c**). Most showed stronger binding to HepG2 exosomes than to normal exosomes, confirming tumor specificity (**Fig. 2e, Supplementary Fig. S4e**). This two-tier validation confirmed the top 10 high-affinity, tumor-selective aptamers (**Fig. 2f**), demonstrating the efficiency and robustness of our screening platform. The top 40 aptamers were designated as HYC, followed by their enrichment ranking in the sequencing pool.

To elucidate the functional importance of IndU modifications, we conducted systematic mutagenesis studies. Progressive substitution of IndU residues with thymidine (T) demonstrated that specific modification sites were critical for target binding affinity (**Supplementary Fig. S5a**). Furthermore, evaluation of four rationally designed truncation variants by flow cytometry revealed significant differences in target binding capacity, enabling identification of the minimal functional sequence domain required for target recognition (**Supplementary Fig. S5b**).

### Development of purification and analytical methods for clinical sample detection

To apply the selected aptamers could differentiate hepatocellular carcinoma (HCC) exosomes from those derived from healthy plasma to clinical samples, we first optimized both the exosome purification process and aptamer-based detection methods. Given the limited availability and clinical importance of patient plasma, we evaluated several isolation techniques. Ultracentrifugation (UC), the gold standard, was initially employed, with exosome size and morphology confirmed by nanoparticle tracking analysis (NTA) and transmission electron microscopy (TEM) (**Supplementary Fig. S6a**). To minimize lipoprotein contamination, we also tested a modified UC protocol incorporating a deproteinization step. In parallel, alternative methods—including immunoaffinity capture, polymer-based precipitation, and size-exclusion chromatography (SEC)—were systematically compared (**Supplementary Fig. S6b–e**). SEC showed the best performance, with the third elution fraction yielding uniformly sized (∼200 nm) particles by NTA and intact lipid bilayer vesicles confirmed by TEM (**Fig. 3a**), establishing it as the optimal method for plasma exosome isolation.

**Fig. 3.**
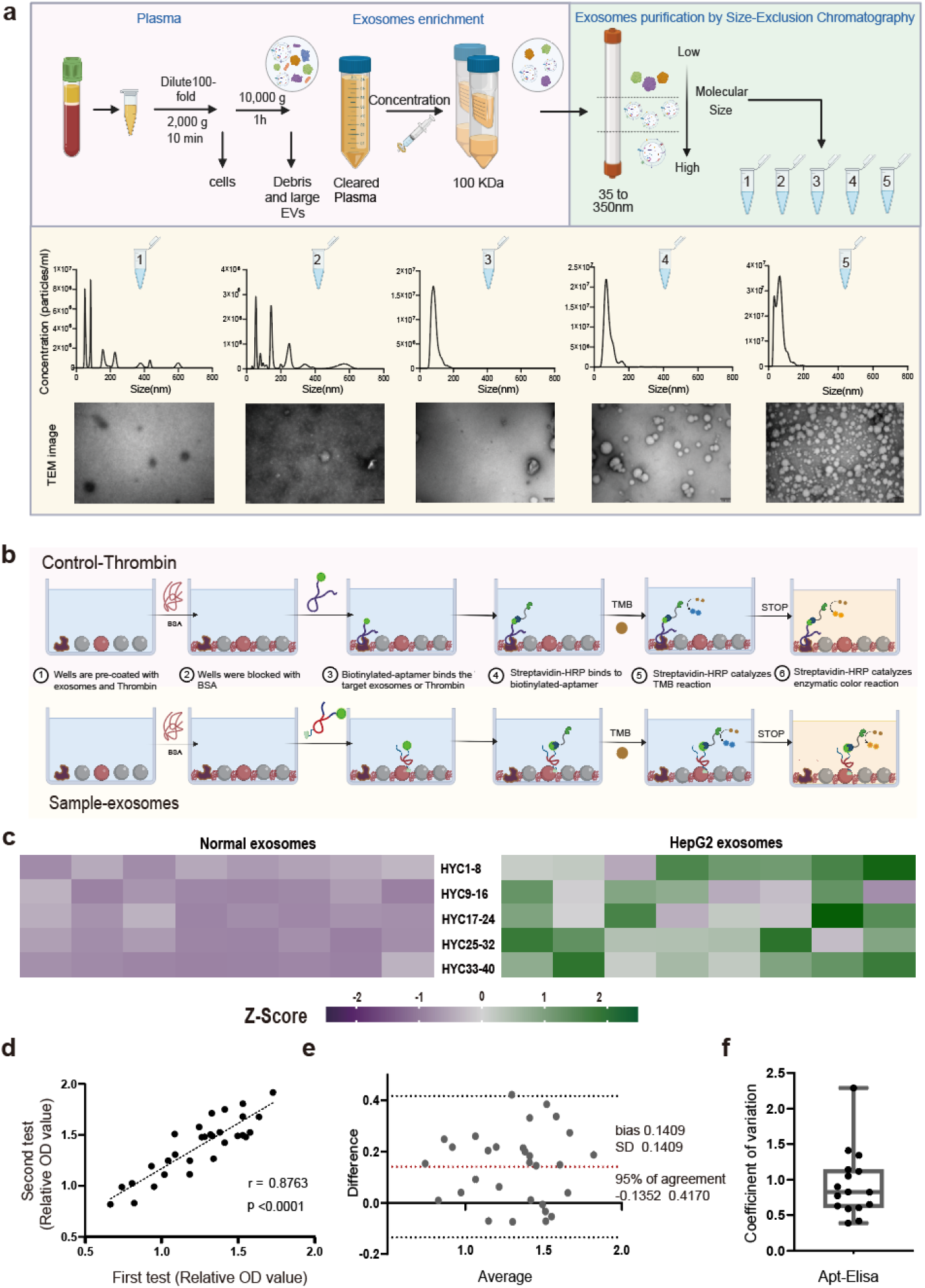
Aptamer-Based ELISA Assay for Extracellular Vesicle Detection. **a** Exosome Isolation and Characterization: Plasma-derived exosomes were isolated using Size-Exclusion Chromatography (SEC), followed by characterization through NTA and cryo-TEM imaging. **b** Aptamer-Based ELISA Development: An aptamer-based ELISA was established to detect exosomes, with thrombin serving as a reference for assay consistency. **c** Z-Score of Absorption Values: Z-scores of the relative absorption at 450 nm for each aptamer from normal human plasma-derived and HepG2 exosomes. **d** Linear Regression: Regression analysis of the first and second test signals from different batches of the same sample, with probe HYC11. Pearson r = 0.8763. **e** Bland-Altman Plot: The Bland-Altman plot compares the consistency of the two detection methods, where a bias value near zero indicates better agreement. Data from normal human plasma-derived exosomes (n=30). **f** Coefficient of Variation: Coefficient of variation (CV) of the Apt-Elisa assay (Aptamer-CD63) for two or more serum samples from the same patient, shown as a box-and-whiskers plot. The CV was 1.98% (n=16).

To enable quantitative analysis of exosomal surface markers, we developed a colorimetric ELISA platform using biotin-labeled aptamers as detection probes. Unlike traditional antibody-based assays, our design leverages the target specificity of aptamers and the strong biotin–streptavidin-HRP interaction for signal amplification. To minimize variability in chromogenic reactions, we incorporated an internal reference system based on the thrombin–TBA29 aptamer pair (**Fig. 3b**), ensuring consistent and reliable quantification across replicates.

To establish a robust detection platform, we systematically optimized detection conditions using three setups: (i) plasma-spiked exosomes followed by recovery, (ii) direct detection in Binding Buffer E, and (iii) detection in exosome-depleted plasma dilutions. Sensitivity analysis in the incubation buffer revealed concentration-dependent responses, identifying the optimal solution for subsequent assays (**Supplementary Fig. S7a**).

We then compared signal amplification strategies, evaluating a commercial TMB chromogenic substrates and an ultra-sensitive TMB formulation (**Supplementary Fig. S7b**), alongside an HRP-based fluorescent assay (**Supplementary Fig. S7c**). The ultra-sensitive TMB system provided superior signal differentiation between tumor and normal plasma exosomes and was chosen for further development. To ensure consistent assay performance, we optimized the internal reference system (thrombin-TBA29) by titrating thrombin concentrations for a matched dynamic range (**Supplementary Fig. S7d**).

The aptamer-based ELISA (Apta-ELISA) platform was used to compare plasma-derived and HepG2-derived exosomes (**Supplementary Fig. S7e**). Z-score normalization and heatmap analysis revealed distinct molecular signatures, highlighting the aptamers’ strong discriminatory ability (**Fig. 3c**).

To assess reproducibility, duplicate measurements were performed on the same samples. Pearson correlation showed a strong positive correlation (r = 0.863, p < 0.0001) (**Fig. 3d**), while Bland-Altman analysis confirmed consistency, with minimal bias and 95% of data within agreement limits (**Fig. 3e**). A low coefficient of variation (CV = 1.98%) further demonstrated assay stability (**Fig. 3f**). These results confirm the reliability and technical robustness of the platform.

### Clinical samples analysis reveals actionable aptamer panel for HCC diagnostic uses

The aptamer selected was then used to identify biomarkers on exosome surfaces. To assess the diagnostic potential of aptamer panels, we tested selected candidates on clinical samples using our optimized analytical platform. Peripheral blood was collected from 50 hospitalized HCC patients (**Fig. 4a**) and 30 healthy volunteers as controls.

**Fig. 4.**
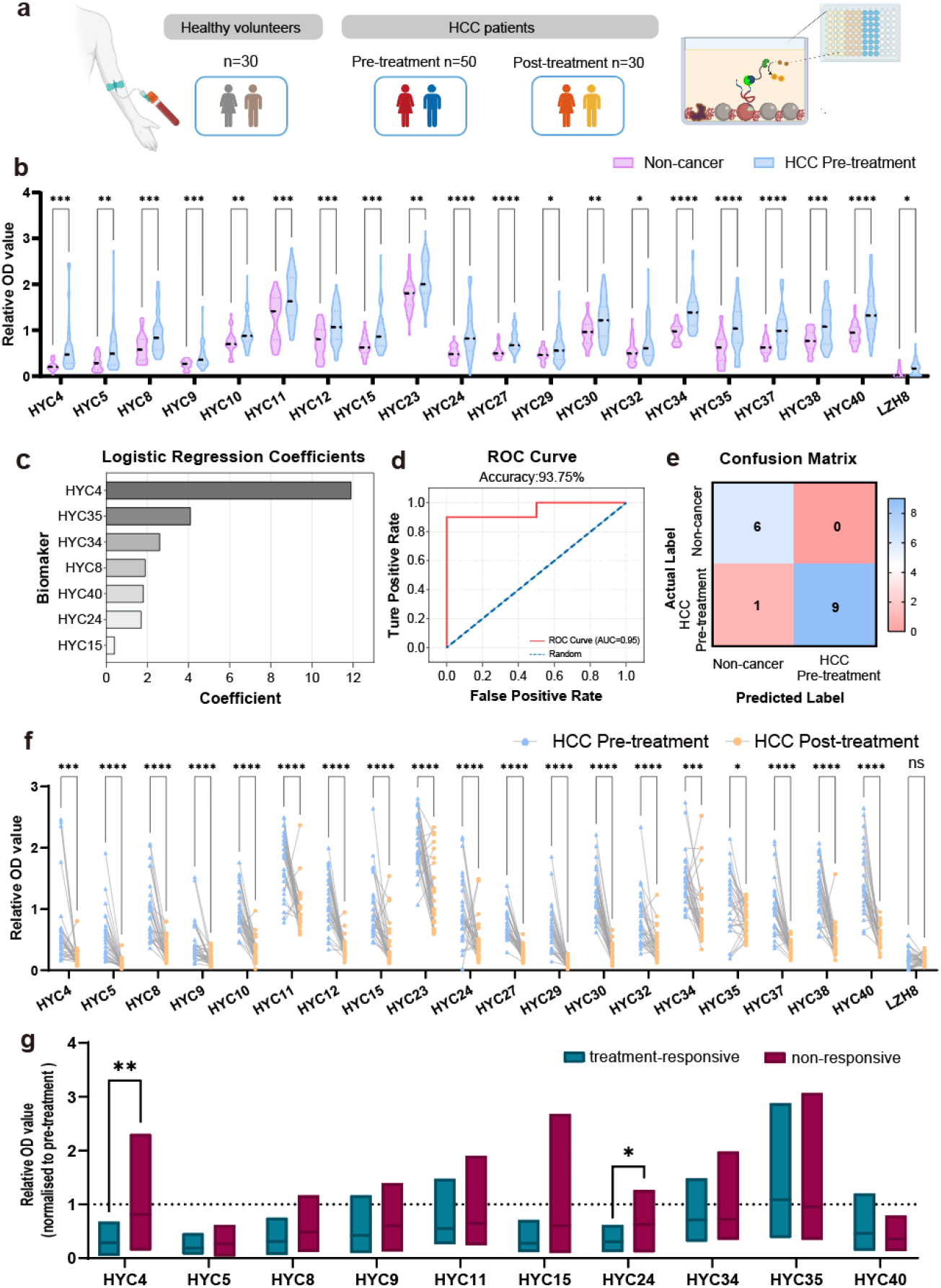
Clinical samples analysis with selected aptamers and established analytical methods. **a** Clinical Application of Liquid Biopsy: Schematic diagram illustrating the potential of liquid biopsy in clinical settings. **b** Relative OD Values of Aptamers: OD values at 450 nm for aptamers from exosomes of healthy volunteers and HCC pre-treatment patients. Non-cancer (n = 30), HCC pre-treatment (n = 50). *p < 0.05; **p < 0.01; ***p < 0.001; ****p < 0.0001. **c** Feature Importance Analysis: Analysis of the relative contribution of each biomarker to the model, with HYC4 showing the highest importance score (11.90). **d** ROC Curve: Receiver Operating Characteristic (ROC) curve of the logistic regression model on the independent test set, achieving an AUC of 0.95. **e** Confusion Matrix: Performance of the model, showing 90% sensitivity and 100% specificity. **f** Paired Analysis of Aptamer-Based ELISA: Comparison of aptamer-based ELISA assay results before and after targeted therapy and immunotherapy in each patient. Dots connected by a line represent a single patient before and after treatment (n = 30). *p < 0.05; ***p < 0.001; ****p < 0.0001; ns: not significant. **g** Comparative Analysis of OD Values: Comparison of relative OD values between treatment-responsive (n = 12) and non-responsive (n = 14) groups. OD values were normalized to baseline, and statistical significance was determined by unpaired t-test (*p < 0.05; **p < 0.01).

Initial screening was conducted using 29 aptamers from the current selection and an additional aptamer, LZH8, which targets GRP78 on HCC-derived exosomes (**Supplementary Fig. S8a**)^46^. Among these, 20 aptamers showed significantly higher relative OD values in HCC patient-derived exosomes compared to those from healthy volunteers, demonstrating strong discriminatory capability (**Fig. 4b**).

We then developed a diagnostic model using machine learning based on data from these 20 aptamers. The workflow included data preprocessing from 80 samples (50 HCC, 30 controls), biomarker (aptamer) selection, feature optimization via recursive feature elimination with cross-validation, model training with hyperparameter tuning, and evaluation on an independent test set (**Supplementary Fig. S8b**).

Statistical analysis confirmed all 20 aptamers were significantly differentially expressed (p < 0.05), with HYC34, HYC24, and HYC40 showing the highest t-values (6.99, 5.64, and 5.57, respectively) (**Supplementary Fig. S9a–b**). Feature selection identified a 7-aptamer signature (HYC4, HYC8, HYC15, HYC24, HYC34, HYC35, HYC40) that achieved optimal classification performance (cross-validation ROC-AUC: 0.967 ± 0.047) (**Supplementary Fig. S9c, S10**). Notably, six biomarkers (HYC4, HYC15, HYC24, HYC34, HYC35, HYC40) were consistently selected across multiple methods, highlighting their robustness.

Feature importance analysis ranked HYC4 as the top predictor (score: 11.90), followed by HYC35, HYC34, HYC8, HYC40, HYC24, and HYC15 (**Fig. 4c**). The final logistic regression model demonstrated strong performance on an independent test set (n=16), with 93.75% accuracy, 100% precision, 94.74% F1-score, and ROC-AUC of 0.95 (**Fig. 4d, Supplementary Fig. S9c**). Confusion matrix analysis showed 100% specificity and 90% sensitivity (**Fig. 4e**), indicating excellent diagnostic potential—minimizing false positives while maintaining high detection accuracy.

### Further analysis to evaluate the potential of the aptamers for HCC immunotherapy monitoring

After the initial blood sample collection, a second time-point sample was collected from each patient following 2–3 cycles of targeted immunotherapy (approximately 40–60 days later). To evaluate early treatment response, we analyzed paired pre- and post-treatment samples for changes in exosome-associated markers.

Among the 20 aptamer probes tested, most showed a significant decrease in marker expression after treatment, indicating therapy-induced modulation (**Fig. 4f**).

Based on clinical follow-up, patients were categorized as either treatment-responsive or non-responsive. Comparative analysis of relative optical density (OD) values between the two time points identified two aptamers—HYC4 and HYC24— that effectively distinguished treatment outcomes by detecting early changes in biomarker expression (**Fig. 4g**). These results highlight the potential of aptamer-based exosome profiling as a valuable tool for early therapeutic monitoring in HCC patients receiving targeted therapy.

### Aptamer-based target elucidation and biomarker identification

After validating the aptamer panel for HCC diagnosis, we aimed to identify binding targets of the seven aptamers on the exosome surface—i.e., HCC-associated exosomal biomarkers. Given that tumor-derived exosomes carry protein signatures reflective of their parent cells, we first confirmed aptamer binding to HepG2 cells via flow cytometry (**Fig. 5a**). Trypsin digestion of membrane proteins prior to aptamer incubation significantly reduced binding (**Supplementary Fig. S11**), confirming that the targets are proteinaceous.

**Fig. 5.**
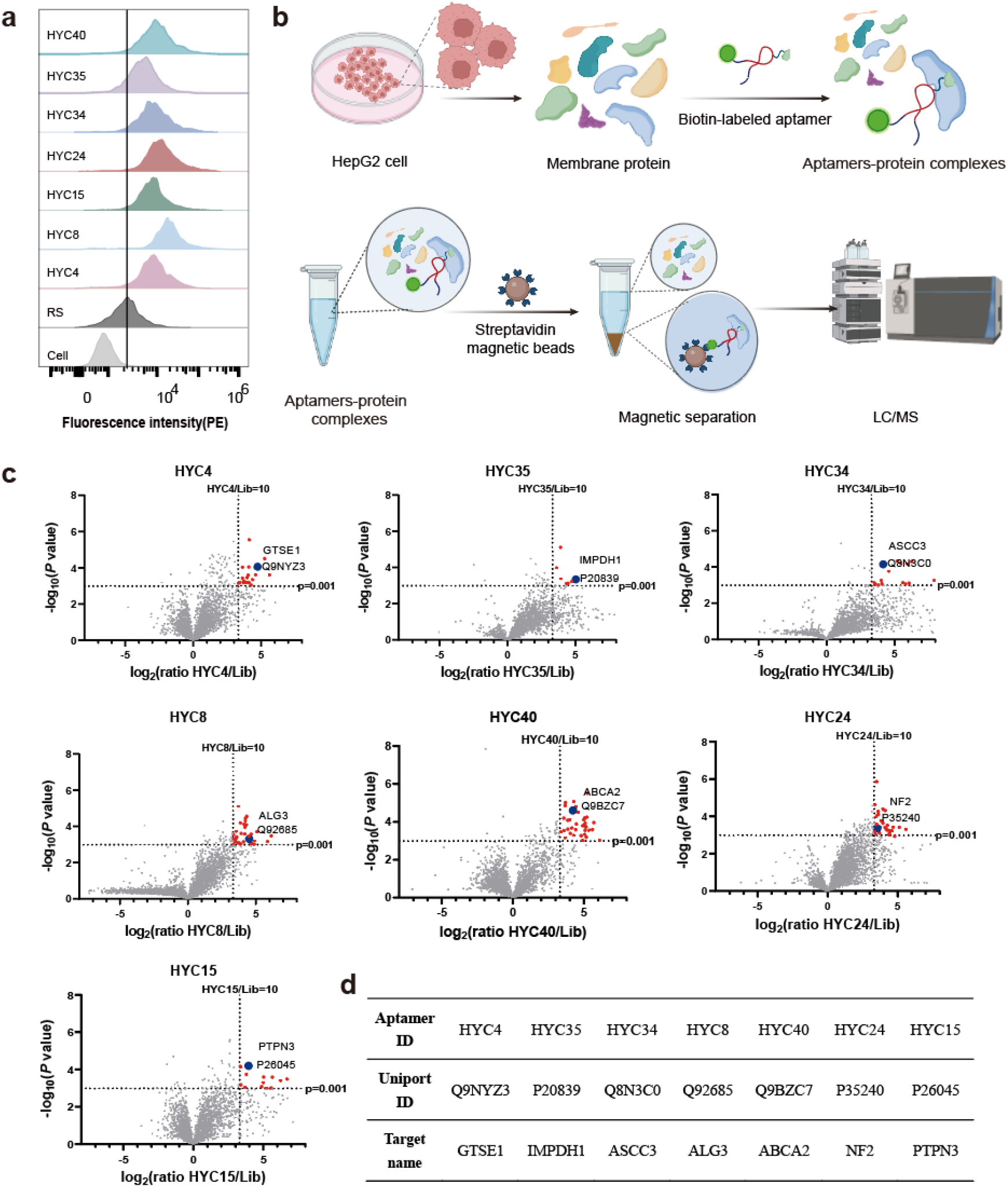
Identification of the binding target of the aptamers. **a** Verify the binding ability of aptamers and HepG2 cells. **b** Schematic illustration of procedures to identify binding target of aptamers. **c** Volcano diagram analysis of differential proteins based on proteomics. log2 (ratio)>3.32, P value>2.99. **d** Characteristics of selected aptamers and their validated protein targets.

For target identification, biotin-labeled aptamers were synthesized, characterized and incubated with HepG2 lysates, and aptamer-protein complexes were purified using streptavidin-coated magnetic beads (**Fig. 5b, Supplementary Fig. S12-13**). These complexes were analyzed by LC-MS/MS, and differentially enriched proteins were identified through comparative proteomics (**Fig. 5c**). Through systematic analysis of the available data and integration of the target proteins’ membrane localization, membrane interaction properties, and cancer relevance, we prioritized and identified the candidate target proteins for further study (**Fig. 5d**).

To assess the clinical relevance of the identified targets, we conducted comprehensive bioinformatics analyses using TIMER2.0 (http://timer.comp-genomics.org/) and GEPIA2.0 (http://gepia.cancer-pku.cn/). Differential expression analysis showed that all validated markers were significantly upregulated in LIHC (Liver hepatocellular carcinoma) tissues compared to adjacent non-tumor tissues (**Supplementary Fig. S14**). Survival analysis revealed that high expression levels of these markers were strongly associated with poorer overall survival in LIHC patients (**Supplementary Fig. S15**). Additionally, ExoCarta database (http://www.exocarta.org/) interrogation confirmed that nearly all candidate biomarkers had been previously detected in extracellular vesicle (EV) proteomic studies, supporting their potential as EV-based diagnostic markers.

IMPDH1, a known regulator of p53-dependent growth, is frequently overexpressed in various cancers^47,48^. ASCC3, a DNA helicase involved in damage repair and immune modulation, has been increasingly linked to solid tumor development ^49,50^. GTSE1, a p53-responsive gene expressed during the S and G2 phases, plays a key role in the DNA damage response and is implicated in multiple malignancies ^51,52^. PTPN3 (PTPH1), a non-receptor protein tyrosine phosphatase, displays both tumor-suppressive and oncogenic properties depending on the cancer type, highlighting its tissue-specific roles^53–56^.

The tumor suppressor gene NF2 encodes merlin, a membrane-associated protein. While NF2 mutations are well-documented in schwannomas and meningiomas, paradoxical overexpression in certain cancers suggests a context-dependent tumor-suppressive role (**Supplementary Fig. S12**) ^57^, paradoxical NF2 overexpression has been observed in certain cancer patients, ABCA2, an ABC transporter, contributes to multidrug resistance and has been linked to disease progression in various cancers^58,59^. Lastly, ALG3, an ER-resident enzyme in N-glycan biosynthesis, is markedly overexpressed and associated with tumorigenesis in several cancers, including hepatocellular carcinoma^60,61^.

### Functional Validation of Aptamer-Target Interactions

Antibodies against these proteins were tested on the HepG2 cell surface to confirm the presence of these proteins **(Supplementary Fig. S16a)**. Before validating target proteins of the aptamers, we optimized transfection conditions for HepG2 cells. Flow cytometry showed that Lipofectamine 3000 provided the highest transfection efficiency (**Supplementary Fig. S16b**) and was used for all gene silencing experiments. siRNA-mediated knockdown of candidate proteins led to reduced mRNA and protein levels, confirmed by qPCR and Western blot. Correspondingly, flow cytometry revealed decreased aptamer binding, demonstrating a clear inverse correlation between target expression and aptamer affinity (**Fig. 6, Supplementary Fig. S17a**), supporting specific aptamer-target interactions.

**Fig. 6.**
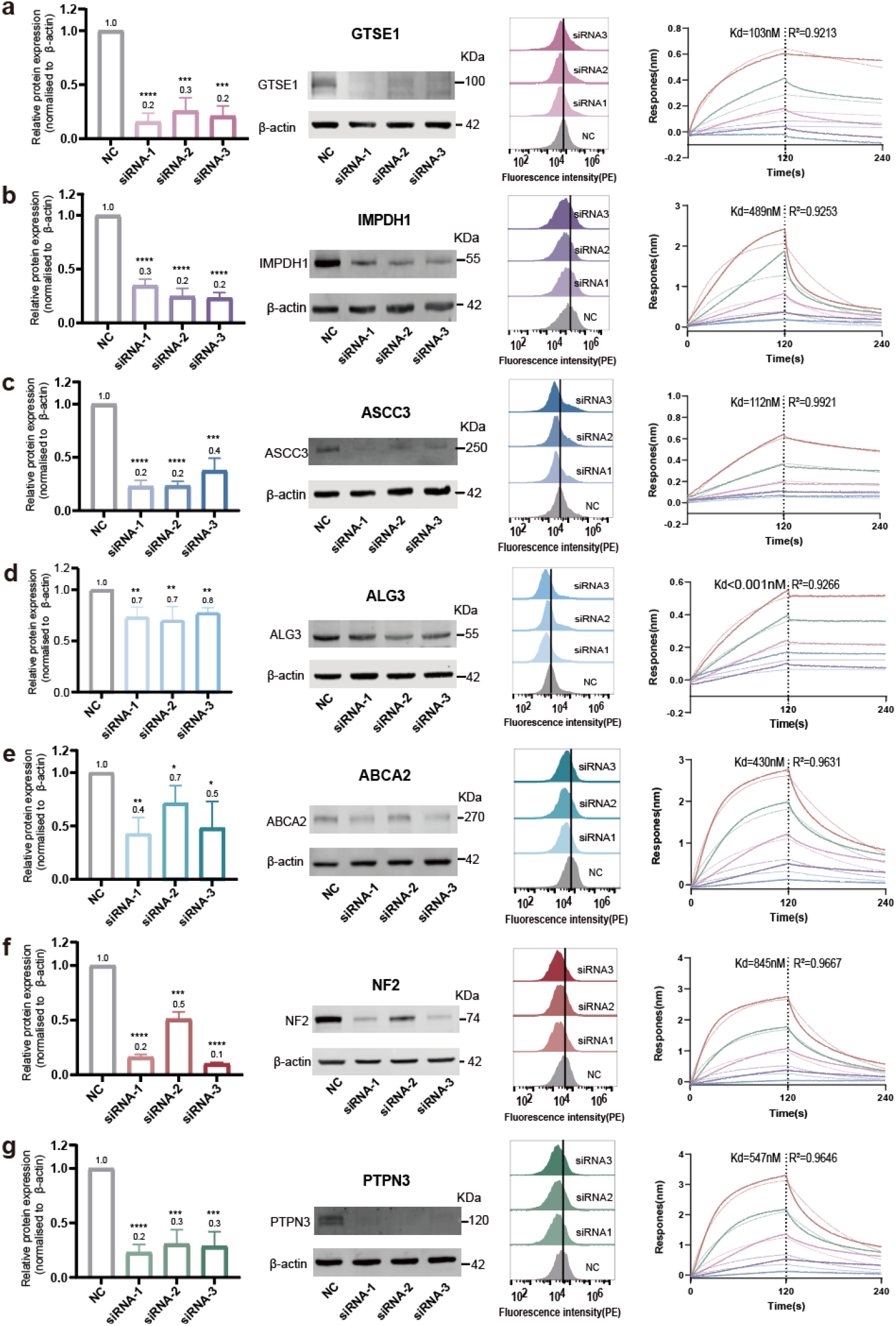
Molecular validation of aptamer-binding targets. (**a**-**g**) From top to bottom are the target proteins GTSE1, IMPDH1, ASCC3, ALG3, ABCA2, NF2, and PTPN3. The verification of each target protein from left to right is as follows: Typical WB images and quantitative analysis showing that the protein expression level was reduced by siRNA (n = 3, *p < 0.05; **p < 0.01; ***p < 0.001; ****P < 0.0001; unpaired Student’s t test); Flow cytometry analysis of the binding ability of aptamer on knockdown HepG2 cells; and the BLI assay determining the Kd value of targets and aptamers.

To further characterize these interactions, we used bio-layer interferometry to assess direct binding between aptamers and recombinant target proteins. The resulting dissociation constants indicated high-affinity interactions (**Fig. 6, Supplementary Fig. S17b**). No binding was observed with scrambled sequences, confirming specificity. These biophysical measurements aligned with cellular data, reinforcing the validity and functional relevance of the aptamer-protein interactions. Structural predictions were also performed to understand the molecular basis of aptamer-target interactions. Secondary structure analysis and tertiary structure simulations revealed distinct conformational features of the IndU-modified aptamers (**Supplementary Fig. S18 and S19**). Molecular docking simulations further predicted potential interaction interfaces between the modified aptamers and their target proteins (**Supplementary Fig. S19**).

In addition to the seven predicted biomarkers, three additional aptamer targets—HYC5, HYC9, and HYC11—were identified and validated to evaluate the robustness of this method **(Supplementary Fig. S20 and 21)**. This approach successfully identified three more potential biomarkers that differentiate HCC-derived exosomes from healthy exosomes.

## Discussion

In this study, we developed an aptamer surface selection–enabled platform for the discovery of exosomal membrane protein or membrane-associated protein biomarkers. This approach screens microbead-displayed aptamer library targeting exosomal surface proteome that enables rapid, high-specificity identification of native protein targets directly from intact exosomes. The platform offers a powerful tool for unveiling functional surface biomarkers with potential applications in diagnostics, prognostics, and therapeutic monitoring.

Notably, our microbead-displayed aptamer selection method produced aptamers with exceptional specificity, outperforming traditional SELEX and EV-SELEX approaches. Impressively, this high level of specificity was achieved in just three rounds of selection, underscoring both the efficiency and effectiveness of the strategy. Through this approach, we rapidly identified seven candidate surface biomarkers within a markedly shortened experimental timeframe, demonstrating the platform’s strong potential for high-throughput biomarker discovery. Such efficiency has not been observed with other molecular probes, including antibodies. In fact, constructing a discovery platform targeting the exosomal membrane proteome using conventional probes remains highly challenging.

Although we validated our method using a cohort of 50 clinical samples, we acknowledge that the sample size is limited. The machine learning models built upon this dataset serve primarily as proof-of-concept demonstrations rather than fully optimized predictive tools. Nevertheless, these preliminary results underscore the platform’s capability to rapidly identify disease-associated targets. With access to larger clinical cohorts and extended follow-up periods, we believe this platform holds great promise for biomarker discovery in clinical contexts—ranging from early diagnosis and prognosis to monitoring therapeutic response.

Given the broad relevance of exosomes across various disease states, including numerous cancers, neurodegenerative disorders, and inflammatory conditions, this platform could be readily adapted and expanded to other disease models. Its modular nature allows for customization based on disease-specific exosomal profiles, paving the way for broader translational impact.

Furthermore, by integrating this aptamer-based discovery pipeline with high-throughput DNA chip technologies, it is possible to build an ultramultiplexed exosomal surface proteomics system. Such a platform could enable parallel profiling of dozens to hundreds of biomarkers, greatly enhancing analytical resolution and providing deeper insights into disease heterogeneity.

While we successfully identified putative protein targets in this study, questions remain regarding their specificity and uniqueness to disease states. Importantly, aptamers themselves—independent of their exact molecular targets—can serve as functional biomarkers. Their binding profiles often correlate with the expression levels of specific surface proteins but, more critically, reflect the physiological or pathological state of the originating cells. Thus, aptamer panels may provide valuable readouts of disease progression or therapeutic response, even in the absence of complete molecular annotation.

## Materials and Methods

### Study Design and Clinical Sample Collection

This study aimed to identify hepatocellular carcinoma (HCC)-specific exosomal biomarkers using a structure- and function-guided aptamer screening platform, followed by validation in patient-derived samples. A total of 50 unresectable HCC patients and 30 healthy donors were enrolled at Peking University Cancer Hospital under IRB approval (2023KT129), with written informed consent. Blood samples were collected at baseline and during treatment (Day 40–60), processed to obtain plasma, and stored at –80°C. Inclusion criteria for patients included confirmed HCC diagnosis (CNLC stage Ia–IIIb), no prior systemic therapy, Child-Pugh ≤7, ECOG 0 – 1, and no significant comorbidities. Treatment regimens included immune checkpoint inhibitors and anti-angiogenic agents, and tumor responses were evaluated by mRECIST.

### Cell Lines and Culture

HepG2 hepatoma cells (ATCC® HB-8065™) were cultured in DMEM supplemented with 10% FBS at 37°C in a 5% CO_2_ incubator. Cells were passaged using trypsin-EDTA or enzyme-free dissociation solution and resuspended in appropriate buffer for downstream applications.

### Exosome Isolation and Characterization

Exosomes were isolated from both HepG2 cell culture media and human plasma via multiple methods. Tumor cell-derived exosomes were collected after serial centrifugation (300×g, 2000×g, 10,000×g) and ultracentrifugation at 100,000×g for 3 hours. Plasma-derived exosomes were prepared by similar ultracentrifugation or via size-exclusion chromatography (SEC), magnetic bead immunoprecipitation, or polymer-based precipitation. Exosome morphology was assessed by transmission electron microscopy (TEM), and size and concentration were measured by nanoparticle tracking analysis (NTA; NanoSight NS300).

### Exosome Labeling and Pre-enrichment of Aptamers

Exosomes from tumor and normal plasma were differentially labeled with DiO and DiD membrane dyes, respectively. Pre-enrichment of aptamers was performed using a cell-SELEX approach with a 25N randomized ssDNA library. Libraries were chemically modified with indole moieties via click chemistry and incubated with exosome-latex microbead complexes for negative and positive selection. Two rounds of pre-enrichment were conducted to enhance specificity before high-throughput screening.

### Microbead-Based Aptamer Screening (AptEx-ID)

Aptamer libraries were immobilized onto magnetic microbeads via emulsion PCR, enabling high-throughput functional screening. Indole-modified aptamer-displaying beads were incubated with DiO-labeled tumor-derived and DiD-labeled normal exosomes. Following competitive binding, fluorescence-based flow cytometry was used to isolate aptamer-bound complexes with preferential binding to tumor exosomes. Selected aptamer sequences were amplified, barcoded, and sequenced using the Illumina NovaSeq platform. Sequence clustering and enrichment analysis were conducted using Aptasuite to identify candidate aptamers.

### Aptamer Validation by Flow Cytometry

Binding specificity of selected aptamers was assessed via flow cytometry. Biotin-or Cy3-labeled aptamers were incubated with HepG2 cells, DiO-labeled tumor-derived exosomes, or DiD-labeled normal exosomes. Streptavidin-PE or direct fluorescence readouts were analyzed using a Cytoflex flow cytometer (Beckman), with FlowJo software. Random sequences served as negative controls.

### Apta-ELISA for Exosome Detection

Quantification of target exosomes was performed using an aptamer-based ELISA (Apta-ELISA). Exosomes (10^7^ particles) were immobilized on high-binding 96-well plates, blocked, and incubated with 250 nM biotinylated aptamers. After washing, SA-HRP was added, and color development was performed using TMB substrate. OD_450_ values were measured using a microplate reader. Exosome detection was further optimized using thrombin as an internal reference standard, and the aptamer-based detection system was validated across varying concentrations and matrix backgrounds.

### Target Identification of Aptamers

Membrane proteins from HepG2 cells were extracted and incubated with biotin-labeled aptamers. The resulting aptamer-protein complexes were pulled down using streptavidin magnetic beads, separated by SDS-PAGE, and visualized by silver staining. Protein bands were excised and analyzed by LC-MS (Orbitrap Velos Pro) to identify aptamer targets.

### Biolayer Interferometry (BLI) for Binding Affinity

Binding affinity between aptamers and recombinant target proteins was measured by biolayer interferometry using a ForteBio system. Biotinylated aptamers were immobilized on streptavidin biosensors, and binding kinetics were analyzed to calculate dissociation constants (Kd).

### Molecular Modeling and Structure Prediction

Three-dimensional structures of aptamers were predicted using the 3dRNA/DNA webserver and refined via molecular dynamics (MD) simulations using the AMBER 16 software suite. Indole-EdU modification parameters were derived from quantum chemical calculations. Protein structures were obtained from PDB or predicted using AlphaFold. Docking simulations for aptamer-protein complexes were performed with HDOCK, and the top-scoring conformations were selected based on docking scores and RMSD.

### Functional Validation via siRNA Knockdown

HepG2 cells were transfected with siRNAs targeting candidate proteins identified from aptamer pull-down experiments. Forty-eight to seventy-two hours post-transfection, cells were analyzed by flow cytometry and Western blot to evaluate knockdown efficiency and downstream effects.

### Western Blotting

Proteins extracted from HepG2 cells or exosomes using RIPA buffer were quantified by BCA assay, separated by SDS-PAGE, and transferred to PVDF membranes. Primary antibodies targeting candidate biomarkers, exosomal markers (TSG101, CD63, CD81), and controls (β-actin, Calnexin) were used. Detection was performed using IRDye-conjugated secondary antibodies and visualized via the Odyssey imaging system.

### Quantitative RT-PCR

Total RNA was extracted from cells and tissues using standard protocols, reverse-transcribed into cDNA, and analyzed via qRT-PCR using SYBR Green Master Mix. Gene expression levels were normalized to ACTB and quantified using the 2^ΔΔCt^ method.

### Differential Expression and Survival Analysis

Candidate biomarkers were evaluated for differential expression and clinical relevance using public datasets. GEPIA2 and TIMER web platforms were used to assess associations between biomarker expression and overall survival (OS), disease-free survival (DFS), and tumor stage. Hazard ratios (HRs) were calculated to quantify the prognostic significance.

### Machine Learning for Diagnostic Model Development

To evaluate the diagnostic potential of identified biomarkers, we analyzed data from 80 subjects (50 HCC, 30 controls) across 31 features. Stratified sampling was used to divide the dataset into training (n=64) and testing (n=16) sets. Feature selection included Welch’s t-test and recursive feature elimination (RFECV) using logistic regression. Several classification models were evaluated (Naive Bayes, KNN, SVM, Decision Tree, Random Forest, XGBoost), with L1-regularized logistic regression selected for its performance and interpretability. Hyperparameters were tuned using 8-fold cross-validation with ROC-AUC as the optimization metric. Final performance was evaluated using accuracy, precision, recall, specificity, F1-score, and ROC-AUC on the independent test set.

### Statistical Analysis

All statistical analyses were conducted using GraphPad Prism (versions 8.0 and 10.0). Box plots display the full data range with interquartile range and mean. For comparisons among multiple groups, one-way ANOVA was applied; pairwise comparisons used t-tests. Statistical significance was denoted as follows: p ≤ 0.05 (*), p ≤ 0.01 (**), p ≤ 0.001 (***), p ≤ 0.0001 (****); p > 0.05 was considered not significant (ns).

## Supporting information

Supplementary Information for Aptamer-Enabled Discovery and Clinical Analysis of Exosomal Surface Biomarkers in Hepatocellular Carcinoma

## Competing interests

Authors declare that they have no competing interests.

## Data and materials availability

All data are available in the main text or the supplementary materials.

## Acknowledgments

We would like to thank the Key Laboratory of Carcinogenesis and Translational Research, Hepatopancreatobiliary Surgery Department I for HCC samples and non-HCC samples. We also extend our thanks to the staff at the Peking University Medical and Health Analysis Center and the State Key Laboratory of Natural and Biomimetic Drugs for their invaluable assistance with instrumental analysis.

## Funding

This work was supported by the National Key Research and Development Program of China [2023YFF1205902, 2022YFA1304501 to L.Z. and 2023YFC3405100 to D.X.]; National Natural Science Foundation of China [22227805, 22374004 to L.Z. and 82303727 to D.X.], Science Foundation of Peking University Cancer Hospital [JC202404 to D.X. and L.Z.], Capital Medical Science and Technology Innovation Achievement Transformation and Promotion Program [YC202301JS0083 to K.W.].

## Author contributions

Conceptualization: H.C., D.X., K.W., B.X., and L.Z. Methodology: H.C., Y.S., D.X. and L.Z. Formal analysis: H.C., Y.S. and C.Z. Investigation: H.C., Y. S., C.Z., S.B., X.Y., S.L., H.X., P.Y., X.L., X.C., H.S., Y.C., Y.L., and J.L. Funding acquisition: D.X., K.W. and L.Z. Supervision: D.X., K.W., B.X. and L.Z. Validation: H.C., Y.S., D.X. and L.Z. Visualization: H.C., and Y.S. Writing—original draft: H.C. and L.Z. Writing—review & editing: D.X. and L.Z.

## Notes

### Competing Interest Statement

The authors have declared no competing interest.

