## Supplementary Information for Aptamer-Enabled Discovery and Clinical Analysis of Exosomal Surface Biomarkers in Hepatocellular Carcinoma for "Aptamer-Enabled Discovery and Clinical Analysis of Exosomal Surface Biomarkers in Hepatocellular Carcinoma"

**The PDF file includes:**

Supplementary Materials and Methods

Figs. S1 to S21

Tables S1 to S10

**Supplementary Materials and Methods**

**Materials**

Reagents for Oligonucleotide Synthesis Reaction, including DMT-dC (Bz)-CE phosphoramidite, DMT-dG (iBu)-CE phosphoramidite, and DMT-dA (Bz)-CE phosphoramidite, as well as 5'-Biotin CE Phosphoramidite were acquired from Wuhu Huaren Science and Technology. The DMT-5-Ethynyl-2’-deoxyuridine-3’-CE phosphoramidite was purchased from Nanjing Huijutong Biotechnology Co., Ltd. The 2'-deoxyribonucleotide triphosphate dNTP (set of dATP, dTTP, dGTP, dCTP, 5-Ethynyl-dUTP) were from Beyotime Biotechnology. Reagents for chemical reactions, including propargylamine, N,N'-diisopropylcarbodiimide (DIC), 1-hydroxy-7-azabenzotriazole (HOAt), Tris(3-hydroxypropyltriazolylmethyl)amine (THPTA), and 3-(2-Bromoethyl) indole were purchased from Innochem Technology. Hot Start Taq DNA polymerase was obtained from GenStar Biosolutions. Lambda exonuclease was purchased from Suzhou Novoprotein Scientific. Dulbecco’s Modified Eagle’s Medium (DMEM), fetal bovine serum (FBS), 10 μg/mL of streptomycin, 100 units/mL of penicillin, Phosphate-buffered saline (PBS), and 0.25% trypsin-EDTA were purchased from Thermo Fisher Scientific. Streptavidin-coated magnetic microbeads were obtained from Suzhou Beaverbio. Dynabeads™ MyOne™ Carboxylic Acid were purchased from Thermo Fisher Scientific. PE- Streptavidin conjugates were obtained from Sangon Biotech. DiO cell membrane green fluorescent dye and DiD cell membrane red fluorescent dye were purchased from Shanghai Yien Chemical Technology. Fast Silver Stain Kit, BCA Protein Concentration Determination Kit, ultra-sensitive TMB color developer, and TMB termination solution were purchased from Beyotime Biotechnology. All siRNAs were designed and synthesized by Sangon Biotech Co., Ltd., and all primers were biosynthesized by Tsingke Biotechnology.

The sources of antibodies and proteins used in this study are as follows: anti-TSG101 Monoclonal antibody (Proteintech, Cat. No.67381-1-Ig), anti-CD81 Monoclonal antibody (Proteintech, Cat. No. 66866-1-Ig), anti-CD63 Monoclonal antibody (Proteintech, Cat. No. 67605-1-Ig), anti-Calnexin Monoclonal antibody (Proteintech, Cat. No. 66903-1-Ig), anti-Beta Actin Monoclonal antibody (Proteintech, Cat. No.66009-1-Ig), anti-ASCC3 Polyclonal antibody (Proteintech, Cat. No. 17627-1-AP), anti-IMPDH1 Polyclonal antibody (Proteintech, Cat. No. 22092-1-AP), anti-GTSE1 Rabbit pAb (Abclonal, Cat. No. A13902), anti-ABCA2 Rabbit pAb (Proteintech, Cat. No. 20681-1-AP), anti-ALG3 antibody (Abmart, Cat. No. PS13522), anti-NF2/Merlin antibody (Abcam, Cat. No. ab109244), anti-PTPN3 antibody (Santa Cruz, Cat. No. sc-515181), anti-Moesin Polyclonal antibody (Proteintech, Cat. No.16495-1-AP), anti-MTCL1 antibody (Arigo, Cat. No. ARG44222), anti-SNX9 Polyclonal antibody (Proteintech, Cat. No. 15721-1-AP); IRDye 680RD Goat anti-Mouse IgG (Li-Cor, Cat. No. 926-68070); IRDye 800CW Goat anti-Rabbit IgG (Li-Cor, Cat. No. 926-32211). Human GTSE1 (Biodragon, Cat. No. BA09419), Human IMPDH1 (Biodragon, Cat. No. BA11364), Human ALG3 / NOT Protein (Tocris Bioscience, Cat. No. LS-G77737-20), Recombinant ATP Binding Cassette Transporter A2 (ABCA2) (Cloud-clone, Cat. No. RPD618HU01), Human ASCC3 (Biodragon, Cat. No. BA11095), Recombinant Protein Tyrosine Phosphatase, Non-Receptor Type 3 (PTPN3) (Cloud-clone, Cat. No. RPD586Hu01), Recombinant Human Merlin Protein (TargetMol, Cat. No. TMPH-01662), Human MSN (Biodragon, Cat. No. BA05360), Human MTCL1 (Biodragon, Cat. No. BA07545), Recombinant Sorting Nexin 9 (SNX9) (Cloud-clone, Cat. No. RPE879Hu01).

**Oligonucleotide synthesis and purification**

Oligonucleotides were synthesized in the laboratory using a standard phosphoramidite protocol on a DNA synthesizer (96 Channel DNA Synthesizer, Tsingke Biotechnology Co., Ltd., Beijing, China) at a 50 nmol scale. For library and aptamer synthesis, DMT-dC (Bz)-CE phosphoramidite, DMT-dG (iBu)-CE phosphoramidite, DMT-dA (Bz)-CE phosphoramidite, and DMT-5-Ethynyl-2’-deoxyuridine-3’-CE phosphoramidite were mixed in a vial, placed at the N vial position, and the random sequences were set on the synthesis programs to 25 N. The 3-(2-Azidoethyl)-1H-indole was then introduced to the alkyne-modified oligonucleotide sequences through a click reaction directly on the CPG solid-phase synthesis column.

40 μL of MeOH, 10 μL of 10× PBS, 25 μL of 500 mM 3-(2-Azidoethyl)-1H-indole in DMSO, and 25 μL of CuAAC catalyst solution (4 mM THPTA, 1 mM CuSO4, and 25 mM sodium ascorbate in ddH2O) were mixed and incubated with the CPG column for 3 hours at 37°C, 1500 rpm, three times. The solid-phase synthesis columns were washed with 200 μL of 50% water/MeOH. Deprotection was carried out using an ammonia resolver (Tsingke Biotechnology Co., Ltd.) at 95°C for 3 hours, followed by washing the CPG synthetic columns with 100% acetonitrile to remove short chains and desalinate. Purification was achieved by gel excision and nucleic acid purification columns (Oligo Clean & Concentrator Kits, RNA Clean & Concentrator Kits, Zymo Research). The full-length DNA strands were subsequently eluted with DEPC water, and the resulting DNA strand was freeze-dried and stored at -20°C.

**Synthesis and Purification of 3-(2-Azidoethyl)-1H-indole**

Weigh NaN_3_ (1.16 g, 18.5 mmol) and add it to a 250 mL round-bottom flask. Add 25 mL of DMSO and stir the mixture at room temperature for 15 minutes to fully dissolve. Then, add 3-(2-bromoethyl) indole (3.16 g, 14.6 mmol) to the mixture. Stir at room temperature overnight. After the reaction, slowly add 100 mL of ultrapure water to the reaction solution, which will be slightly exothermic. Transfer the reaction mixture to a separating funnel, add an equal volume of ethyl acetate, and extract three times. Retain the upper organic phase and collect all organic phases for spin steaming. Wash the concentrated organic phase three times with saturated salt water to remove excess NaN_3_ and collect the organic phase. Add anhydrous magnesium sulfate to the organic phase to remove water and remove the solid powder by suction filtration. Collect the filtrate. Use a rotary evaporator to remove the ethyl acetate solvent until the product turns into a brownish-yellow oily liquid. Store the product at -20°C. Identify the products by ^1^H NMR and ^13^C NMR.

**Cell culture**

HepG2 (human hepatocarcinoma, ATCC® HB-8065TM) cells were cultured in Dulbecco’s Modified Eagle’s Medium (DMEM, Gibco) supplemented with 10% fetal bovine serum (FBS, Gibco) in a humidified atmosphere containing 5% CO_2_ at 37°C (ThermoFisher). For sub-culturing cells from the monolayer, the cell culture medium was removed, and the cells were rinsed gently with PBS twice, followed by detachment with 0.05% trypsin EDTA at 37°C until the cells were rounded up and detached from the surface of the dish. Then, complete medium was added to the dish to inactivate the trypsin. Alternatively, cells can be digested and separated from the cell culture dish using enzyme-free cell dissociation buffer. The cells were centrifuged at 400×g for 5 minutes and collected at low temperature (4°C) before being resuspended with the appropriate buffer or medium.

**Preparation of human plasma**

Whole blood samples in the sodium heparin tube were obtained from the Department of Hepatobiliary and Pancreatic Surgery, Peking University Cancer Hospital, including 50 hepatocellular carcinoma patients and 30 healthy volunteers (Supplementary Table S1). This study was reviewed and approved by the Institutional Review Board and Ethics Committee of Peking University Cancer Hospital (2023KT129). Written informed consent for the use of clinical parameters and collected samples for further studies at the time of patient admission was obtained as a general standard procedure; the samples were anonymous and deidentified before use. The blood was then centrifuged at 2500×g for 10 minutes at 4°C (Eppendorf, 5910Ri). The plasma was transferred to a clean centrifuge tube, centrifuged again at 2500×g for 10 minutes at 4°C, then aliquoted and stored at -80°C until use.

**Clinical Specimen Collection and Patient Selection Criteria**

Patients were enrolled based on predefined inclusion and exclusion criteria. Inclusion criteria required a diagnosis of unresectable hepatocellular carcinoma (HCC) according to the Chinese Liver Cancer Staging (CNLC) system, classified as stage Ia-IIa or IIb-IIIb, with no prior systemic therapy. Eligible patients were aged 18-79 years and exhibited at least one measurable lesion or non-measurable but evaluable lesion. Additional inclusion criteria included a Child-Pugh score of 7 or less, an Eastern Cooperative Oncology Group (ECOG) performance status score of 0 or 1, and absence of severe cardiac, hepatic, or psychiatric comorbidities that would preclude surgical or medical intervention. Patients with active upper gastrointestinal bleeding within the preceding 12 months were excluded.

Exclusion criteria comprised a history of other malignant tumors within 3 years prior to HCC diagnosis and histopathological confirmation of non-HCC liver malignancies, including cholangiocarcinoma, mixed hepatocellular-cholangiocarcinoma, hepatosarcoma, or fibrolamellar hepatocellular carcinoma.

**Treatment Procedures and Follow-up**

Prior to treatment, all patients underwent standard imaging assessments, including contrast-enhanced computed tomography (CT) or magnetic resonance imaging (MRI). Routine laboratory tests were performed on peripheral blood samples, including complete blood count (CBC), liver and renal function tests, and tumor markers such as alpha-fetoprotein (AFP) and protein induced by vitamin K absence or antagonist-II (PIVKA-II). Diagnosis of HCC was established based on either clinical or pathological criteria in accordance with current guidelines. For patients with suspected extrahepatic metastases, positron emission tomography-computed tomography (PET-CT) was additionally conducted. In cases where gastroesophageal varices were suspected, upper gastrointestinal endoscopy was performed to assess the risk of bleeding. Following accurate tumor staging, a multidisciplinary team (MDT) discussion was conducted to determine the resectability of the liver tumor and to develop an individualized treatment plan.

Patients with unresectable HCC at baseline were enrolled in this study and received combination therapy consisting of immune checkpoint inhibitors (anti-PD-1/PD-L1 antibodies) and targeted agents. The targeted therapy options included either anti-VEGF antibody (e.g., bevacizumab) or tyrosine kinase inhibitors (e.g., lenvatinib, donafenib, or apatinib). Treatment continued until radiographic disease progression, unacceptable toxicity, or patient withdrawal due to personal reasons.

Follow-up evaluations, including imaging (CT or MRI) and tumor marker tests, were performed every 6–12 weeks. For patients who responded well to combination therapy and were assessed as having achieved resectability, surgical resection was subsequently performed. Postoperative patients continued immunotherapy for one year as per protocol requirements. Tumor response was evaluated using the modified Response Evaluation Criteria in Solid Tumors (mRECIST), classifying responses as complete response (CR), partial response (PR), stable disease (SD), or progressive disease (PD). PD was defined as non-responsive and CR, PR, and SD were defined as responsive. Study endpoints included objective response rate (ORR), disease control rate (DCR), recurrence-free survival (RFS, defined as the time from initial treatment to recurrence or death), and overall survival (OS, defined as the time from initial treatment to death).

**Isolation of Exosomes**

**Ultracentrifugation**

Isolation of Tumor Cell-Derived Exosomes: HepG2 cells were cultured in complete DMEM medium without exosomes, and the supernatant medium was collected after 48 hours of culture. The medium was collected by centrifugation at 4°C, 300×g for 10 minutes to remove cell debris, followed by 4°C, 2000×g for 10 minutes to remove dead cell debris, and 4°C, 10000×g for 30 minutes to remove the remaining debris. The supernatant was transferred to a clean 45Ti ultracentrifuge tube after passing through a 0.45 μm sterile filter, centrifuged at 4°C, 100,000×g in an ultracentrifuge (Beckman, XPN-100) for 3 hours, and the supernatant was discarded. The pellet was processed with sterile PBS and transferred to an 11.2 mL disposable ultracentrifuge tube, centrifuged at 4°C, 100,000×g for 3 hours, and the supernatant was discarded. Resuspend the extracellular vesicles at the bottom of the ultracentrifuge tube with 500 μL of sterile PBS and store in aliquots at −80°C, minimizing repeated freeze-thaw cycles.

Isolation of Plasma-Derived Exosomes: The pretreated plasma was thawed and transferred to a clean centrifuge tube, diluted 100-fold with sterile PBS, and centrifuged at 4°C, 16,000×g for 30 minutes to remove the remaining debris and large extracellular vesicles. The rest is identical to the isolation of cell-derived exosomes described above.

**Magnetic microbead immunoassay**

Thaw 1 mL of normal human plasma sample on ice, carefully extract the supernatant and transfer it to a clean 1.5 mL centrifuge tube. Centrifuge at 4°C and 16000 rpm for 10 minutes, discard the sediment, and transfer the supernatant to a clean 50 mL centrifuge tube. Add 8 mL of ultrapure water (filtered through a 0.22 μm sterile membrane), 10 mL of buffer EXA, 1 mL of buffer EXH, and 1 mL of pretreated plasma samples in turn. Mix by inverting 20 times, then centrifuge at 4°C and 10000 rpm for 10 minutes. Discard the sediment, and transfer the supernatant to a clean 50 mL centrifuge tube. Add 0.5 mL of vortex-dispersed and evenly mixed magnetic microbeads. Place the sample in a rotary mixer at 4°C, rotate at medium speed, and mix for 60-90 minutes. Transfer the sample to a 1.5 mL centrifuge tube, use magnetic separation, and let it stand at 4°C for 5 minutes, then discard the supernatant. Centrifuge at 4°C and 5000 rpm for 3 minutes, use magnetic separation, and further discard the residual liquid. Add 1 mL buffer EXE, vortex and mix for 30 seconds, and elute the extracellular vesicles. Use magnetic separation, let it stand at 4°C for 5 minutes, and centrifuge at 4°C and 10000 rpm for 3 minutes. After magnetic separation, transfer the supernatant to a new 1.5 mL centrifuge tube to obtain the exosomes.

**Protein-free ultracentrifugation**

Thaw 1 mL of normal human plasma sample on ice. Carefully extract the supernatant and transfer it to a clean 15 mL centrifuge tube. Add 8 mL of ultrapure water (filtered through a 0.22 μm sterile membrane) and 1 mL of buffer EXH. Mix well and observe for any white precipitation. If white precipitation occurs, centrifuge at 4°C and 10000 rpm for 10 minutes, then transfer the supernatant to a new 50 mL centrifuge tube. Dilute with sterile PBS and centrifuge at 4°C and 2000×g for 10 minutes to remove cell debris. After passing through a 0.22 μm sterile membrane, transfer the supernatant to a clean 70Ti ultracentrifuge tube. Centrifuge at 4°C and 100000×g for 3 hours, and discard the supernatant. Resuspend the pellet with sterile PBS and transfer it to an 11.2 mL disposable ultracentrifuge tube. Centrifuge again at 4°C and 100000×g for 3 hours, discard the supernatant, and resuspend the exosomes at the bottom of the ultracentrifuge tube with 500 μL of sterile PBS.

**Polymer precipitation**

Thaw 1 mL of normal human plasma sample on ice, carefully extract the supernatant, and transfer it to a clean 1.5 mL centrifuge tube. At room temperature, centrifuge at 10000×g for 20 minutes to remove cell debris. Transfer the pretreated plasma to a new 1.5 mL centrifuge tube, add 0.5 times the volume of PBS, and vortex mix the sample. Then add 0.05 times the volume of protease K, and vortex the sample, incubating it at 37°C for 10 minutes. Add 0.2 volumes of exosome precipitation reagent (from the total exosome isolation kit) to the sample, vortex and mix well until it becomes turbid. Incubate the samples at 4°C for 30 minutes. After incubation, centrifuge the samples at room temperature for 5 minutes at 10000×g, and discard the supernatant. Then, centrifuge again at room temperature at 10000×g for 30 seconds and discard the supernatant. Resuspend the precipitated exosomes with 500 μL of sterile PBS.

**SEC for isolation of plasma-derived exosomes**

Thaw 0.5 mL of normal human plasma samples on ice, carefully aspirate the supernatant and transfer it to a clean 50 mL centrifuge tube, and add a 1× 100-fold dilution of D-PBS to dilute the plasma diluent. The plasma dilution was mixed well, centrifuged at 4°C, 2000×g for 10 minutes, the supernatant was removed, and the sample passed through a 0.22 μm sterile filter membrane. At 4°C, 5000×g was centrifuged, and the obtained filtrate was concentrated to within 500 μL with a 100 kDa molecular weight cutoff (MWCO) ultrafiltration tube. Transfer the concentrate to a clean 1.5 mL centrifuge tube, centrifuge at 4°C, 10,000×g for 1 hour, and add 5 mL of D-PBS to wash the size-exclusion chromatography (SEC) column for later use. After centrifugation, transfer the samples to a clean 1.5 mL centrifuge tube and add D-PBS to keep the volume at 500 μL. Add 500 μL of concentrate to the SEC column; once the sample is fully in the frit, add D-PBS, start collecting components, label one component per 500 μL effluent, and store the collected components at -80°C for subsequent analysis or downstream applications.

**Transmission electron microscope (TEM)**

The morphology of the exosomes was characterized by transmission electron microscopy (TEM). In brief, exosomes were dispersed and deposited onto an ultrathin copper mesh for 2 minutes. Then, the exosomes were fixed with a 2% phosphotungstic acid negative dye solution for 1 minute to absorb the excess solution, and the process was repeated for several rounds. Finally, the prepared samples were completely dried at room temperature. Images were captured using the JEM1400 TEM system (JEOL, Japan).

**Nanoparticle Tracking Analysis (NTA)**

Exosome samples were diluted to a final concentration of 1×10^7^–10^9^ particles/mL using PBS and examined using a NanoSight NS300 (NanoSight Ltd., UK). Automatic settings were selected for minimum expected particle size of 30 nm, minimum track length of 10 frames, detection threshold of 5, and constant temperature maintenance at 25°C. For each sample, three independent 60-second videos were captured under controlled fluid flow conditions, with subsequent analysis generating quantitative measurements of EV size distribution (nm) and concentration (particles/mL).

**Pre-enrichment of aptamer**

**Library preparation**

Aptamers were pre-enriched using the cell-SELEX procedure. The selection began with a single-stranded DNA library (Integrated DNA Technologies) consisting of a randomized region of 25 nucleotides (N25) flanked by two constant primer-hybridization sites (5’- CACGACGCAAGGGACCACAGG-𝑁25-CAGCACGACACCGCAGAGGCA-3’; N=A:EdU:G:C). Before each round of selection and binding experiments, the DNA library and aptamer pools were denatured by heating for 5 minutes at 95°C and then renatured on ice for 10 minutes. Information on relevant library sequences and primer sequences can be found in Supplementary Table S2.

**Synthesis of exosomes latex microbead complex**

Exosomes are cross-linked on the surface of aldehyde-ylated latex microbeads (A37304, Thermo Fisher Scientific), and the exosome-latex microbead complex is recovered by centrifugation. Exosomes (8×10^9^ particles) were incubated with 4 μL of latex microbeads for 15 minutes at room temperature, then PBS was added in a final volume of 200 μL for 1.5 hours at room temperature. Add 100 μL of Blocking Buffer (containing 1 M glycine, 20% BSA (w/v), 1× D-PBS, pH 7.4) and incubate for 30 minutes at room temperature. Centrifuge at 10,000×g at 4°C for 2 minutes, then wash twice with 200 μL PBS, and finally resuspend in 50 μL PD Binding Buffer (containing 2.5mM MgCl_2_, 1 mg/mL BSA, 0.1mg/mL Salmon sperm DNA, 0.1% Tween 20, 1× D-PBS, pH 7.4) and store for later use.

**Library incubation and preconcentration completed**

4 μL of empty latex microbeads of unmodified exosomes were added to 1 mL of Prebinding buffer (containing 2.5mM MgCl_2_, 1 mg/mL BSA, 0.1mg/mL Salmon sperm DNA, 0.1% Tween 20, 1 × D-PBS, pH 7.4) for 1 wash. Then, 1 nmol of the initial library was added, Prebinding buffer was supplemented to a total volume of 500 μL, incubated at 1500 rpm, 37°C for 1 hour, centrifuged at room temperature at 10,000 × g for 2 minutes, and the supernatant was collected. The supernatant was mixed with 50 μL of exosome-latex microbead complexes in Prebinding buffer, and 1.6 nmol of competing library was added to prevent non-specific action of extracellular vesicles and initial screening stock in a total volume of 500 μL and incubated at 1500 rpm at 37°C for 1 hour. Unbound DNA sequences were washed away with the Prebinding buffer 3 times. Then, the remaining DNA sequences bound to the microbeads were eluted in 95°C water 3 times. The eluted sequences were digested by λ exonuclease to separate the selected ssDNA from the double-stranded DNA. The resulting ssDNA was indole modified, further purified, and served as a new library for the second round of selection. After 2 rounds of pre-enrichment, the new ssDNA library was ready for the AptEx-ID.

**Microbead-displayed aptamer screening strategy targeting HCC-specific exosomes**

**Preparation of alkyne-based magnetic microbeads**

To couple forward primers (FP) to 100 µL 1 µm MyOne carboxylic acid magnetic microbeads (ThermoFisher, 65012), the microbeads were washed with 100 µL DMF 3 times. Then, 50 µL of 2M propargylamine in DMF, 50 µL of 2.8M DIC in DMF, and 100 µL of 1M HOAt in DMF were added to the microbeads. The mixture was shaken for 2.5 hours at 50°C at 1500 rpm. Afterward, the microbeads were washed with DMF 3 times and resuspended in DMF. Subsequently, 80 µL of the microbeads were washed with 1% Tween 20.

The microbeads were incubated in CuAAC reaction mixture (containing 1.0 mM CuSO4, 1.2 mM THPTA, 6 mM ascorbic acid, 0.05% Tween 20, 20 µM 5'-N3-PEG18-modified FP, 5% DMSO, 220 mM TEAA, pH 7.4, 1× D-PBS, and DEPC water with a total volume of 100 µL) for 3 hours at 37°C and 1500 rpm. Then, the FP-microbeads were washed with TE buffer (containing 10 mM Tris, 1 mM EDTA, pH 8.0) twice and resuspended in 100 µL TE buffer. To block the excess uncoupled alkynyl epitopes on the surface of the microbeads, 100 mM N3-PEG18 was added instead of 20 µM 5'-N3-PEG18-modified FP according to the conjugation system described above, so that the conjugated microbeads were completely blocked.

**Emulsion PCR**

For the emulsion PCR, the oil phase consisted of 7% KF-6038 and 93% DMF-A-6CS and was spun for 2 hours to mix well. The aqueous phase contained 6 µL of dNTPs (2.5 mM of each nucleotide: dATP, dGTP, dCTP, dEduTP), 3 µL of 100 µM reverse primer (RP), 1.5 µL of Hot Start Taq, 15 µL of 5× Hot Start Taq Buffer, 8 pM template DNA, 12.5 µL of 60 mM PEG-8000, and 5 µL of the FP-microbeads. Water was added to reach a total volume of 75 µL. The O/W emulsions were created by adding 300 µL of the oil phase to 75 µL of the aqueous phase and emulsifying the mixture in a grinder at 40 Hz for 50 seconds. The emulsion was then distributed to PCR tubes for 50 µL aliquots per tube. PCR was performed with the following cycling conditions: 95 °C for 3 minutes, followed by 30 cycles of 95 °C for 15 seconds, 60 °C for 15 seconds.

**Break emulsion**

After PCR, the emulsion was collected into a 2 mL PCR tube and vortexed for 30 seconds. The particles were collected after 2 minutes of centrifugation at 5000 rpm, then the clear oil phase was carefully removed. Next, 1 mL of the emulsion breaking buffer (containing 100 mM NaCl, 1% Triton X-100, 1% SDS, 10 mM Tris-HCl, and 1 mM EDTA, pH 7.5) was added to the particles. After vortexing for 30 seconds and centrifugation for 90 seconds at 15,000×g, the tube was placed in magnetic separation, and the buffer was carefully discarded. This step was repeated 2 times, followed by washing the particles with TE buffer 3 times using magnetic separation.

To generate ssDNA, the particles were incubated in 200 µL of 100 mM NaOH at 50 °C for 2 minutes, and the supernatant was carefully removed using magnetic separation. This step was repeated 2 times, and the particles were washed with TE buffer 3 times using magnetic separation.

**Emulsion PCR efficiency verification**

After PCR, the 0.5 µL of Alexa Fluor 633 modified RP (10 µM) was incubated with 0.5 µL of the emulsion containing magnetic microbeads in 19 µL of STE buffer (10 mM Tris, 1 mM EDTA, 50 mM NaCl, pH 7.4) in a metal bath at 55°C, 1500 rpm, for 10 minutes. At the end of the incubation, the mixture was quickly cooled on ice for 2 minutes. The efficiency of emulsion PCR was determined by adding 100 µL of TE buffer for 2 washes, followed by magnetic separation and resuspension in 100 µL of TE buffer. Detection was performed with a flow cytometer. According to the Poisson distribution, when less than 35% of the microbeads contain PCR products, most microbeads are monoclonal.

**Library indole modification**

The library microbeads were washed with TE buffer three times using magnetic separation. Firstly, 1.2 μL of 100 mM THPTA solution, 1 μL of 100 mM CuSO_4_ solution, and 6 μL of 100 mM sodium ascorbate solution were added to prepare the CuAAC catalytic system and placed at room temperature for 10 minutes. Then, 10 μL of 1% Tween-20 solution, 10 μL of 2.2 M triethylamine acetic acid (TEAA) solution, 20 μL of DMSO, 10 μL of 100 mM azide indole solution, 10 μL of 10×D-PBS, and 31.8 μL of ddH_2_O were added and the mixed system was placed in a metal bath at 37°C at 1500 rpm and incubated for 3 hours. After incubation, magnetic separation was performed. The particles were washed with 100 μL TE buffer three times using magnetic separation. Finally, the particles were resuspended in 200 μL TE buffer and stored at 4°C.

**Labeling of exosomes**

Tumor cell-derived exosomes were selected for screening, while plasma-derived exosomes from healthy humans were used for the negative selection step. Tumor-derived exosomes and healthy human plasma-derived exosomes were incubated with DiO (1 μM) and DiD (1 μM) for 30 minutes at 1500 rpm at room temperature, respectively. To remove the free dye, transfer the exosomes labeled with the fluorescent dye to a clean ultracentrifuge tube and discard the supernatant. The resulting exosomes were thoroughly washed in sterile PBS and centrifuged again at 4 °C, 100,000×g for 90 minutes. The resulting labeled exosomes were resuspended in 100 μL of sterile PBS and stored at 4°C.

**Exosomes magnetic microbead incubation**

Indole-modified library magnetic microbeads and DiO-labeled tumor-derived exosomes are incubated together, while DiD-labeled healthy human plasma-derived exosomes are used as competitive molecules. Concurrently, 100 pmol of competitive DNA is added to prevent non-specific interactions between the library magnetic microbeads and exosome surface proteins in each round of screening. The PD binding buffer is then supplemented to 500 μL and the mixture is incubated at 37°C in a metal bath at 1500 rpm for 1 hour. Subsequently, 500 μL of PD washing buffer (containing 2.5 mM MgCl_2_, 0.1% Tween 20, 1×D-PBS, pH 7.4) is added for two washes. The mixture is then incubated at 50°C in a metal bath at 1500 rpm for 2 minutes.

**Flow sorting**

Resuspend in 2 mL of sorting buffer (containing 2.5 mM MgCl2, 1×D-PBS, pH 7.4) to ensure that the flow cytometer separates the first 0.1% of the fluorescence intensity of the magnetic microbead particles under the condition of protection from light for subsequent amplification. After the collection of green fluorescence-labeled particles, we amplified the sequences through PCR to create an enriched pool for the subsequent round of aptamer particle synthesis. In order to further improve the affinity and specificity of the library in multiple rounds of PD screening, the screening pressure was increased round by round by reducing the number of library microbeads, increasing the negative sieve molecules round by round, and adding competing libraries.

**ssDNA recovery**

After the screening procedure, primers with barcode sequences (Supplementary Table S3) were used to amplify samples from pre-enrichment to three rounds of aptamer selection. A total of 250 pmol dsDNA was used for 10 cycles of PCR. The aqueous phase contained 0.8 µL of dNTPs (2.5 mM of each nucleotide: dATP, dGTP, dCTP, dTTP), 0.5 µL of 10 µM forward primer (FP), 0.5 µL of 10 µM reverse primer (RP), 0.2 µL of Hot Start Taq, and 0.2 µL of 5× Hot Start Taq Buffer. Water was added to reach a total volume of 10 µL. PCR products were recovered using the QIAquick® Gel Extraction Kit (28706, QIAGEN) and analyzed using the Illumina NovaSeq Sequencing System.

**Sequences analysis**

Data processing and analysis were performed using Aptasuite. Initially, the software tallied the number of replicates for each sequence and computed the edit distance between all sequences, which was then stored using locality-sensitive hashing. Subsequently, the sequences were clustered into families based on an edit distance threshold of 5. These clusters were further organized by sorting them according to the abundance of their members. Clusters that enriched during the screening process and ranked highest in the last round were selected, with the highest sequence in each cluster picked (Supplementary Table S4).

**Flow cytometry analysis**

**Flow cytometric verification of nucleic acid magnetic microbeads-exosomes**

After selecting the appropriate verification sequence through data analysis, the sequence used to prepare nucleic acid magnetic microbeads was obtained from Tsingke Biotechnology. According to the method shown in the screening process, the synthesized sequence was used as a template, the nucleic acid magnetic microbeads were prepared to verify the PCR system, and the PCR reaction cycle was 30 rounds. The PCR reaction products were single-stranded, and the PCR efficiency was verified. If the efficiency was within a reasonable range, the nucleic acid magnetic microbeads were modified with indole. Tumor-derived exosomes (DiO) and healthy human plasma-derived exosomes (DiD) were fluorescently labeled, respectively. The labeled extracellular vesicles (2 × 108 particles) were incubated with 1µL nucleic acid magnetic microbeads, supplemented to 50µL with PD binding buffer, and incubated at 37°C in a metal bath at 1500 rpm for 45 minutes. Add 200µL PD washing buffer for washing twice, and re-suspend it in 100µL PD washing buffer. Then, the resuspended samples were evaluated using flow cytometry (Cytoflex, Beckman). The initial library was used as the negative control. The fluorescence intensity was analyzed using FlowJo (v10.8) software.

**Flow cytometric validation of latex microbeads-exosomes**

After selecting the appropriate verification sequence through data analysis, the Cy3-labeled DNA sequence was synthesized using a DNA synthesis instrument, and the corresponding nucleic acid sequence was obtained by indole modification and purification. First, tumor-derived exosomes latex microbead complexes and normal human plasma-derived exosomes latex microbead complexes were prepared. The two types of exosomes were mixed with the latex microbeads (at a ratio of 8 × 10^8^ particles/0.5 µL latex microbeads) and incubated at 1000 rpm for 15 minutes. Then, sterile PBS was added to 1 mL and incubated at 1000 rpm for 2 hours. The mixture was centrifuged at 12000 rpm for 2 minutes and the supernatant was discarded. After adding a certain amount of PD binding buffer for washing once, the latex microbeads were re-suspended, and each 0.5 µL of the latex microbead compound was evenly distributed into the centrifuge tubes to be tested. Each Cy3-labeled snap cool aptamer (500 nM) was added to the system, bringing the total volume to 50 µL, and incubated at 1500 rpm at room temperature for 30 minutes. The mixture was centrifuged at 12000 rpm for 2 minutes and the supernatant was discarded. After washing twice with 200 µL of PD washing buffer, the latex microbeads were finally re-suspended in 100 µL of PD washing buffer. Then, the resuspended samples were evaluated using flow cytometry (Cytoflex, Beckman). The initial library was used as the negative control. The fluorescence intensity was analyzed using FlowJo (v10.8) software.

**Cell flow cytometry：**

To analyze the binding ability of DNAs, after being collected by a non-enzyme cell detachment solution (Leagene Biotech, China), 105-106 HepG2 cells were incubated with biotin-labeled or Cy3-labeled ssDNA (250 nM final concentration) in cell binding buffer (containing 4.5 mg/mL glucose, 2.5 mM MgCl_2_, 1 mg/mL BSA, 0.1 mg/mL Salmon sperm DNA, 1×D-PBS, pH 7.4) at 4 °C for 30 min. The cell suspension was centrifuged and washed twice with cell washing buffer (containing 4.5 mg/mL glucose, 2.5 mM MgCl_2_, 1×D-PBS, pH 7.4). Then cells were incubated with 1 µL PE-Streptavidin conjugates in cell binding buffer at 4 °C for 10 min in the dark. If the DNA is Cy3-labeled, the previous step can be skipped. The cell suspension was centrifuged and washed twice with cell washing buffer. Then, the resuspended samples were evaluated using flow cytometry (Cytoflex, Beckman). The initial library was used as the negative control. The fluorescence intensity was analyzed using FlowJo (v10.8) software.

**Apta-ELISA procedures**

**Apta-ELISA detection:**

Dilute the target extracellular vesicles in 0.05M carbonate bicarbonate buffer. Add 100 µL (containing 10^7^ particles) of the target extracellular vesicles per well to a high adsorption well plate, and incubate overnight at 4°C (approximately 10 hours) to ensure that the extracellular vesicles are fully adsorbed onto the well plate. After incubation, wash three times with 0.05% PBS-T solution, allowing each wash to stand for three minutes. Shake dry. Add 100 µL of ELISA blocking solution and block at room temperature for one hour to reduce non-specific binding. Wash three times with 0.05% PBS-T solution, allowing each wash to stand for three minutes. Shake dry. Add 250 nM biotin-modified corresponding aptamer (snap cool). Add Binding buffer ELISA to a total system volume of 50 µL and incubate at room temperature for one hour. Wash three times with 0.05% PBS-T solution, allowing each wash to stand for three minutes. Shake dry. Add 100 µL of freshly prepared 0.1 µg/mL SA-HRP working solution and incubate in the dark for one hour. Wash four times with 0.05% PBS-T solution, allowing each wash to stand for three minutes. Shake dry. Add 100 µL of TMB chromogenic solution and incubate in the dark for approximately 15 minutes until a blue color forms. Add 100 µL of termination solution to stop the reaction, changing the color from blue to yellow. Use a multifunctional microplate reader (BioTek Synergy Neo2, BioTek) to detect and record data at 450 nm (OD_450_).

**Inclusion testing of tumor derived extracellular vesicles:**

According to the preliminary established method, three types of incorporation tests were conducted: direct gradient incorporation of tumor-derived exosomes into 0.05M carbonate bicarbonate buffer, direct gradient incorporation of tumor-derived exosomes into plasma samples and ultracentrifugation recovery, or direct gradient incorporation of tumor-derived exosomes into plasma dilutions depleted of exosomes. The gradients were set to 0, 5×10^6^, 10^7^, 2×10^7^, 5×10^7^, and 8×10^7^ particles per well. 250 nM biotin-modified aptamer HYC11 (snap cool) was added, and three wells were set up. TMB color reagent was used for color development, and a multifunctional microplate detector was used at 450 nm to measure (OD_450_), record, and analyze data.

**Exploration of color development methods:**

According to the preliminary established method, tumor-derived exosomes were directly incorporated into normal human plasma-derived exosomes, with the total count of extracellular vesicles maintained at 1×10^8^ particles per well. To the tumor extracellular vesicles, 250 nM biotin-modified aptamer (snap cool) was added, and three replicate wells were established. Various color development methods were chosen, specifically TMB color development (with gradients of 0%, 0.1%, 1%, 10%, 20%, 50%, 80%, and 90%) and fluorescent color development (with gradients of 0%, 0.1%, 1%, 2%, 5%, 10%, 25%, 50%, 80%, and 100%). Measurements were taken at 450 nm (OD_450_) using a multifunctional microplate detector, and the data were subsequently recorded and analyzed.

**Exploration of internal standard reference materials:**

Thrombin standard was chosen as the reference material for the component to be tested and quantitatively incorporated into the sample. The component and the thrombin standard were detected using the corresponding aptamer probes, yielding respective measurement data (OD_450_). This allowed us to calculate the ratio of OD_450_ between the exosomes test group and the thrombin group. To maintain the OD_450_ of the internal reference detection within an appropriate range, we explored both the concentration of the internal reference and its OD450 value. We selected thrombin and its corresponding biotin-modified aptamer TBA29 (5’-AGTCCGTGGTAGGGCAGGTTGGGGTGACT-3’), established a concentration gradient, and measured at 450 nm (OD450) using a multifunctional microplate detector, following the established aptamer-based enzyme-linked immunosorbent assay protocol. The data were recorded and analyzed to determine the optimal concentration of the internal reference.

**Optimization of APTA ELISA detection method:**

The target exosomes and thrombin standard were mixed and diluted in 0.05M carbonate-bicarbonate buffer. 100 µL of the mixture, containing target exosomes (1×10^7^ particles) and 250 ng/µL of thrombin standard, was added to a high-adsorption plate, divided into two, and incubated at 4°C overnight to ensure thorough adsorption of the exosomes and thrombin mixture on the plate. The remaining steps were identical to those before optimization. The OD_450_ values of the target extracellular vesicle group and the thrombin group were recorded, and the relative OD_450_ ratio was calculated. To compare the ability of aptamers to distinguish tumor exosomes from normal human plasma-derived exosomes, the established method was used to verify the specificity of aptamers. Then the clinical samples were detected by the same method.

**Target identification of Aptamer**

HepG2 cells (1×10^8^ cells) were detached using a non-enzymatic cell detachment solution, washed twice, and then mixed with 2 mL of pre-cooled hypotonic buffer (50 mM Tris-HCl, pH 7.4, supplemented with 10 mM PMSF and 1× protease inhibitor cocktail before use) on ice for 30 minutes. Cytosolic and nuclear proteins were removed by centrifugation, and the cell debris containing the membrane proteins was further treated with 500 µL of lysis buffer (2% Triton X-100 and 1% Tween-20 in hypotonic buffer) for 30 minutes. Subsequently, the mixture was centrifuged at 4000×g for 10 minutes, and the supernatant containing the membrane proteins was collected. The resulting supernatant was blocked with 0.1 mg/mL BSA and 1 mg/mL tRNA. Then, the membrane protein solution was incubated with 250 nM biotin-aptamer and biotin-library for 20 minutes, respectively. Streptavidin-coated magnetic microbeads were then added to pull down the DNA-protein complex for 1 hour. After washing three times, the magnetic microbeads were mixed with SDS loading buffer and denatured at room temperature for 30 minutes. The proteins were separated using a 10% SDS-PAGE gel. Subsequently, the gel was visualized using silver staining. The aptamer-purified protein bands were excised for digestion and analyzed by LC-MS (Orbitrap Velos Pro, Thermo Fisher).

**Proteomic analysis based on mass spectrometry**

Quantitative proteomics analysis was performed using liquid chromatography-mass spectrometry (LC-MS) to identify differential proteins and determine their corresponding target information. Protein information was retrieved from the UniProt human protein database for data processing. Based on the quantitative mass spectrometry data, the average intensity of the control group and the experimental group was calculated, and the ratio was obtained. The ratio value, expressed as log2 (ratio), was used as the X-axis, and the p-value, expressed as -log10 (p-value), was used as the Y-axis. Abnormal values or null values were not included in subsequent data analysis. A differential protein analysis volcano plot was generated using the criteria of log2 (ratio) > 3.32 and p-value > 2.99.

**RNA interference experiments**

All siRNAs were synthesized by Sangon Biotech Co., Ltd., and the information can be found in Supplementary Table S5. HepG2 cells were seeded on 6-well plates at a concentration of 2×10^5^ cells/mL 24 hours before transfection and cultured in 2 mL DMEM complete medium. The next day, when the degree of cell fusion reached 60%-80%, 1.5 mL DMEM double-antibody-free medium was added, gently mixed, and incubated at room temperature for 15 minutes. 500 µL of the incubation mixture was then added evenly to each well, and the cells were cultured without changing the medium. Both the NC group and siRNA group were transfected simultaneously and cultured for 48-72 hours. Subsequently, the cells were harvested and analyzed by flow cytometry, and protein extracts were detected by Western blot.

**Quantitative reverse transcription-PCR (qPCR)**

Molpure Cell/Tissue Total RNA Kit (Yeasen, China) was used to extract both cell and tissue RNA according to the manufacturer's protocol. Complementary DNA (cDNA) was generated utilizing the Hifair AdvanceFast One-Step RT-gDNA Digestion SuperMix for qPCR Kit (Yeasen, China). Furthermore, the Hieff® qPCR SYBR® Green Master Mix Kit (Yeasen, China) was utilized to perform qRT-PCR. The reaction conditions were 95°C for 5 minutes, followed by 40 cycles of 95°C for 10 seconds, 55°C for 20 seconds, and 72°C for 20 seconds, with a final step at 95°C for 15 seconds. The relative expression of the target gene was assessed using the 2^ΔΔCt^ method, followed by normalization against ACTB. The primers are listed in Supplementary Table S6.

**Protein extraction and Western blotting**

HepG2 cells or exosomes were washed with cold phosphate-buffered saline (PBS) and subsequently lysed with RIPA buffer supplemented with protease inhibitors. Protein concentration was measured using a BCA Protein Concentration Determination Kit (Beyotime Biotechnology, China) according to the manufacturer's instructions. Equal amounts of protein (30 µg) were loaded onto SDS-PAGE gels and transferred to polyvinylidene fluoride (PVDF) membranes. After blocking with 5% non-fat milk for 60 minutes, the membranes were incubated with primary antibodies against the target proteins and β-actin overnight at 4°C. The following day, the membranes were washed three times with TBST (2% Tween) for 7 minutes each, followed by incubation with the corresponding secondary antibodies conjugated to fluorescent dyes for 60 minutes at room temperature. The membranes were then washed twice with TBST (2% Tween) for 7 minutes each and once with TBS for 7 minutes. Protein expression was detected using the Odyssey Infrared Imaging System (Odyssey® CLX, Li-Cor), and the intensity of the bands was quantified using ImageJ software.

**Biolayer interferometry (BLI)**

The binding affinity of aptamers to target recombinant protein was assessed by biolayer interferometry. The binding experiments were performed in black 96-well plates with 200 µL per well of BLI binding buffer (containing 2.5 mM MgCl_2_, 0.1% Tween 20, 1×D-PBS, pH 7.4). The SA sensors were first prehydrated with BLI buffer for a minimum of 10 minutes. Biotin-labeled aptamers were immobilized on SA sensors until a signal shift of 1-2 nm was achieved. The binding affinity of each aptamer to different concentrations of target recombinant protein (6.25 nM, 12.5 nM, 25 nM, 50 nM, 100 nM, 200 nM, 400 nM) was then determined using the following protocol: association (120 s), dissociation (120 s), and regeneration (5 s, repeated 4 times). The dissociation constant (Kd) for each aptamer was calculated using the analysis software provided with the BLI system (Octet Red96e, Sartorius).

**Structure prediction**

**Molecular simulation of aptamers**

For the molecular simulation of aptamers, the 3dRNA/DNA web server was used to build the 3D structure of aptamers. The simulation was performed with the AMBER 16 molecular simulation package. To obtain molecular mechanical parameters for Indole-EdU, ab initio quantum chemical methods were employed using the Gaussian 16 program. The geometry was fully optimized, and then the electrostatic potentials around them were determined at the B3LYP/6-31G* level of theory. The RESP strategy was used to obtain the partial atomic charges. The starting structure of aptamers was solvated in TIP3P water using an octahedral box, which extended 8 Å away from any solute atom. To neutralize the negative charges of simulated molecules, Na+ counter-ions were placed next to each negative group. Molecular dynamics (MD) simulation was carried out using the PMEMD module of AMBER 16. The calculations began with 500 steps of steepest descent followed by 500 steps of conjugate gradient minimization with a large constraint of 500 kcal mol-1 Å-2 on the atoms of the aptamers. Then 1000 steps of steepest descent followed by 4000 steps of conjugate gradient minimization with no restraint on the complex atoms were performed. Subsequently, after 200 ps of MD, during which the temperature was slowly raised from 0 to 300 K with a weak (5 kcal mol-1Å-2) restraint on the aptamers, the final unrestrained production simulations of 100.0 ns were carried out at constant pressure (1 atm) and temperature (300 K). Throughout the entire simulation, SHAKE was applied to all hydrogen atoms. Periodic boundary conditions with minimum image conventions were applied to calculate the non-bonded interactions. A cutoff of 10 Å was used for the Lennard–Jones interactions. The final conformations of the complexes used for discussion were produced from the 1000 steps of minimized averaged structure of the last 20.0 ns of MD.

**Acquisition of protein structure:**

The protein structure can be obtained from the PDB database or UniProt database, and the corresponding single crystal analysis structure file can be downloaded from the respective link. Alternatively, unresolved protein structure information can be obtained through AlphaFold simulation.

**Complex structure prediction:**

The structure predictions of protein and aptamer complexes were performed on the following two web servers: https://alphafoldserver.com/ and http://hdock.phys.hust.edu.cn/. In the HDOCK prediction results, models with highly ranked docking scores, high confidence scores, low ligand RMSD, and good experimental fit were considered.

**IndU substitution and truncation analysis**

Biotin-labeled IndU substitution and truncation sequences were synthesized according to the sequences in Supplementary Table S7. HepG2 cells were dissociated with a non-enzyme cell detachment solution and then incubated with Cy3-labeled DNA for flow cytometric analysis.

**Differential gene expression analysis**

Differential gene expression analysis was performed on the following two websites: http://gepia2.cancer-pku.cn/#index and https://cistrome.shinyapps.io/timer/. The correlation between biomarkers and overall survival (OS), disease-free survival (DFS), and tumor stage was assessed using the GEPIA2 database (https://gepia2.cancer-pku.cn/). The hazard ratio (HR) was calculated as the ratio of the risk in the high biomarker expression group to that in the low expression group, serving as a measure to assess the influence of biomarker expression on patient survival.

**Machine learning**

We analyzed data from 80 subjects (50 HCC patients, 30 non-cancer controls) with measurements of 31 biomarkers. Given the moderate sample size, we implemented specific strategies to ensure model reliability and minimize overfitting. We employed a stratified sampling strategy to maintain the class distribution across partitioned datasets. The cohort was divided into training (80%, n=64; 40 HCC, 24 controls) and testing (20%, n=16; 10 HCC, 6 controls) sets. For hyperparameter tuning, we implemented 8-fold stratified cross-validation on the training set, ensuring each fold maintained the original class distribution. This approach preserved the test set exclusively for final model evaluation while utilizing cross-validation to prevent overfitting during model development.

For feature selection, we implemented a multi-stage framework. First, statistical filtering using Welch's t-tests reduced the initial feature set by identifying significantly differentially expressed biomarkers. Subsequently, Logistic Regression-based Recursive Feature Elimination with Cross-Validation (RFECV) further refined this set to an optimal subset of features with maximal predictive power. We validated our feature selection approach by comparing results with SelectKBest using ANOVA F-values, which independently identified overlapping high-scoring biomarkers, confirming the robustness of our signature. Multiple classification algorithms were evaluated, including Naive Bayes, K-Nearest Neighbor, Support Vector Machines, Logistic Regression, Decision Trees, Random Forest, and XGBoost. Logistic Regression was selected as our final model based on superior performance, interpretability, and suitability for moderate sample sizes. The model incorporated L1 regularization to perform automatic feature selection during training and mitigate overfitting.

Hyperparameter optimization was conducted via grid search with 8-fold cross-validation across regularization strength, penalty type, solver algorithm, and L1 ratio parameters. ROC-AUC served as the optimization metric. The final model was evaluated on the independent test set using standard performance metrics (accuracy, precision, sensitivity, specificity, F1-score, ROC-AUC). Although our dataset included treatment effectiveness data, the machine learning approach focused exclusively on diagnostic model development due to sample size constraints, as reliable prognostic modeling would require a substantially larger cohort with longitudinal follow-up data.

**Supplementary Figures**

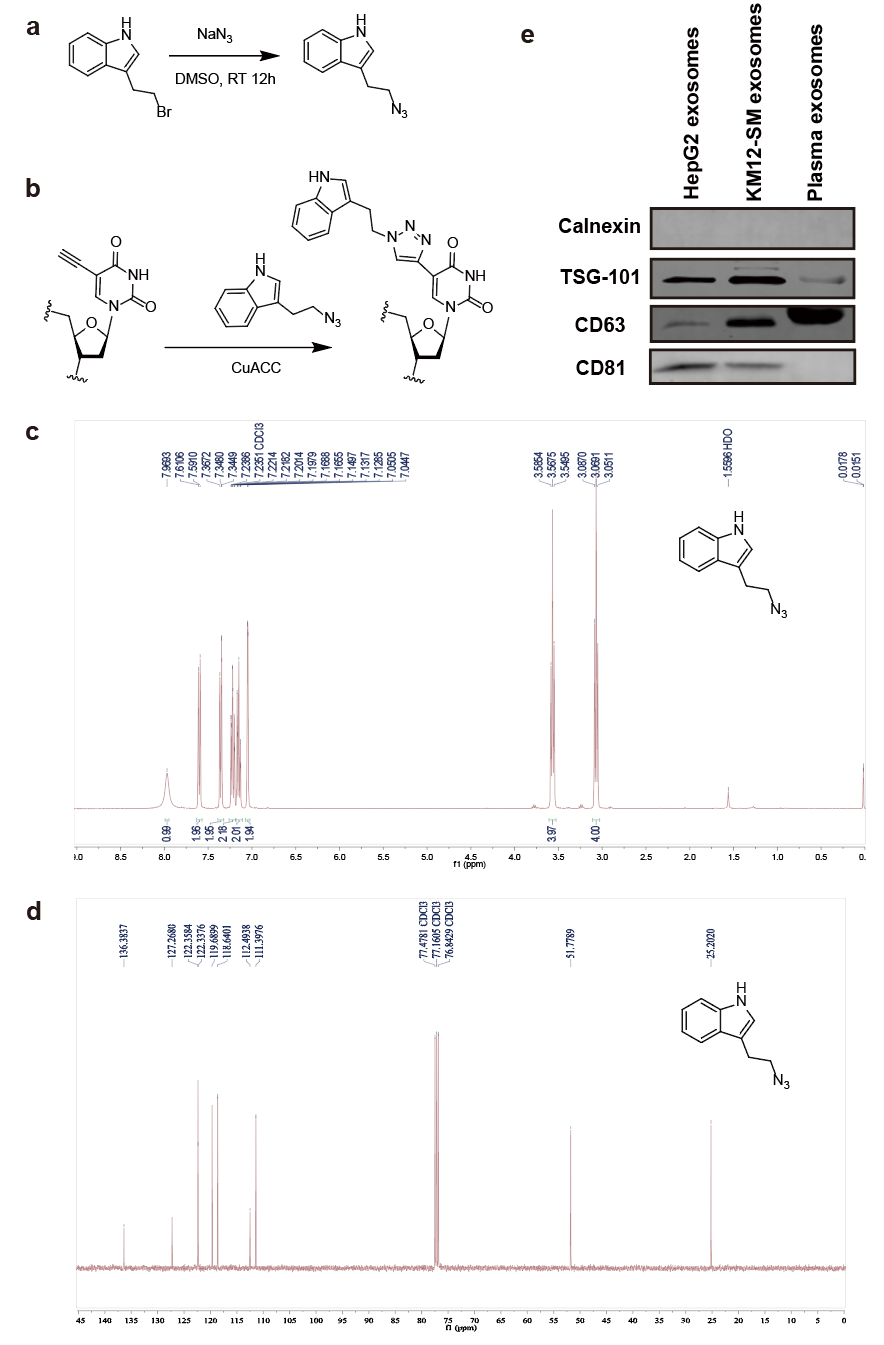

**Supplementary Fig. S1. Synthesis of 3-(2-Azidoethyl)-1H-indole and indole-EdU. a** Schematic diagram of the chemical synthesis of 3-(2-Azidoethyl)-1H-indole. **b** Identification of 3-(2-Azidoethyl)-1H-indole compound by ^1^H NMR. **c** Identification of 3-(2-Azidoethyl)-1H-indole compound by ^13^C NMR. **d** The molecular formula of the click reaction to convert EdU into IndU. **e** Characterization of HepG2 and healthy volunteer plasma-derived exosomes (Western blot image).

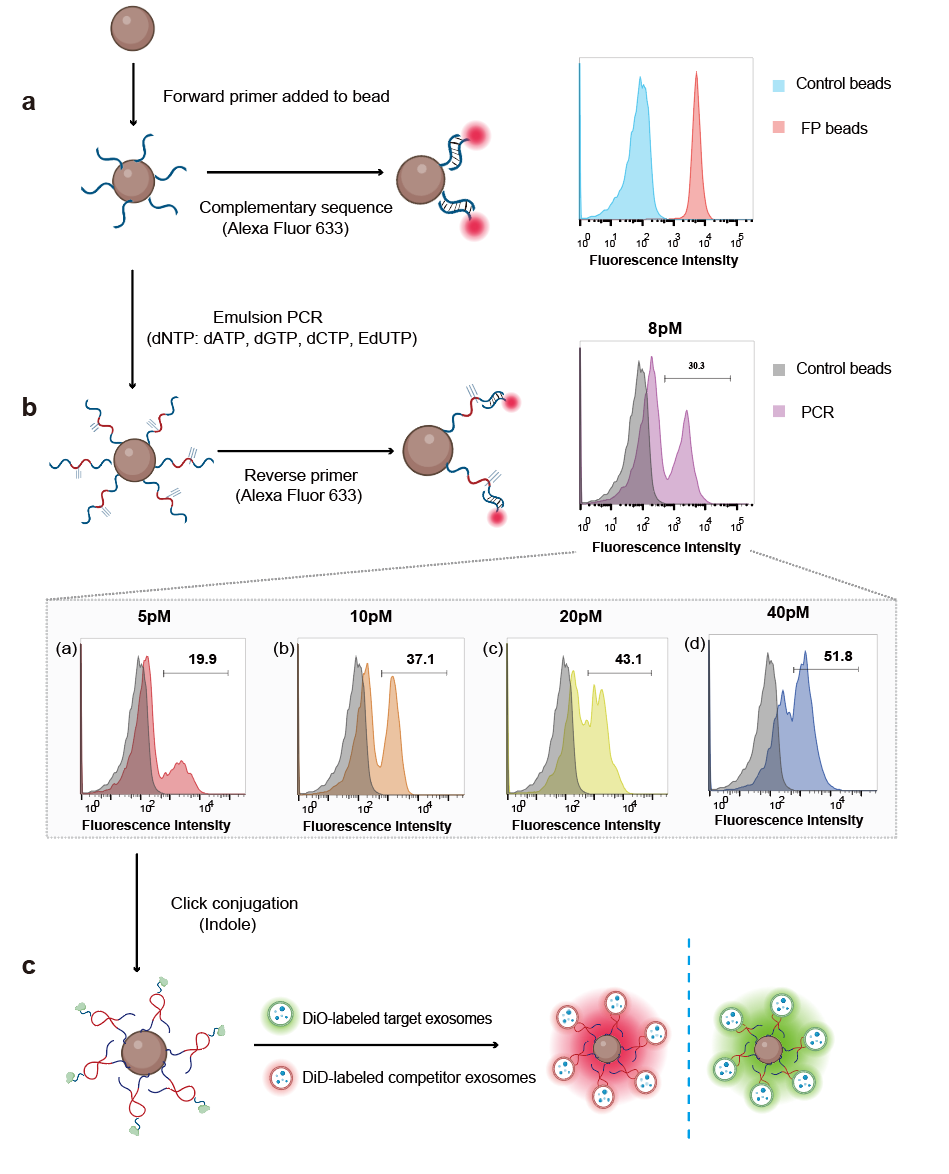

**Supplementary Fig. S2. Library construction and validation of Indole-conjugated library on microbeads. a** Magnetic microbeads are coated with forward primers, and the Alexa Fluor 633 modified complementary sequence was used to verify the conjugation efficiency of the forward primer to the microbeads. **b** Emulsion PCR is performed to create aptamer particles. A reverse primer modified with Alexa Fluor 633 is used to hybridize with the DNA library to assess the efficiency of emulsion PCR. Template concentration optimization is used to determine the appropriate library concentration for PCR amplification. **c** Indole modifications are introduced into the library by click chemistry and incubated with corresponding fluorescently labeled exosomes to prepare samples for sorting.

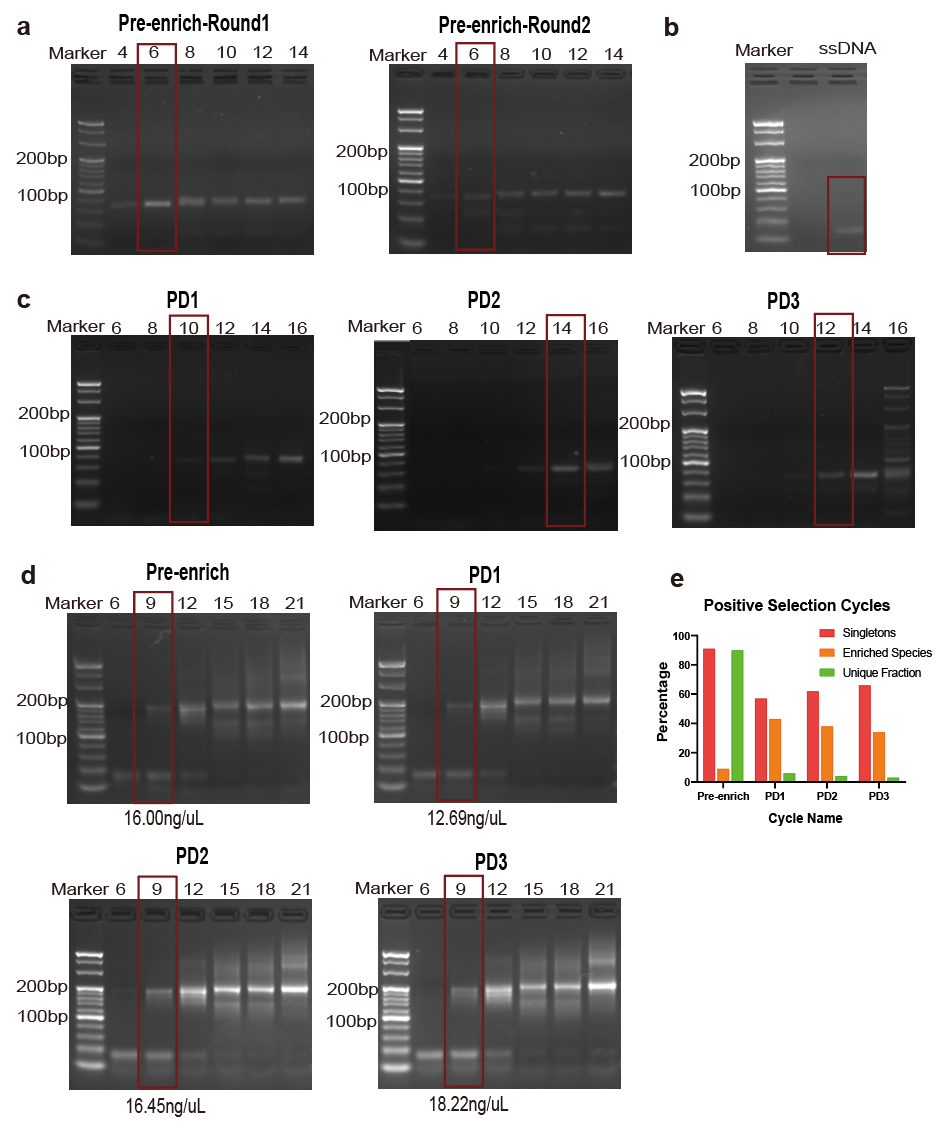

**Supplementary Fig. S3. Library template optimization scheme for microbead-displayed aptamer screening strategy targeting HCC-specific exosomes. a** Agarose gel image of the number of rounds of amplification optimization during the two-round pre-enrichment process. **b** Single-stranding of library double-stranded DNA during screening. **c** The agarose gel profile was optimized for the number of rounds amplified by "reverse" PCR during three rounds of PD screening. **d** Optimization of the number of rounds and concentration determination of sequencing samples. **e** Sequencing results of Aptamer screening targeting exosomal surface.

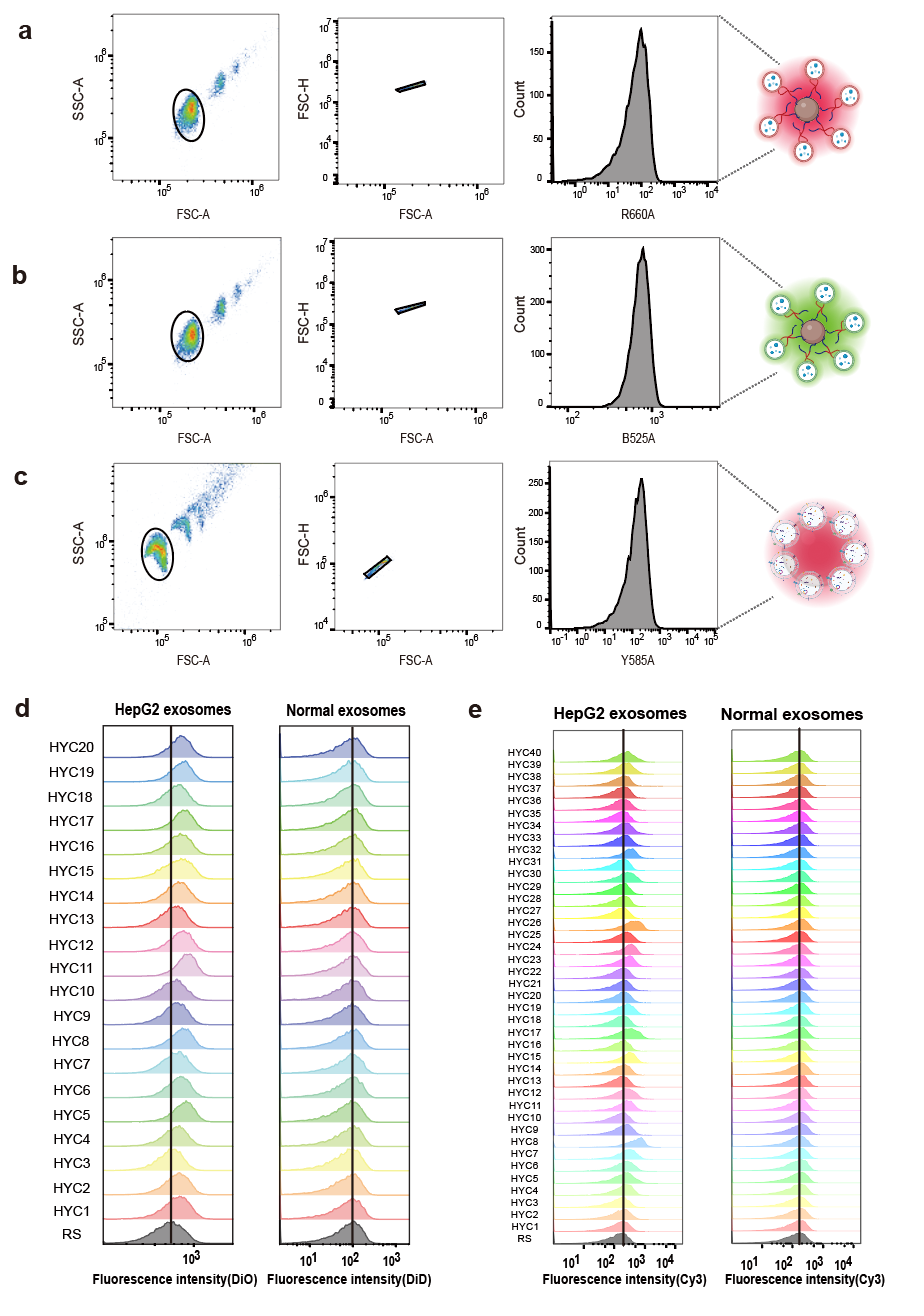

**Supplementary Fig. S4. Flow cytometry verifies the specific binding of selected aptamers. a** The flow cytometry gating strategy for binding the selected aptamer to DiD-labeled normal human plasma-derived exosomes. **b** The flow cytometry gating strategy for binding the selected aptamer to DiO-labeled HepG2 exosomes. **c** The flow cytometry gating strategy for binding the exosomes-coated latex microbeads and Cy3-labeled aptamer. **d** Flow cytometric validation of the top 20 sequences using the magnetic microbead method. **e** Flow cytometric validation of the top 40 sequences using the latex microbead method.

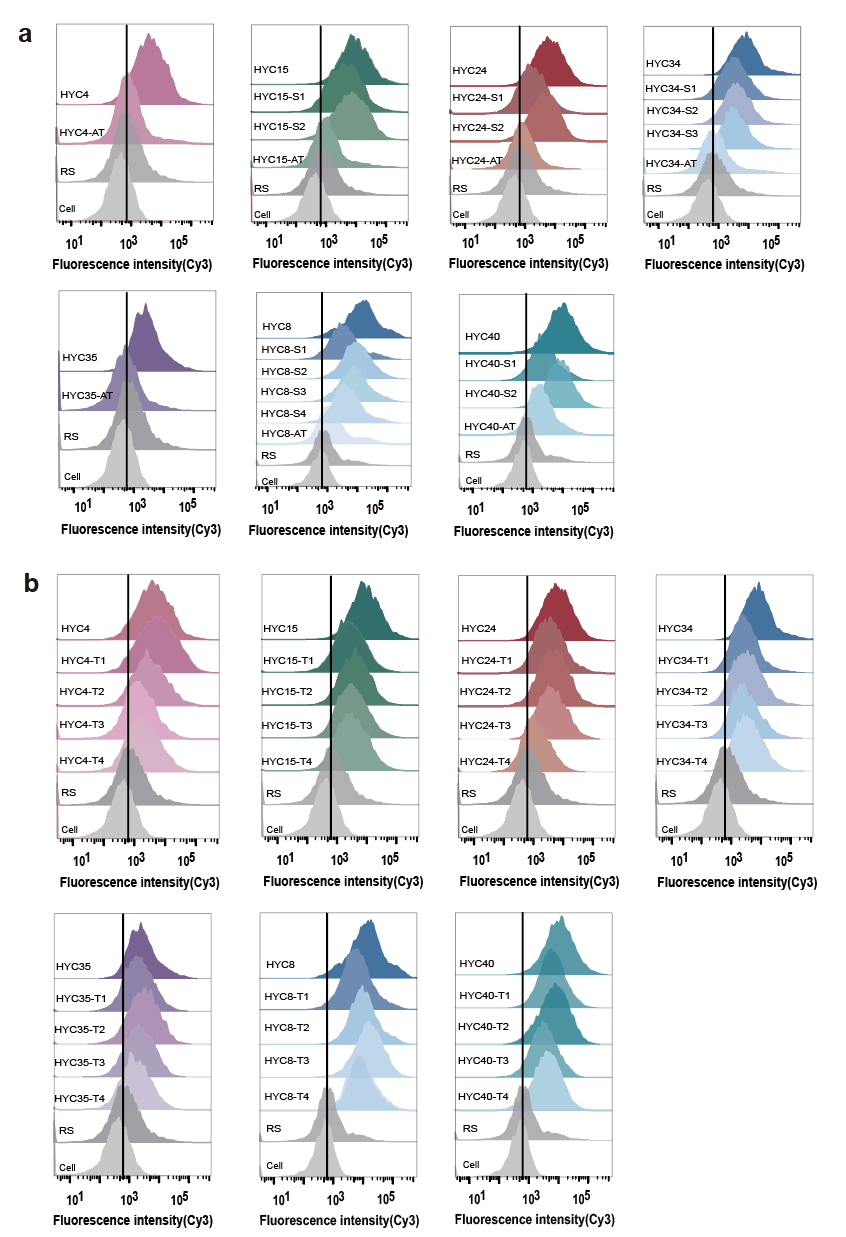

**Supplementary Fig. S5. Identification of the binding sites of the aptamers. a** Flow cytometry analysis showing the binding affinity of aptamer with IndU sites replaced by T individually and all replaced by T. **b** Flow cytometry analysis showing the binding affinity of aptamer and 4 truncated variants.

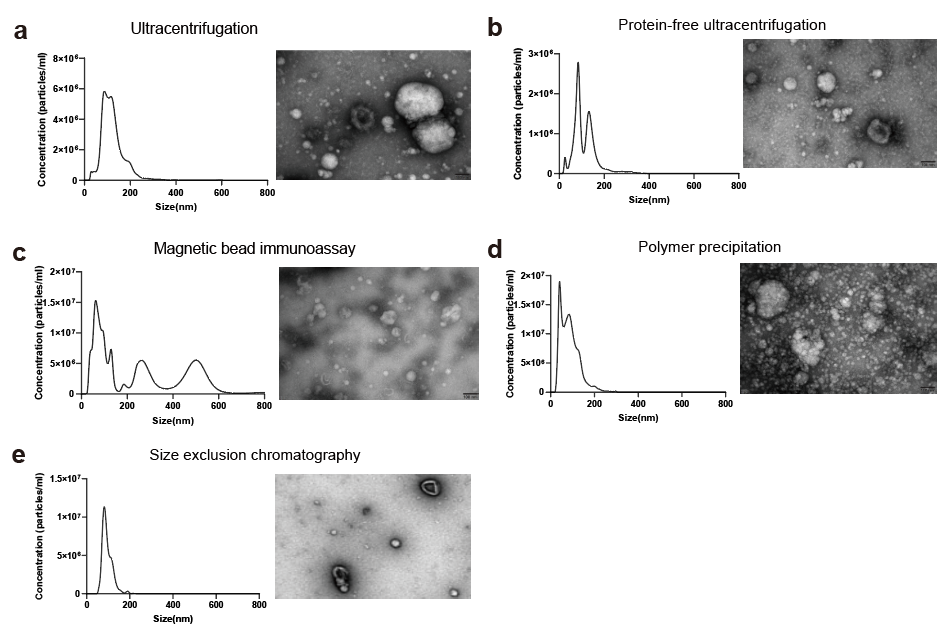

**Supplementary Fig. S6. Isolation and Characterization of Plasma-Derived Exosomes. a** - **e** The ultracentrifugation (UC), Protein-free ultracentrifugation, Magnetic microbead immunoassay, Polymer precipitation, and size-exclusion chromatography (SEC) methods were used to isolate plasma-derived exosomes. The characterization of plasma-derived exosomes was performed using NTA and cryo-TEM imaging.

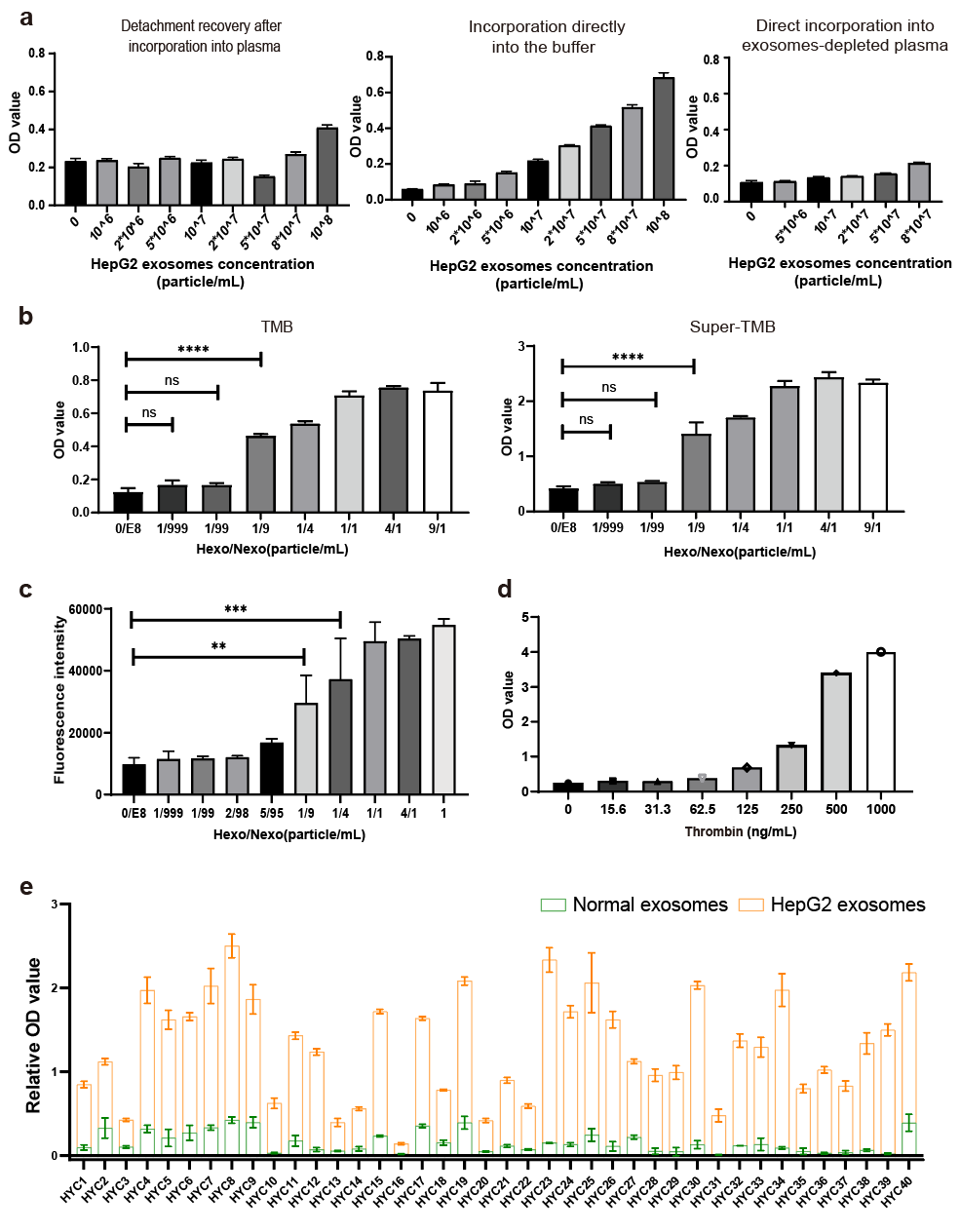

**Supplementary Fig. S7. Development of the aptamer-based ELISA assay for clinical translation.** **a** Exploration of the detection system. The detection of exosomes after incorporation into plasma and subsequent recovery, direct detection in Binding buffer E, and plasma dilution mixed with depleted exosomes were respectively designed. **b** HRP-based TMB chromogenic reaction. Two commercially available TMB chromogenic solutions and ultrasensitive chromogenic solutions were tested. Error bars denote ±SD of the mean for n=3 independent replicates. The p-value was determined by Student’s t-test: ns: not significant; ****p < 0.0001. **c** HRP-based peroxidase fluorescence detection method. Error bars denote ±SD of the mean for n=3 independent replicates. The p-value was determined by Student’s t-test: **p < 0.01; ***p < 0.001. **d** Exploration of thrombin concentrations. **e** The relative OD value at 450nm for each aptamer from normal human plasma-derived exosomes and HepG2 exosomes. Error bars denote ±SD of the mean for n=3 independent replicates.

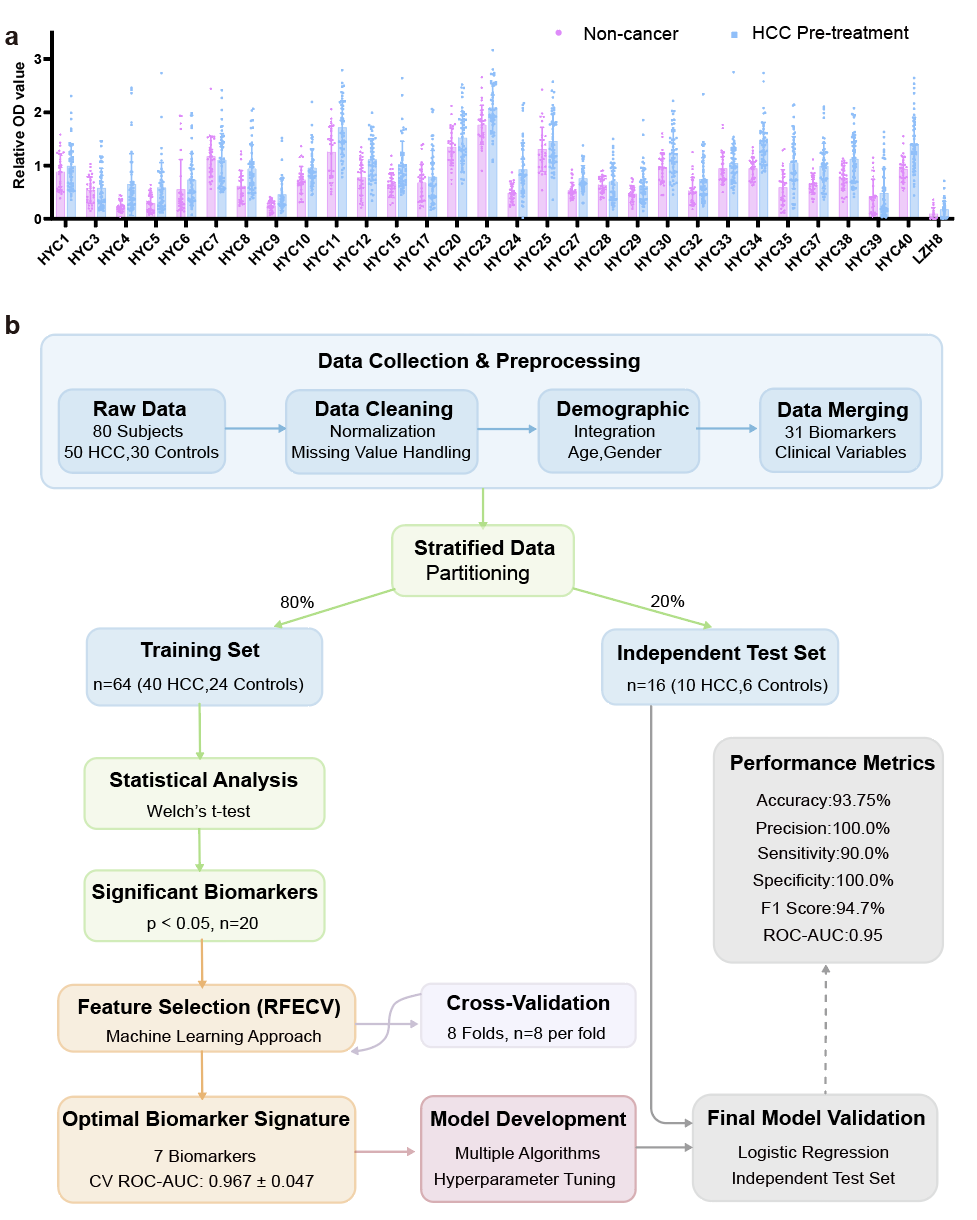

**Supplementary Fig. S8. Application of aptamer-based ELISA in diagnosis and machine learning. a** The relative OD value at 450 nm for aptamers from healthy volunteer plasma-derived exosomes and HCC pre-treatment patient-derived exosomes. Non-cancer, n=30; HCC pre-treatment, n=50. **b** The complete analysis workflow implemented for biomarker discovery and model development.

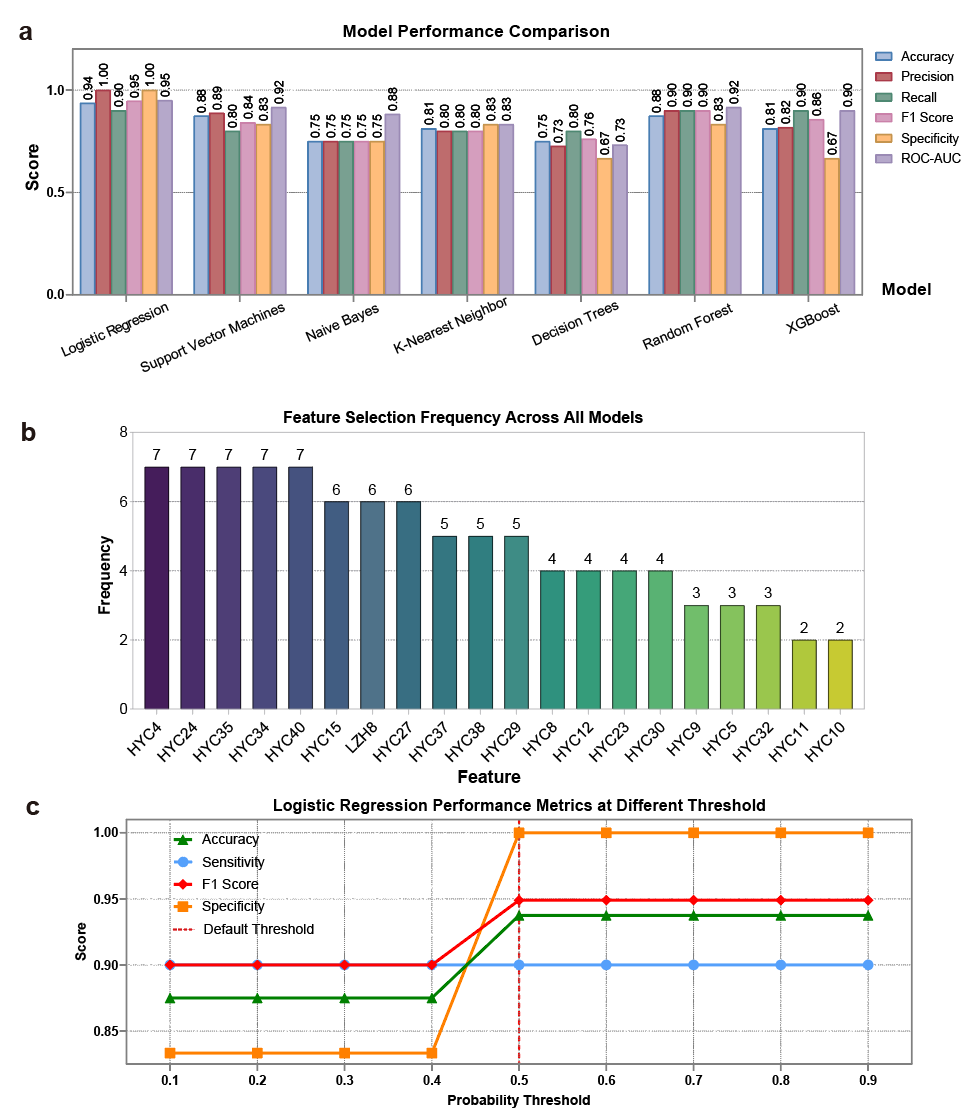

**Supplementary Fig. S9. Machine learning model performance comparison and model threshold analysis.** **a** Comparative analysis of multiple machine learning approaches for HCC diagnosis using biomarker data. Seven classification algorithms were evaluated on an independent test set (n=16) following 8-fold cross-validation and feature selection during model development. Five key performance metrics are shown: Accuracy, Precision, Recall (Sensitivity), F1 Score, and ROC-AUC. Logistic Regression demonstrated superior performance with 93.75% accuracy, 100% precision, and 90% recall, achieving perfect specificity with no false positives. This model was selected for the final 7-biomarker diagnostic signature presented in the main text. **b** Feature frequency analysis across different machine learning models. The bar chart represents how frequently each biomarker was selected as an important feature by different algorithms. Six biomarkers (HYC4, HYC15, HYC24, HYC34, HYC35, and HYC40) were consistently identified across multiple feature selection methods, underscoring their robustness as diagnostic indicators. **c** Performance metrics of the 7-biomarker logistic regression model across probability thresholds (0.1-0.9). At the 0.5 threshold (vertical dashed line), specificity reaches 100% while sensitivity remains at 90%, with optimal accuracy (93.8%) and F1 score (94.7%). This threshold was selected for the final diagnostic model to maximize clinical utility by eliminating false positives while maintaining high detection sensitivity.

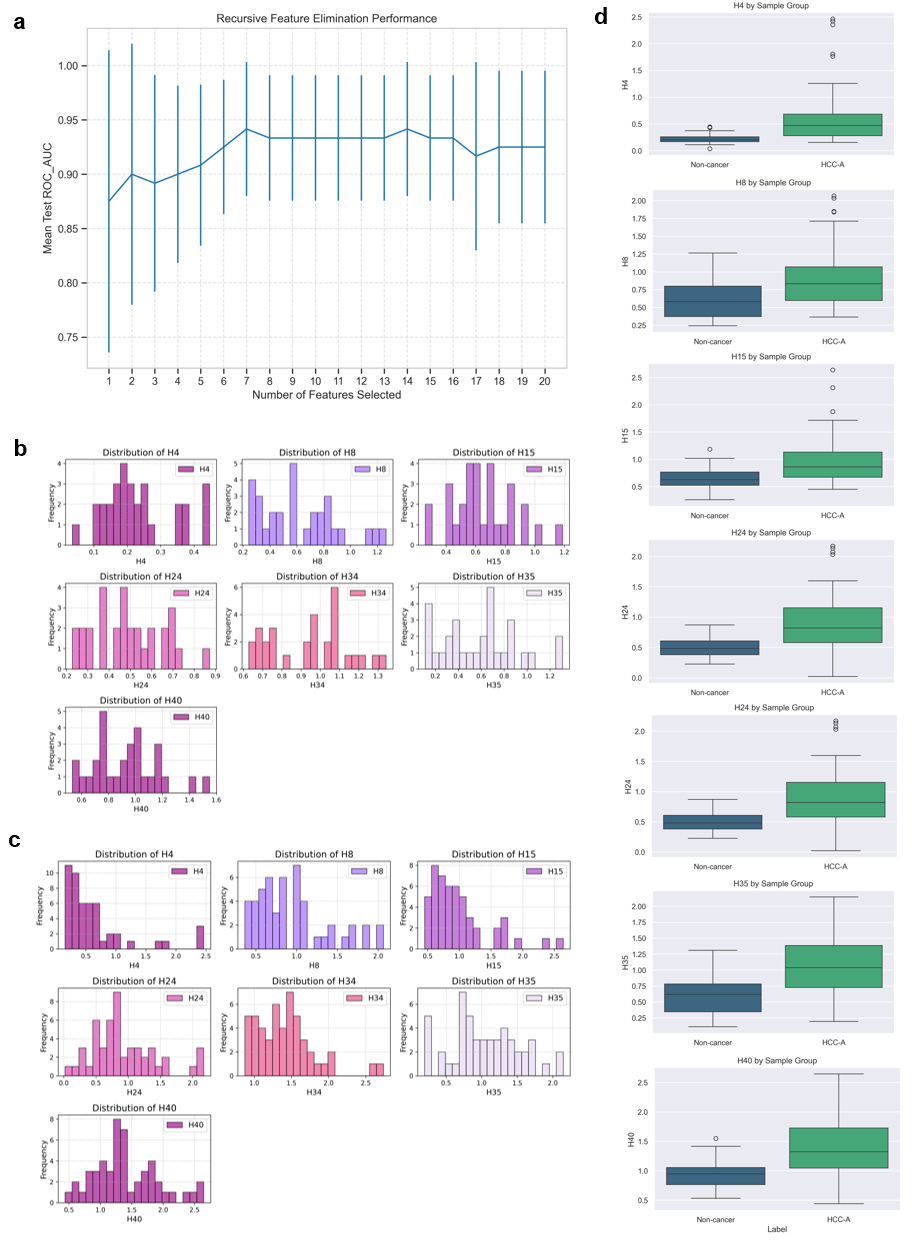

**Supplementary Fig. S10. Feature selection process, individual biomarker distributions, and histograms of selected biomarkers.** **a** Recursive Feature Elimination with Cross-Validation (RFECV) results showing model performance as a function of the number of features. Performance plateaued with 7 biomarkers, indicating an optimal balance between model complexity and predictive power. Cross-validation ROC-AUC of the feature subset was 0.9667 ± 0.0471. **b** and **c** Distribution histograms of the 7 selected biomarkers in **b** non-cancer controls and **c** HCC patients. These visualizations demonstrate the distinct expression patterns between groups, particularly for biomarker H4 which showed the greatest separation. **d** Boxplots showing the distribution of the 7 selected biomarkers in HCC patients versus non-cancer controls. All selected biomarkers show significant differences between groups (p < 0.05). The "H" number shown in the figure corresponds to the "HYC" number in the text.

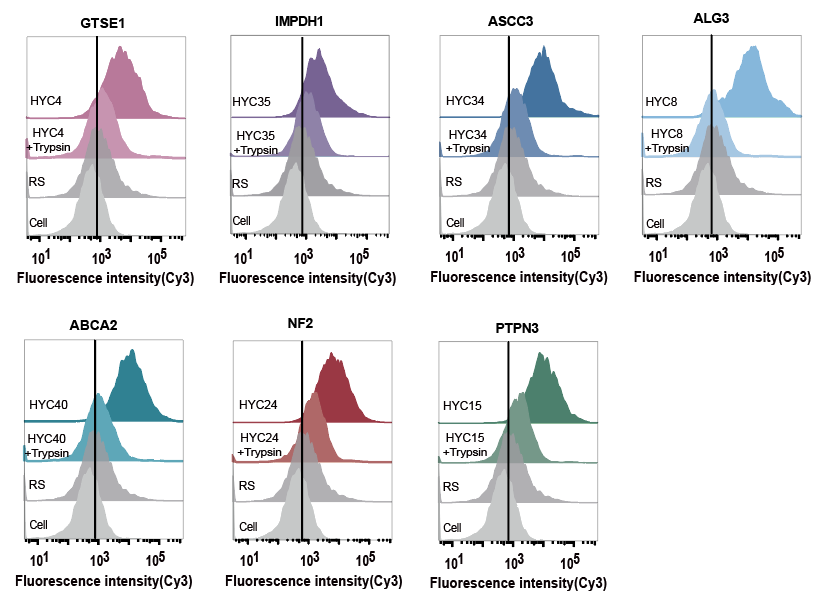

**Supplementary Fig. S11. Identification of the binding target of the aptamers.** Characterization of target type through cell membrane protein digested by trypsin.

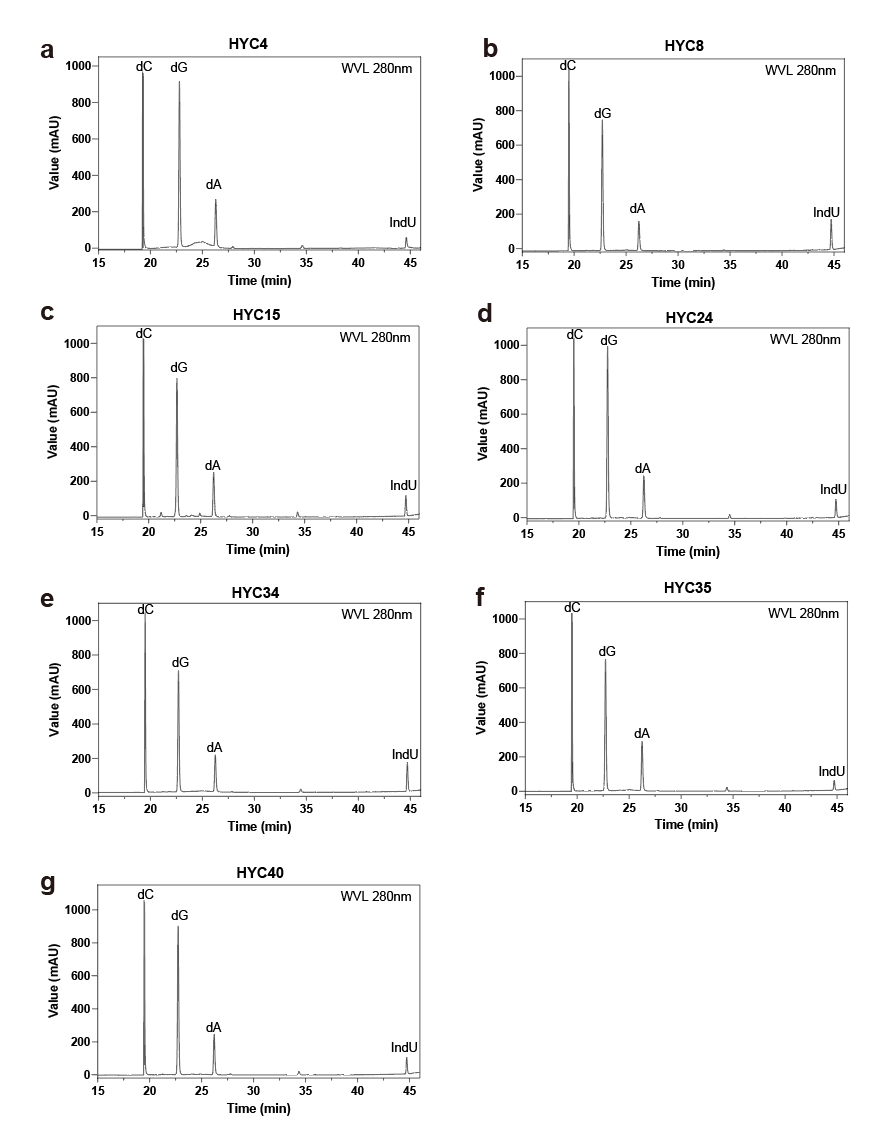

**Supplementary Fig. S12. High Performance Liquid Chromatography (HPLC) analysis of digest products of modified DNA.**

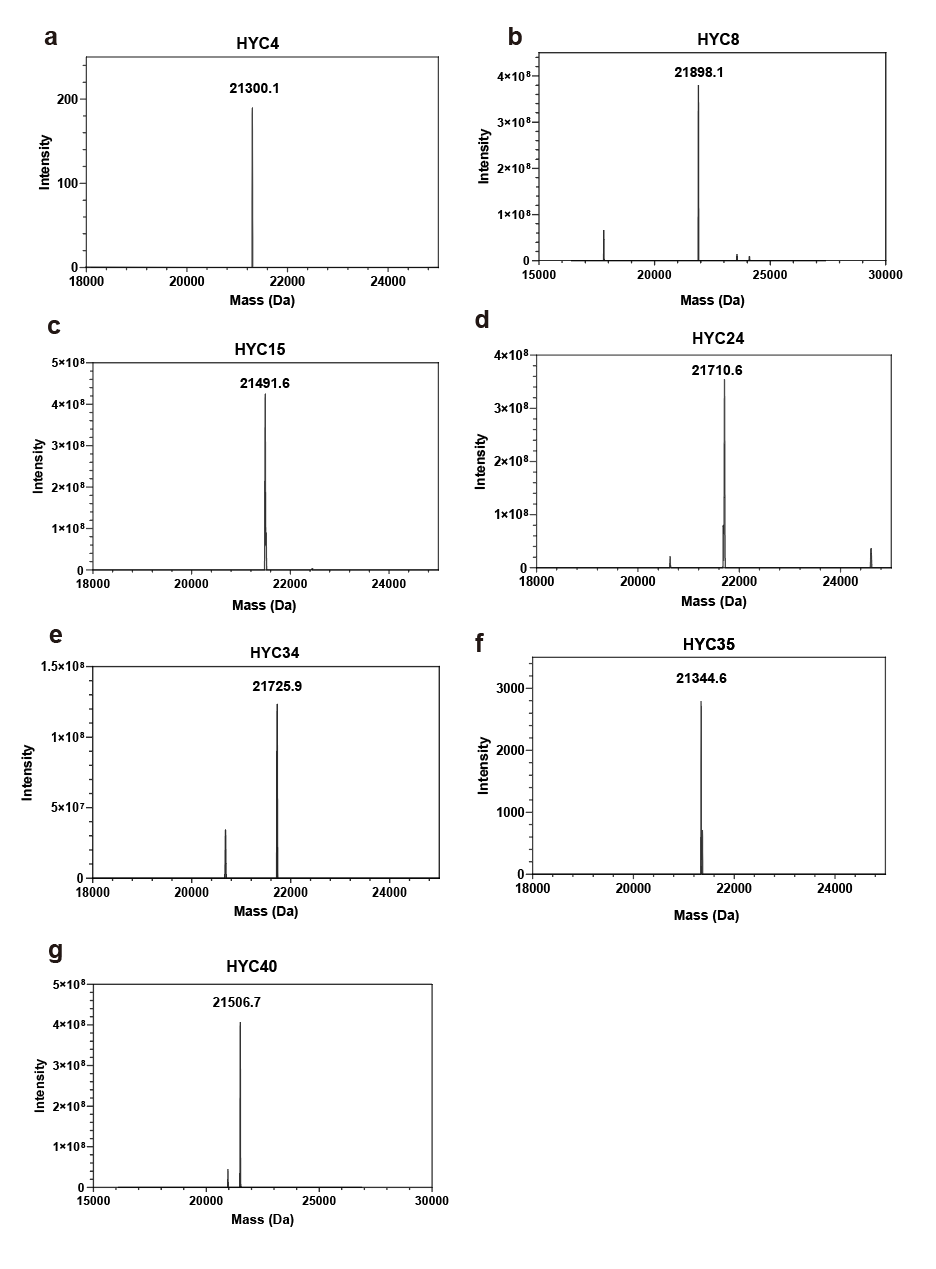

**Supplementary Fig. S13. Mass Spectrometry results of the aptamers.**

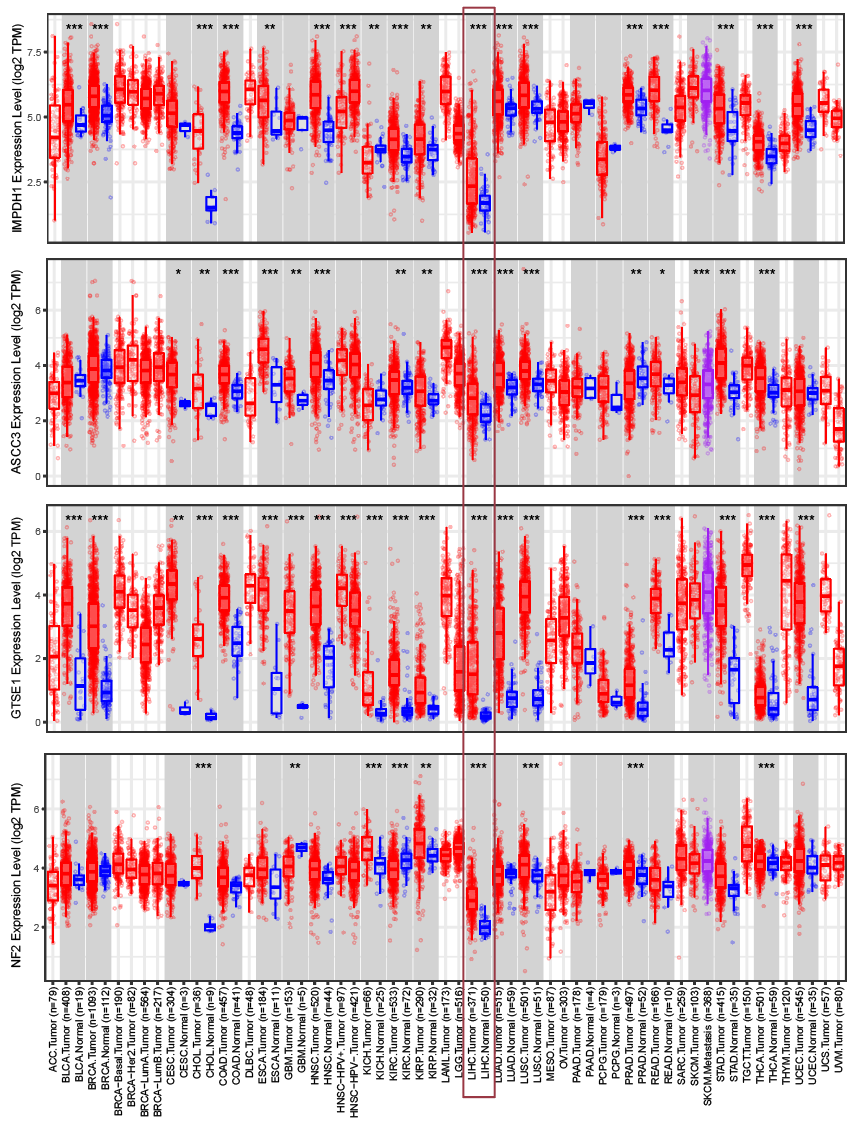

**Supplementary Fig. S14. Differential gene expression analysis.** Differential gene expression analysis revealed significant upregulation of all validated markers in LIHC tissues compared with adjacent non-tumor tissues. LIHC, liver hepatocellular carcinoma.

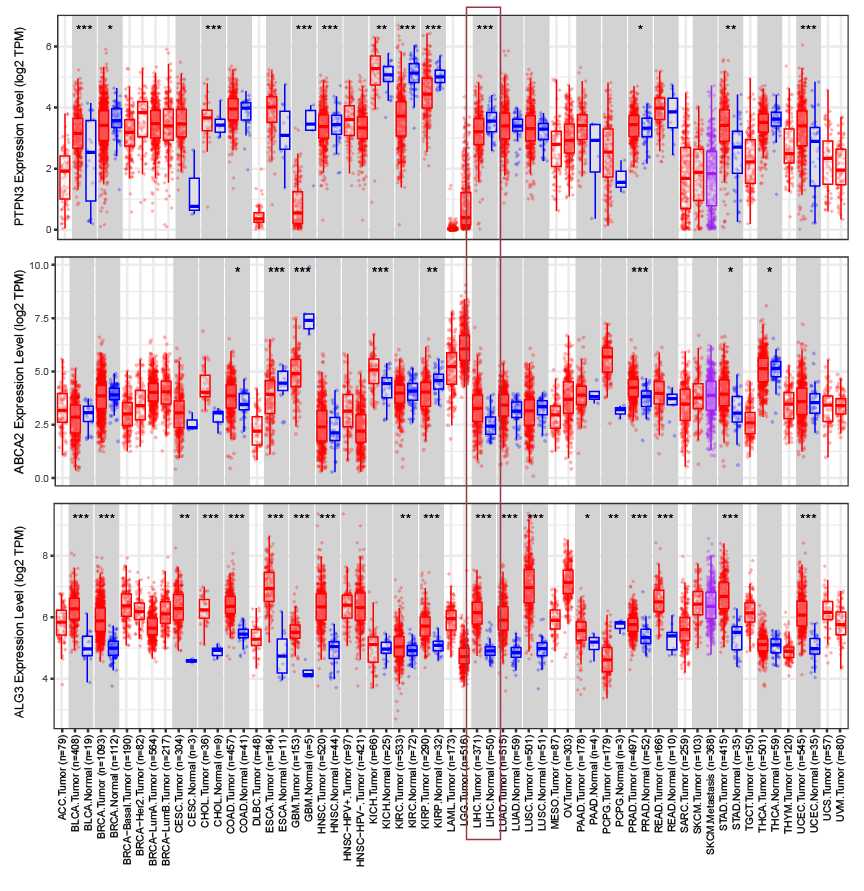

**Supplementary Fig. S14 Differential gene expression analysis (Continued).** Differential gene expression analysis revealed significant upregulation of all validated markers in LIHC tissues compared to adjacent non-tumor tissues, except for ABCA2. LIHC stands for liver hepatocellular carcinoma.

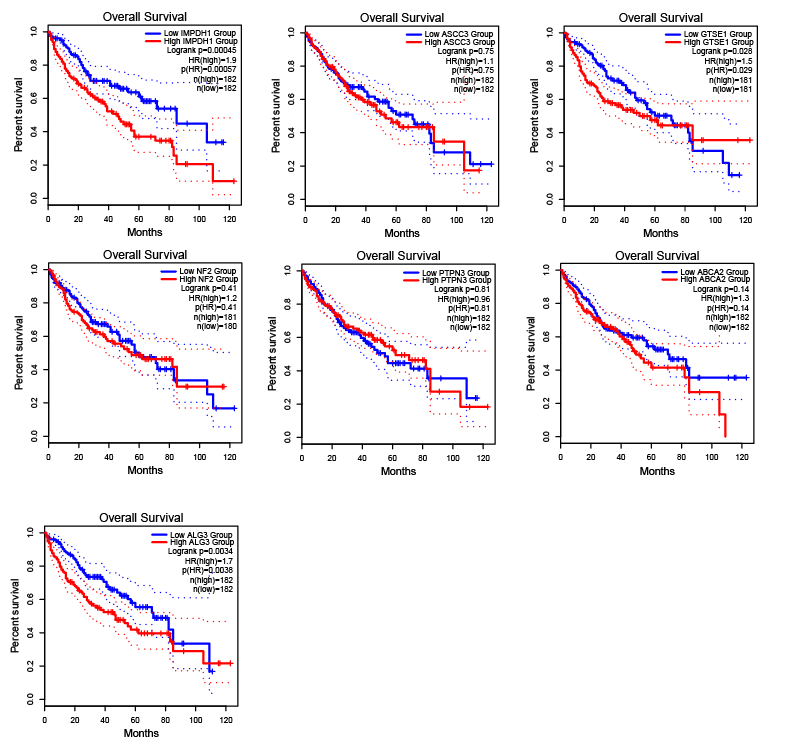

**Supplementary Fig. S15. Survival Analysis.** Survival analysis demonstrated that elevated expression of these markers was significantly associated with reduced overall survival in LIHC patients.

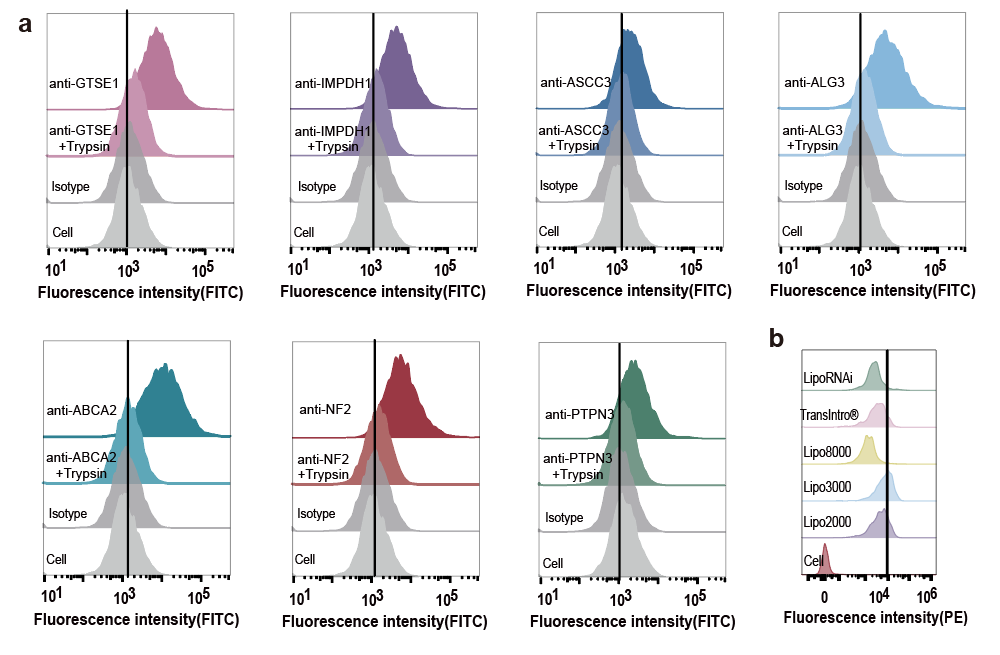

**Supplementary Fig. S16. Identification of the binding target by antibodies and transfection efficiency. a** Identification of the binding target by their antibodies and confirmation of the target type through cell membrane protein digested by trypsin. **b** Transfection reagent for HepG2 cells was tested for transfection efficiency.

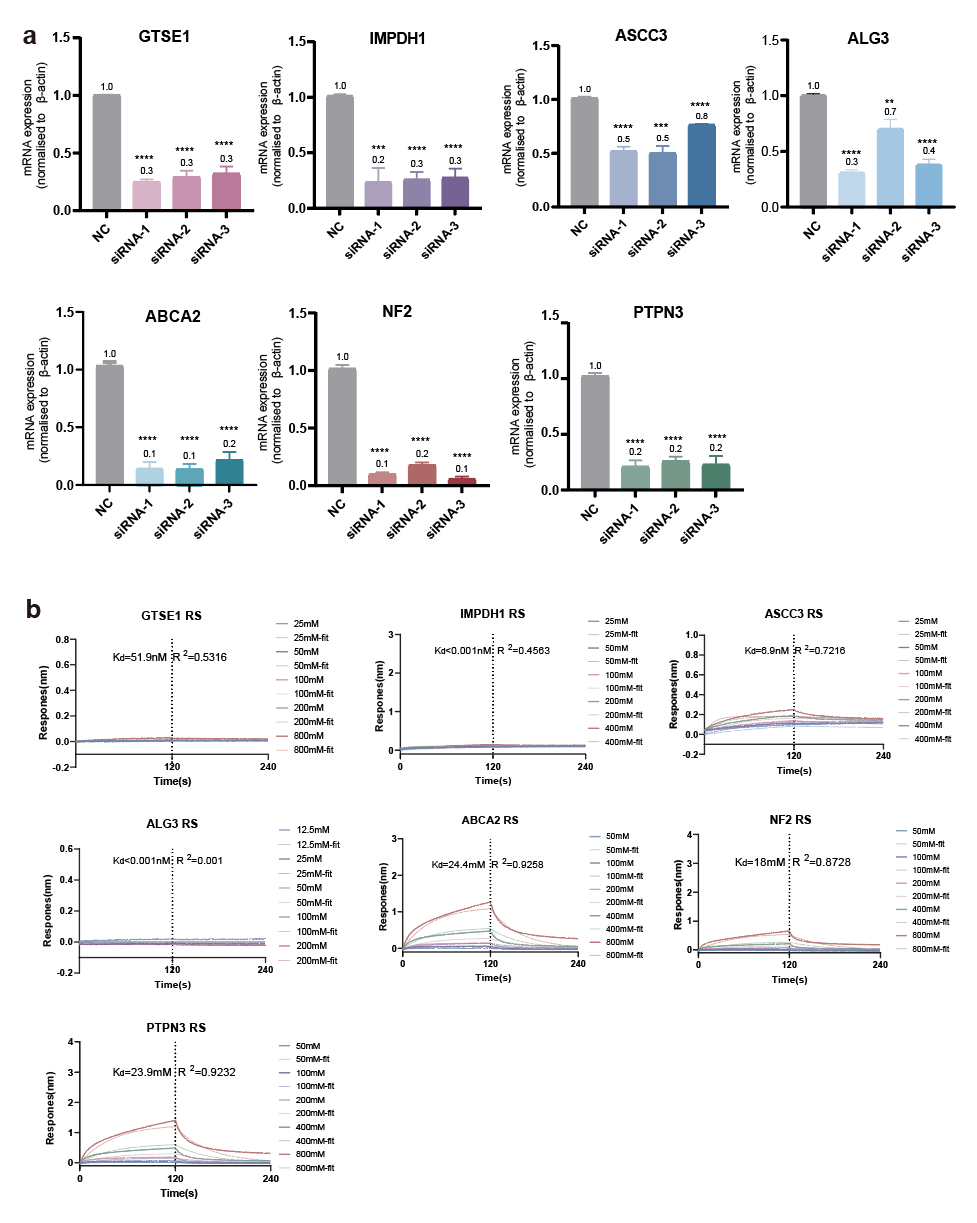

**Supplementary Fig. S17. Molecular validation of aptamer-binding targets. a** The qPCR analysis showing that the mRNA expression level of the target proteins GTSE1, IMPDH1, ASCC3, ALG3, ABCA2, NF2, and PTPN3. (n = 3, *p < 0.05; **p < 0.01; ***p < 0.001; ****P < 0.0001; unpaired Student’s t test). **b** the BLI assay determining the Kd value of targets and random sequence (RS).

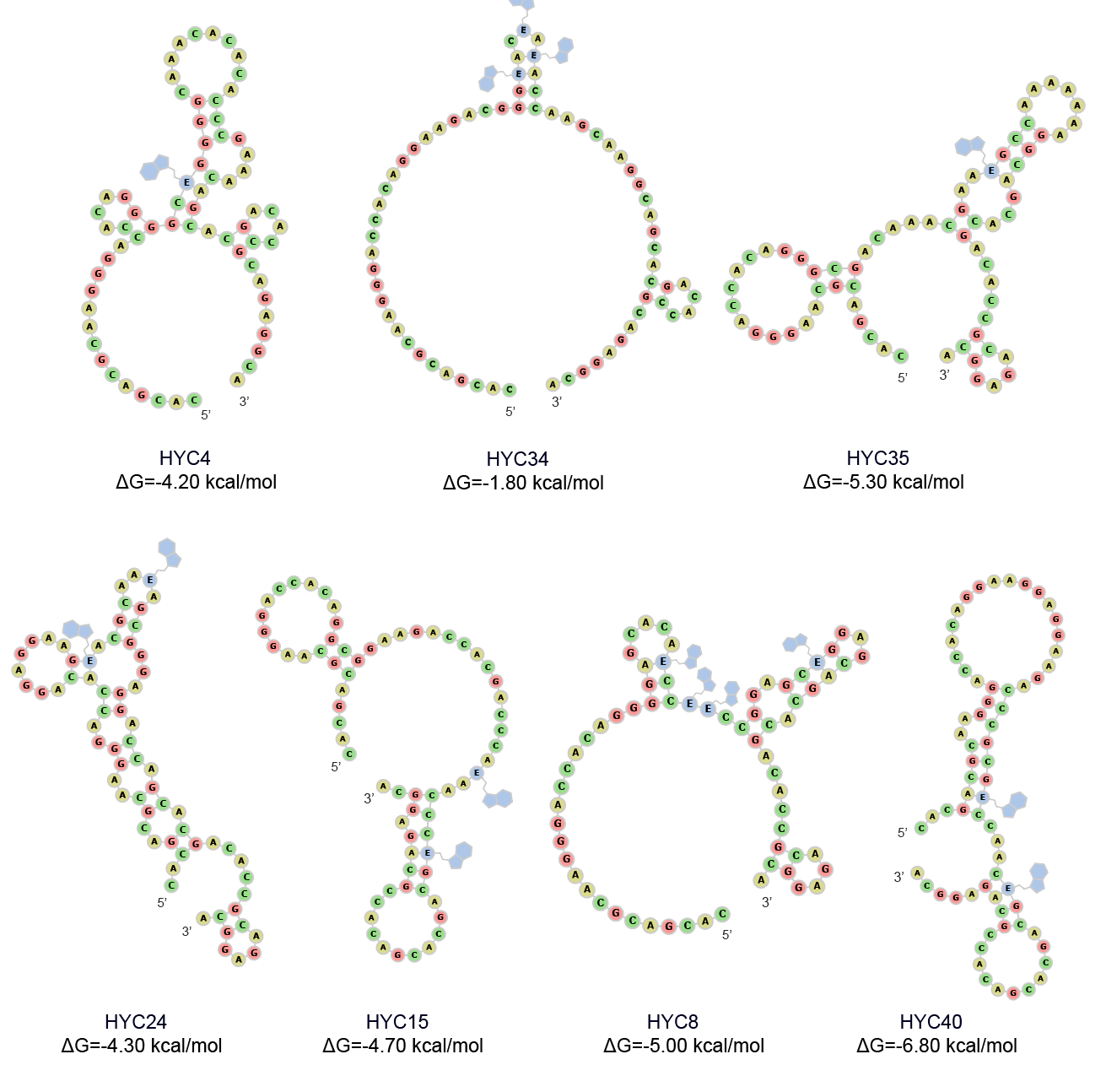

**Supplementary Fig. S18. Predicted 2-dimentional structure of aptamers.**

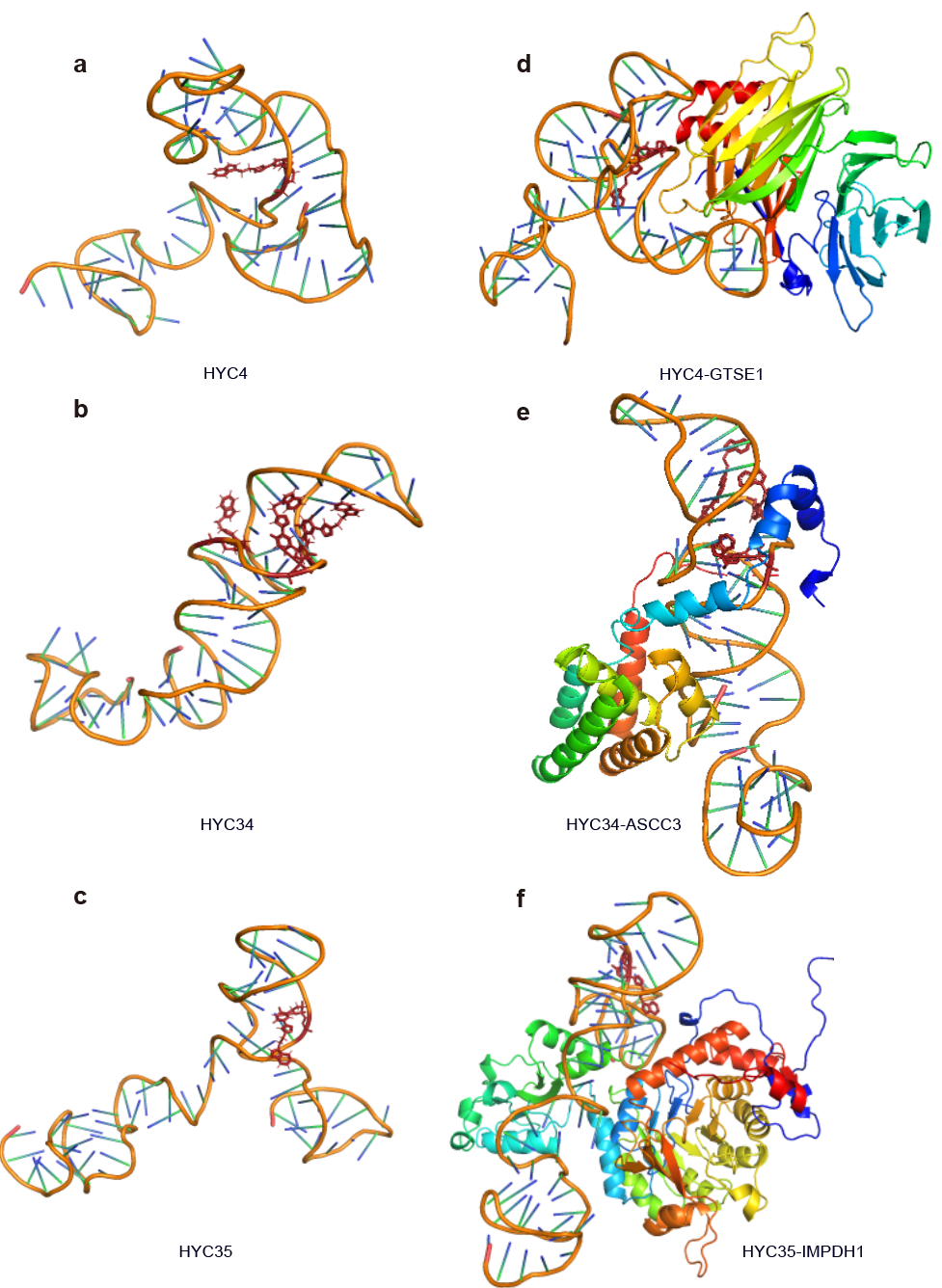

**Supplementary Fig. S19. Predicted tertiary structure of aptamers, Molecular Simulations of Aptamer and protein-binding complexes.**

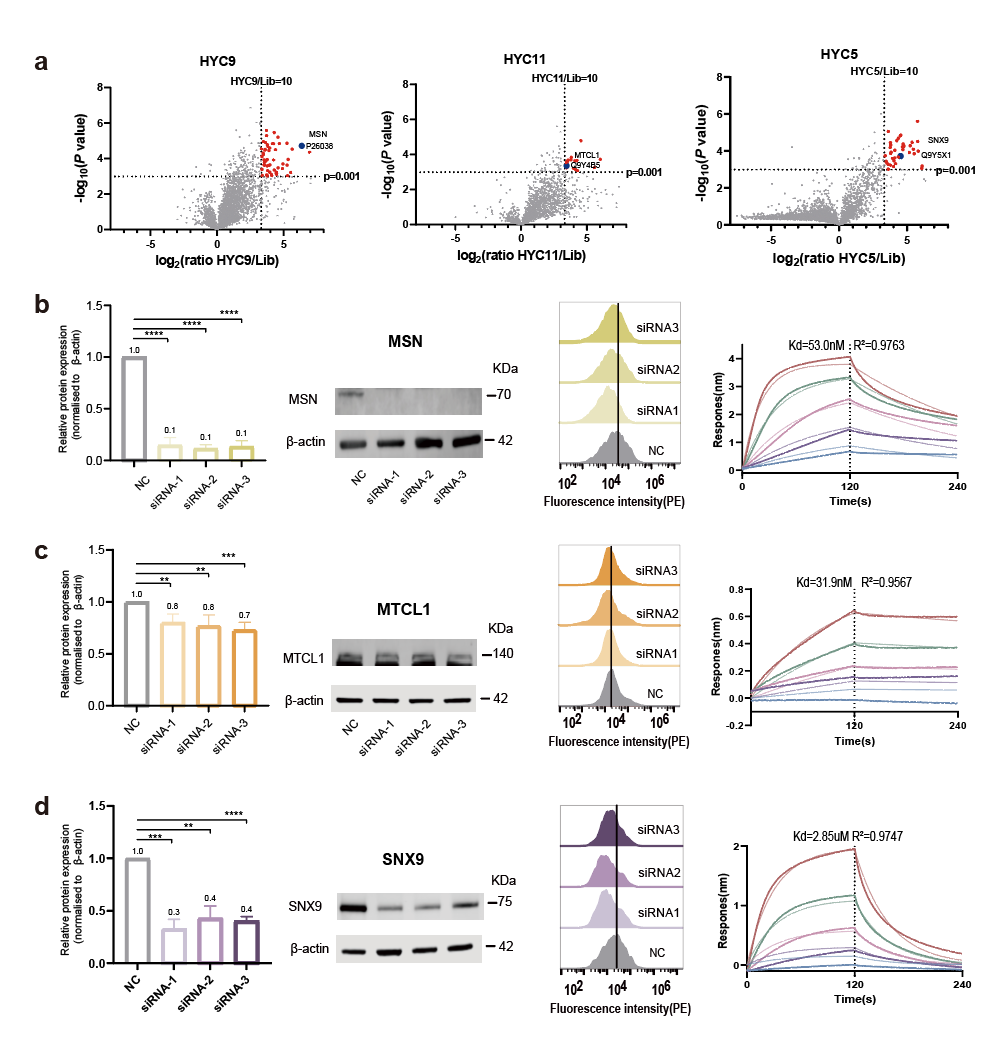

**Supplementary Fig. S20. Molecular validation of three additional aptamer-binding targets.** **a** Volcano diagram analysis of differential proteins based on proteomics. log2 (ratio) > 3.32, P value < 0.0001. **b** - **d** From top to bottom: target proteins MSN, MTCL1, and SNX9. The verification of each target protein from left to right is as follows: Typical WB images and quantitative analysis showing that the protein expression level was reduced by siRNA (n = 3, **p < 0.01; ***p < 0.001; ****p < 0.0001; unpaired Student’s t test); Flow cytometry analysis of the binding ability of aptamer on knockdown HepG2 cells; and the BLI assay determining the Kd value of targets and aptamers.

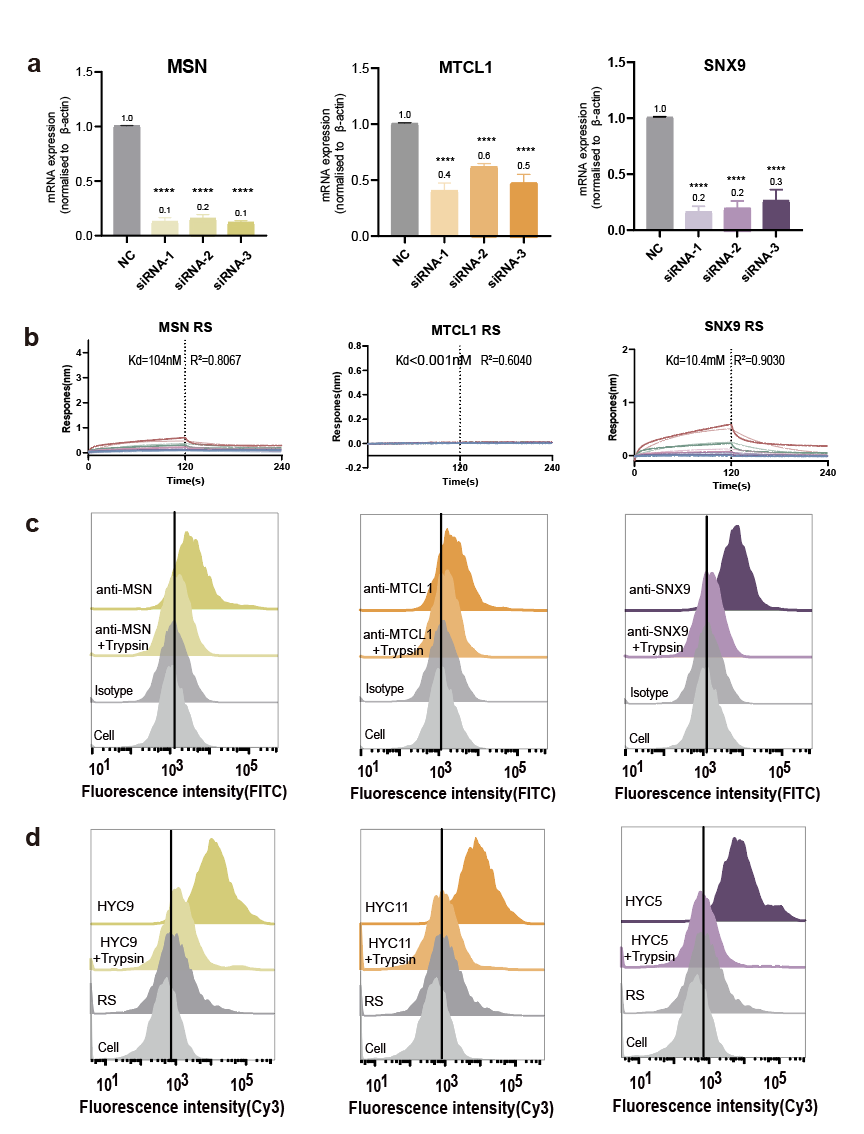

**Supplementary Fig. S21. Molecular validation of three additional aptamer-binding targets. a** qPCR analysis showing the mRNA expression levels of the target proteins MSN, MTCL1, and SNX9 (n = 3, ****P < 0.0001; unpaired Student’s t-test). **b** BLI assay determining the Kd value of targets and random sequence (RS). **c** Analysis of target type through cell membrane protein digestion by trypsin. **d** Further analysis of target type through cell membrane protein digestion by trypsin.

**Supplementary Tables**

**Supplementary Table S1.** Demographic and Clinical Characteristics of HCC Patients and healthy volunteers in Aptamer-Based ELISA Validation Cohort.

| **Characteristic** | **Healthy volunteers (*n* = 30)** | **HCC Patients (Pre-treatment, *n* = 50)** | **HCC Patients (Post-treatment, *n* = 30)** |
| --- | --- | --- | --- |
| Mean age, years±SD | 57±10.9 | 60±9.3 | 62±7.6 |
| Sex |  |  |  |
| Male, n (%) | 24 (80.0%) | 42 (82.6%) | 25 (83.3%) |
| Female, n (%) | 6 (20.0%) | 9 (17.6%) | 5 (16.7%) |
| Smoking^§^ |  |  |  |
| Never, *n* (%) | 17 (56.7%) | 25 (50.0%) | 16 (53.3%) |
| Former, *n* (%) | 3 (10.0%) | 12 (24.0%) | 6 (20.0%) |
| Current, *n* (%) | 10 (33.3%) | 13 (26.0%) | 8 (26.7%) |
| Drinking^†^ |  |  |  |
| Never, *n* (%) | 17 (56.7%) | 38 (76.0%) | 24 (80.0%) |
| Former, *n* (%) | 1 (3.3%) | 9 (18.0%) | 4 (13.3%) |
| Current, *n* (%) | 12 (40.0%) | 3 (6.0%) | 2 (6.7%) |
| BMI^\|\|^ |  |  |  |
| Thin, n (%) | 1 (3.3%) | 0 (0%) | 0 (0%) |
| Normal, n (%) | 9 (30.0%) | 19 (38.0%) | 12 (40.0%) |
| Overweight, n (%) | 16 (53.4%) | 20 (40.0%) | 11 (36.7.0%) |
| Obese, n (%) | 4 (13.3%) | 11 (22.0%) | 7 (23.3%) |
| Diabetes^#^ |  |  |  |
| Yes, *n* (%) | 0 (0%) | 42 (84.0%) | 26 (86.7%) |
| No, *n* (%) | 30 (100%) | 8 (16.0%) | 4 (13.3%) |
| Hepatitis^‡^ |  |  |  |
| Yes, *n* (%) | 0 (0%) | 47 (94.0%) | 27 (90.0%) |
| No, *n* (%) | 30 (100%) | 3 (6.0%) | 3 (10.0%) |

Blank entries indicate “not applicable.” ^§^Smoking status at time of diagnosis. ^†^Drinking status at time of diagnosis. ^||^ BMI<18.5, Thin; 18.5≤BMI<24, Normal; 24≤BMI<28, Overweight; 28≤BMI<32, Obese. ^‡^Includes both viral and nonviral hepatitis.

**Supplementary Table S2.** Sequences used in the screening.

| Name | Sequence (5′-3′) |
| --- | --- |
| Initial library | CACGACGCAAGGGACCACAGG-𝑁25-CAGCACGACACCGCAGAGGCA - |
| Competing library | GATCCCAGTCCGAAGTAATCG-𝑁25-GCTTGTGCAAACGGCTATAGG |
| Forward primer | CACGACGCAAGGGACCACAGG |
| Reverse primer | Phosphate-TGCCTCTGCGGTGTCGTGCTG |

Note: N=A; EdU; G; C.

**Supplementary Table S3.** Primers for selection library sequencing.

| Primer | Sequence (5′-3′) |
| --- | --- |
| CHY- FP | AATGATACGGCGACCACCGAGATCTACACTCTTTCCCTACACGACGCTCTTCCGATCTCACGACGCAAGG |
| CHY-RP-Pre-enrich | CAAGCAGAAGACGGCATACGAGATTCGCCTTAGTGACTGGAGTTCAGACGTGTGCTCTTCCGATCTTGCCTCTGCGGT |
| CHY-RP-PD1 | CAAGCAGAAGACGGCATACGAGATCTAGTACGGTGACTGGAGTTCAGACGTGTGCTCTTCCGATCTTGCCTCTGCGGT |
| CHY-RP-PD1 | CAAGCAGAAGACGGCATACGAGATTTCTGCCTGTGACTGGAGTTCAGACGTGTGCTCTTCCGATCTTGCCTCTGCGGT |
| CHY-RP-PD1 | CAAGCAGAAGACGGCATACGAGATGCTCAGGAGTGACTGGAGTTCAGACGTGTGCTCTTCCGATCTTGCCTCTGCGGT |

Note: The underlined sequences are the index for each round.

**Supplementary Table S4.** Sequence information of aptamer libraries for high-throughput phenotypic screening

| ID | Sequence (5′-3′) |
| --- | --- |
| HYC1 | CACGACGCAAGGGACCACAGGAACGGGGGCAGGAAACCGAGGEGGGCAGCACGACACCGCAGAGGCA |
| HYC2 | CACGACGCAAGGGACCACAGGCGCGGCCAAAAAAGAAGCACACEGGCAGCACGACACCGCAGAGGCA |
| HYC3 | CACGACGCAAGGGACCACAGGGGAGGAGEAGGAAGCAAGGACCGCACAGCACGACACCGCAGAGGCA |
| HYC4 | CACGACGCAAGGGACCACAGGGGCEGGGGCAAACACACACCCGAAACAGCACGACACCGCAGAGGCA |
| HYC5 | CACGACGCAAGGGACCACAGGAACCGACCCCCEECACAACGGGACCCAGCACGACACCGCAGAGGCA |
| HYC6 | CACGACGCAAGGGACCACAGGGGCAEAGCCCCCAAACGGCCGECACCAGCACGACACCGCAGAGGCA |
| HYC7 | CACGACGCAAGGGACCACAGGCECCACGAAAGAGCAGGGCAEGCAACAGCACGACACCGCAGAGGCA |
| HYC8 | CACGACGCAAGGGACCACAGGGGAGCACAECCEECCGGAGCEGGAGCAGCACGACACCGCAGAGGCA |
| HYC9 | CACGACGCAAGGGACCACAGGCCGEAECCAGGCACGACAGCGAGAGCAGCACGACACCGCAGAGGCA |
| HYC10 | CACGACGCAAGGGACCACAGGCACCGGEGCGGACCACGCACAGGCACAGCACGACACCGCAGAGGCA |
| HYC11 | CACGACGCAAGGGACCACAGGAAGGAAGGGGAAGGCGEAGEGGGCACAGCACGACACCGCAGAGGCA |
| HYC12 | CACGACGCAAGGGACCACAGGGCGEGAACGCCEAAECGAGGGGAACCAGCACGACACCGCAGAGGCA |
| HYC13 | CACGACGCAAGGGACCACAGGCEACGCACCCGCCCGGGACACCCGACAGCACGACACCGCAGAGGCA |
| HYC14 | CACGACGCAAGGGACCACAGGCACGGGGCAACAAACCAAAGGGAGCCAGCACGACACCGCAGAGGCA |
| HYC15 | CACGACGCAAGGGACCACAGGCGGAAGACCACGACCCAEAACCCEGCAGCACGACACCGCAGAGGCA |
| HYC16 | CACGACGCAAGGGACCACAGGAGCAGGAGGCGACEAGGAGGCAEAGCAGCACGACACCGCAGAGGCA |
| HYC17 | CACGACGCAAGGGACCACAGGCEACCCCGGEAGAACACAGGACAEACAGCACGACACCGCAGAGGCA |
| HYC18 | CACGACGCAAGGGACCACAGGCEAAAGGCAGCGGCAACCGAGAGCCCAGCACGACACCGCAGAGGCA |
| HYC19 | CACGACGCAAGGGACCACAGGAAGCGGCCGGGEGCGACACCAEGCACAGCACGACACCGCAGAGGCA |
| HYC20 | CACGACGCAAGGGACCACAGGEGAAAGECGGCCAGGGCAECECCGACAGCACGACACCGCAGAGGCA |
| HYC21 | CACGACGCAAGGGACCACAGGGCCEAACGACCAAACGGAGAGCCGGCAGCACGACACCGCAGAGGCA |
| HYC22 | CACGACGCAAGGGACCACAGGEAGACGAAACCEACGGCEAGGEAEGCAGCACGACACCGCAGAGGCA |
| HYC23 | CACGACGCAAGGGACCACAGGGAGCGAAAGGGEGCAEAAACGGAECCAGCACGACACCGCAGAGGCA |
| HYC24 | CACGACGCAAGGGACCACAGGAGGAAGEACGCAAEAGCGGGAGGACCAGCACGACACCGCAGAGGCA |
| HYC25 | CACGACGCAAGGGACCACAGGGGACGEGCCAEAAECGGEAGGACACCAGCACGACACCGCAGAGGCA |
| HYC26 | CACGACGCAAGGGACCACAGGCGCCEAGGGAGAEAACAACCACAAECAGCACGACACCGCAGAGGCA |
| HYC27 | CACGACGCAAGGGACCACAGGGACGACAAAGGGCCCCCCGAAAAGACAGCACGACACCGCAGAGGCA |
| HYC28 | CACGACGCAAGGGACCACAGGAAAAACGGGAAACCGCAGAACGAAECAGCACGACACCGCAGAGGCA |
| HYC29 | CACGACGCAAGGGACCACAGGCCACCGCGACGGGCACAACCGACAECAGCACGACACCGCAGAGGCA |
| HYC30 | CACGACGCAAGGGACCACAGGAAGCACGECCCECCGCGGGAAECCCCAGCACGACACCGCAGAGGCA |
| HYC31 | CACGACGCAAGGGACCACAGGGAGCCCAGCAGECGGAGCAACGCCACAGCACGACACCGCAGAGGCA |
| HYC32 | CACGACGCAAGGGACCACAGGGACGGGGGACCEGCAGCCAACCAEACAGCACGACACCGCAGAGGCA |
| HYC33 | CACGACGCAAGGGACCACAGGACAAGGCACAGAGGAGAAGCECCEACAGCACGACACCGCAGAGGCA |
| HYC34 | CACGACGCAAGGGACCACAGGAAGACGGGEACEAEACCAAGCAAGGCAGCACGACACCGCAGAGGCA |
| HYC35 | CACGACGCAAGGGACCACAGGGCGACAAACGAAEGCCAAAAAAAGGCAGCACGACACCGCAGAGGCA |
| HYC36 | CACGACGCAAGGGACCACAGGCCGACAGGAGEAGCCGAGGGGEEGCCAGCACGACACCGCAGAGGCA |
| HYC37 | CACGACGCAAGGGACCACAGGGACAGAAAAGGGGAGGAGCCCCAGACAGCACGACACCGCAGAGGCA |
| HYC38 | CACGACGCAAGGGACCACAGGGGCGGGEACGCEGGCCCAAGACAGACAGCACGACACCGCAGAGGCA |
| HYC39 | CACGACGCAAGGGACCACAGGCAGCACACGCAEGGCAGGGAGCCGECAGCACGACACCGCAGAGGCA |
| HYC40 | CACGACGCAAGGGACCACAGGAAGGAGGAAGACCCGCGECCAACEGCAGCACGACACCGCAGAGGCA |

Note: E stands for Indole-modified EdU.

**Supplementary Table S5.** Sequence information of siRNA.

| Target | siRNA | Sequence (5′-3′) |
| --- | --- | --- |
| ASCC3-siRNA1 | hASCC3-52-s | GCA AGA UAA UUA UAA UGA A |
|  | hASCC3-52-a | UUC AUU AUA AUU AUC UUG C |
| ASCC3-siRNA2 | ASCC3-190-s | AAG UAU AAA UGA AGA CUU A |
|  | ASCC3-190-a | UAA GUC UUC AUU UAU ACU U |
| ASCC3-siRNA3 | hASCC3-94-s | GCG AUC UAA ACU UCA UGA A |
|  | hASCC3-94-a | UUC AUG AAG UUU AGA UCG C |
| IMPDH1-siRNA1 | hIMPDH1-369-s | CAG GAU UCA UAG ACU UCA U |
|  | hIMPDH1-369-a | AUG AAG UCU AUG AAU CCU G |
| IMPDH1-siRNA2 | hIMPDH1-567-s | GGA AGG UCA AGA AGU UUG A |
|  | hIMPDH1-567-a | UCA AAC UUC UUG ACC UUC C |
| IMPDH1-siRNA3 | hIMPDH1-459-s | CCA UGG ACA CUG UGA CAG A |
|  | hIMPDH1-459-a | UCU GUC ACA GUG UCC AUG G |
| GTSE1-siRNA1 | hGTSE1-173-s | GGA CAU AAA GAA AGA UGU A |
|  | hGTSE1-173-a | UAC AUC UUU CUU UAU GUC C |
| GTSE1-siRNA2 | hGTSE1-292-s | CGU GGA GGU GUA CAA AGA A |
|  | hGTSE1-292-a | UUC UUU GUA CAC CUC CAC G |
| GTSE1-siRNA3 | hGTSE1-58-s | GGA UGA CCC UAA GAA GGA A |
|  | hGTSE1-58-a | UUC CUU CUU AGG GUC AUC C |
| ALG3-siRNA1 | hALG3-305-s | GGU UUC GUG UAC AUC UUU A |
|  | hALG3-305-a | UAA AGA UGU ACA CGA AAC C |
| ALG3-siRNA2 | hALG3-50-s | GAG GGA CUC UGC AAG CAA U |
|  | hALG3-50-a | AUU GCU UGC AGA GUC CCU C |
| ALG3-siRNA3 | hALG3-244-s | CAA UGG UAC CUA UGA CUA U |
|  | hALG3-244-a | AUA GUC AUA GGU ACC AUU G |
| ABCA2-siRNA1 | hABCA2-183-s | CCCUGGUGCUGUUCUUUAU |
|  | hABCA2-183-a | AUAAAGAACAGCACCAGGG |
| ABCA2-siRNA2 | hABCA2-1944-s | CGCCUCACGUGCACUACAA |
|  | hABCA2-1944-a | UUGUAGUGCACGUGAGGCG |
| ABCA2-siRNA3 | hABCA2-1785-s | GCGGCUGGAUCCAGUUCAU |
|  | hABCA2-1785-a | AUGAACUGGAUCCAGCCGC |
| PTPN3-siRNA1 | hPTPN3-887-s | GAAGUUUCUCAGAACCGAA |
|  | hPTPN3-887-a | UUCGGUUCUGAGAAACUUC |
| PTPN3-siRNA2 | hPTPN3-42-s | CAGUGGUUCUAGCGUCCUA |
|  | hPTPN3-42-a | UAGGACGCUAGAACCACUG |
| PTPN3-siRNA3 | hPTPN3-412-s | CAUACAUCAGCGACAGAAA |
|  | hPTPN3-412-a | UUUCUGUCGCUGAUGUAUG |
| NF2-siRNA1 | hNF2-1-s | CUACUUUGCAAUCCGGAAU |
|  | hNF2-1-a | AUUCCGGAUUGCAAAGUAG |
| NF2-siRNA2 | hNF2-2-s | GGAGUUUACUAUUAAACCA |
|  | hNF2-2-a | UGGUUUAAUAGUAAACUCC |
| NF2-siRNA3 | hNF2-93-s | AGAUGGAGUUCAAUUGCGA |
|  | hNF2-93-a | UCGCAAUUGAACUCCAUCU |
| MTCL1-siRNA1 | hMTCL1-1372-s | GCA GAU GAU UGA AGU GGA A |
|  | hMTCL1-1372-a | UUC CAC UUC AAU CAU CUG C |
| MTCL1-siRNA2 | hMTCL1-1032-s | CGG AGA ACG ACU AUC UCA A |
|  | hMTCL1-1032-a | UUG AGA UAG UCG UUC UCC G |
| MTCL1-siRNA3 | hMTCL1-1087-s | GAU GAG AGA CAG UUA UUU A |
|  | hMTCL1-1087-a | UAA AUA ACU GUC UCU CAU C |
| MSN-siRNA1 | hMSN-171-s | GGC UGA AAC UCA AUA AGA A |
|  | hMSN-171-a | UUC UUA UUG AGU UUC AGC C |
| MSN-siRNA2 | hMSN-378-s | CGU AUG CUG UCC AGU CUA A |
|  | hMSN-378-a | UUA GAC UGG ACA GCA UAC G |
| MSN-siRNA3 | hMSN-126-s | GGU UCU UUG GUC UGC AGU A |
|  | hMSN-126-a | UAC UGC AGA CCA AAG AAC C |
| SNX9-siRNA1 | hSNX9-697-s | CAUCAUUGUUGGAGAUUAU |
|  | hSNX9-697-a | AUAAUCUCCAACAAUGAUG |
| SNX9-siRNA2 | hSNX9-577-s | CAGUCGUGCUAGUUCCUCA |
|  | hSNX9-577-a | UGAGGAACUAGCACGACUG |
| SNX9-siRNA3 | hSNX9-830-s | AACACUAAUCGAUCUGUAA |
|  | hSNX9-830-a | UUACAGAUCGAUUAGUGUU |
| NC-siRNA | NC-siRNA-s | UUCUCCGAACGUGUCACGU |
|  | NC-siRNA-a | ACGUGACACGUUCGGAGAA |

**Supplementary Table S6.** Sequence information of Primers for qPCR.

| Target | Primer | Sequence (5′-3′) |
| --- | --- | --- |
| ASCC3 | Forward primer | CAGGCCGAGAGTTTCAGGT |
|  | Reverse primer | TCGCTTTATCTTTAAGTCAGCCAC |
| IMPDH1 | Forward primer | TGAAGAAGAACCGAGACTACCC |
|  | Reverse primer | TCCAGACGGTATTTGTCATCCT |
| GTSE1 | Forward primer | TCTGCCGCTTTGGTGACTT |
|  | Reverse primer | GGTCATCCATGTTCACGTCC |
| ALG3 | Forward primer | AGGTTCATTTTTCTCAGCATCCG |
|  | Reverse primer | CAGTCAATCTCTGTGTCTCCC |
| PTPN3 | Forward primer | GGTGCAGACATCAAGCCAGT |
|  | Reverse primer | GGTTCCTCTTGCTGCTTCCA |
| ABCA2 | Forward primer | GAAGAACGTGACGCTCAAACG |
|  | Reverse primer | CCGCTGTGTAGAAGGAGACTTCC |
| NF2 | Forward primer | GAAGAGGAGCTGGTTCAGGAG |
|  | Reverse primer | GGTCGTAGTCACCATACTTGGC |
| MTCL1 | Forward primer | GGGCCCATGGAGGAACTAC |
|  | Reverse primer | GTGAGGCAGTGGGGGAATG |
| MSN | Forward primer | AGCAGGAAGCCTAACAGTCG |
|  | Reverse primer | ACGCACACTGATCGTTTTGG |
| SNX9 | Forward primer | CATGGCCACCAAGGCTCG |
|  | Reverse primer | TCCAGCCATCCTCCACCTAC |
| ACTB | Forward primer | CATGTACGTTGCTATCCAGGC |
|  | Reverse primer | CTCCTTAATGTCACGCACGAT |

**Supplementary Table S7.** IndU substitution and truncation sequences of XH3C (E stands for Indole-modified EdU)

| ID | Sequence (5′-3′) |
| --- | --- |
| HYC4 | CACGACGCAAGGGACCACAGGGGCEGGGGCAAACACACACCCGAAACAGCACGACACCGCAGAGGCA |
| HYC4 All T | CACGACGCAAGGGACCACAGGGGCTGGGGCAAACACACACCCGAAACAGCACGACACCGCAGAGGCA |
| HYC4 Truc1 | CGCAAGGGACCACAGGGGCEGGGGCAAACACACACCCGAAACAGCACGACACCGCAG |
| HYC4 Truc2 | CGCAAGGGACCACAGGGGCEGGGGCAAACACACACCCGAAACAGCACGA |
| HYC4 Truc3 | CGCAAGGGACCACAGGGGCEGGGGCAAACACACACCCGAAA |
| HYC4 Truc4 | ACCACAGGGGCEGGGGCAAACACACACCCGAAA |
| HYC15 | CACGACGCAAGGGACCACAGGCGGAAGACCACGACCCAEAACCCEGCAGCACGACACCGCAGAGGCA |
| HYC15 Site1 sub | CACGACGCAAGGGACCACAGGCGGAAGACCACGACCCATAACCCEGCAGCACGACACCGCAGAGGCA |
| HYC15 Site2 sub | CACGACGCAAGGGACCACAGGCGGAAGACCACGACCCAEAACCCTGCAGCACGACACCGCAGAGGCA |
| HYC15 All T | CACGACGCAAGGGACCACAGGCGGAAGACCACGACCCATAACCCTGCAGCACGACACCGCAGAGGCA |
| HYC15 Truc1 | CGCAAGGGACCACAGGCGGAAGACCACGACCCAEAACCCEGCAGCACGACACCGCAG |
| HYC15 Truc2 | CGCAAGGGACCACAGGCGGAAGACCACGACCCAEAACCCEGCAGCACGA |
| HYC15 Truc3 | CGCAAGGGACCACAGGCGGAAGACCACGACCCAEAACCCEG |
| HYC15 Truc4 | ACCACAGGCGGAAGACCACGACCCAEAACCCEG |
| HYC24 | CACGACGCAAGGGACCACAGGAGGAAGEACGCAAEAGCGGGAGGACCAGCACGACACCGCAGAGGCA |
| HYC24 Site1 sub | CACGACGCAAGGGACCACAGGAGGAAGTACGCAAEAGCGGGAGGACCAGCACGACACCGCAGAGGCA |
| HYC24 Site2 sub | CACGACGCAAGGGACCACAGGAGGAAGEACGCAATAGCGGGAGGACCAGCACGACACCGCAGAGGCA |
| HYC24 All T | CACGACGCAAGGGACCACAGGAGGAAGTACGCAATAGCGGGAGGACCAGCACGACACCGCAGAGGCA |
| HYC24 Truc1 | CGCAAGGGACCACAGGAGGAAGEACGCAAEAGCGGGAGGACCAGCACGACACCGCAG |
| HYC24 Truc2 | CGCAAGGGACCACAGGAGGAAGEACGCAAEAGCGGGAGGACCAGCACGA |
| HYC24 Truc3 | CGCAAGGGACCACAGGAGGAAGEACGCAAEAGCGGGAGGAC |
| HYC24 Truc4 | ACCACAGGAGGAAGEACGCAAEAGCGGGAGGAC |
| HYC34 | CACGACGCAAGGGACCACAGGAAGACGGGEACEAEACCAAGCAAGGCAGCACGACACCGCAGAGGCA |
| HYC34 Site1 sub | CACGACGCAAGGGACCACAGGAAGACGGGTACEAEACCAAGCAAGGCAGCACGACACCGCAGAGGCA |
| HYC34 Site2 sub | CACGACGCAAGGGACCACAGGAAGACGGGEACTAEACCAAGCAAGGCAGCACGACACCGCAGAGGCA |
| HYC34 Site3 sub | CACGACGCAAGGGACCACAGGAAGACGGGEACEATACCAAGCAAGGCAGCACGACACCGCAGAGGCA |
| HYC34 All T | CACGACGCAAGGGACCACAGGAAGACGGGTACTATACCAAGCAAGGCAGCACGACACCGCAGAGGCA |
| HYC34 Truc1 | CGCAAGGGACCACAGGAAGACGGGEACEAEACCAAGCAAGGCAGCACGACACCGCAG |
| HYC34 Truc2 | CGCAAGGGACCACAGGAAGACGGGEACEAEACCAAGCAAGGCAGCACGA |
| HYC34 Truc3 | CGCAAGGGACCACAGGAAGACGGGEACEAEACCAAGCAAGG |
| HYC34 Truc4 | ACCACAGGAAGACGGGEACEAEACCAAGCAAGG |
| HYC35 | CACGACGCAAGGGACCACAGGGCGACAAACGAAEGCCAAAAAAAGGCAGCACGACACCGCAGAGGCA |
| HYC35 All T | CACGACGCAAGGGACCACAGGGCGACAAACGAATGCCAAAAAAAGGCAGCACGACACCGCAGAGGCA |
| HYC35 Truc1 | CGCAAGGGACCACAGGGCGACAAACGAAEGCCAAAAAAAGGCAGCACGACACCGCAG |
| HYC35 Truc2 | CGCAAGGGACCACAGGGCGACAAACGAAEGCCAAAAAAAGGCAGCACGA |
| HYC35 Truc3 | CGCAAGGGACCACAGGGCGACAAACGAAEGCCAAAAAAAGG |
| HYC35 Truc4 | ACCACAGGGCGACAAACGAAEGCCAAAAAAAGG |
| HYC8 | CACGACGCAAGGGACCACAGGGGAGCACAECCEECCGGAGCEGGAGCAGCACGACACCGCAGAGGCA |
| HYC8 Site1 sub | CACGACGCAAGGGACCACAGGGGAGCACATCCEECCGGAGCEGGAGCAGCACGACACCGCAGAGGCA |
| HYC8 Site2 sub | CACGACGCAAGGGACCACAGGGGAGCACAECCTECCGGAGCEGGAGCAGCACGACACCGCAGAGGCA |
| HYC8 Site3 sub | CACGACGCAAGGGACCACAGGGGAGCACAECCETCCGGAGCEGGAGCAGCACGACACCGCAGAGGCA |
| HYC8 Site4 sub | CACGACGCAAGGGACCACAGGGGAGCACAECCEECCGGAGCTGGAGCAGCACGACACCGCAGAGGCA |
| HYC8 All T | CACGACGCAAGGGACCACAGGGGAGCACATCCTTCCGGAGCTGGAGCAGCACGACACCGCAGAGGCA |
| HYC8 Truc1 | CGCAAGGGACCACAGGGGAGCACAECCEECCGGAGCEGGAGCAGCACGACACCGCAG |
| HYC8 Truc2 | CGCAAGGGACCACAGGGGAGCACAECCEECCGGAGCEGGAGCAGCACGA |
| HYC8 Truc3 | CGCAAGGGACCACAGGGGAGCACAECCEECCGGAGCEGGAG |
| HYC8 Truc4 | ACCACAGGGGAGCACAECCEECCGGAGCEGGAG |
| HYC40 | CACGACGCAAGGGACCACAGGAAGGAGGAAGACCCGCGECCAACEGCAGCACGACACCGCAGAGGCA |
| HYC40 Site1 sub | CACGACGCAAGGGACCACAGGAAGGAGGAAGACCCGCGTCCAACEGCAGCACGACACCGCAGAGGCA |
| HYC40 Site2 sub | CACGACGCAAGGGACCACAGGAAGGAGGAAGACCCGCGECCAACTGCAGCACGACACCGCAGAGGCA |
| HYC40 All T | CACGACGCAAGGGACCACAGGAAGGAGGAAGACCCGCGTCCAACTGCAGCACGACACCGCAGAGGCA |
| HYC40 Truc1 | CGCAAGGGACCACAGGAAGGAGGAAGACCCGCGECCAACEGCAGCACGACACCGCAG |
| HYC40 Truc2 | CGCAAGGGACCACAGGAAGGAGGAAGACCCGCGECCAACEGCAGCACGA |
| HYC40 Truc3 | CGCAAGGGACCACAGGAAGGAGGAAGACCCGCGECCAACEG |
| HYC40 Truc4 | ACCACAGGAAGGAGGAAGACCCGCGECCAACEG |

Note: E stands for Indole-modified EdU.

**Supplementary Table S8.** Cluster and enrichment information of aptamer libraries for high-throughput phenotypic screening. CPM means Counts Per Million.

| ID | Cluster ID | PD3 CPM | PD2 CPM | PD1 CPM | Pre-enrich CPM |
| --- | --- | --- | --- | --- | --- |
| HYC1 | 91972 | 4581.794 | 2031.045 | 442.025 | 9.034 |
| HYC2 | 41715 | 4081.588 | 1939.22 | 637.844 | 9.369 |
| HYC3 | 86025 | 4036.105 | 1719.015 | 707.36 | 8.432 |
| HYC4 | 140110 | 3251.977 | 1723.43 | 398.53 | 7.294 |
| HYC5 | 53549 | 3538.817 | 3747.609 | 732.783 | 8.733 |
| HYC6 | 242468 | 3635.617 | 3299.907 | 1135.426 | 7.06 |
| HYC7 | 47137 | 3599.877 | 2315.634 | 607.745 | 8.633 |
| HYC8 | 219877 | 3075.421 | 2740.897 | 479.086 | 7.796 |
| HYC9 | 229311 | 2825.936 | 2352.939 | 428.347 | 7.629 |
| HYC10 | 187997 | 2854.868 | 1908.862 | 190.404 | 5.253 |
| HYC11 | 66059 | 3006.184 | 1352.829 | 538.511 | 7.662 |
| HYC12 | 214697 | 2919.011 | 3059.894 | 417.834 | 5.655 |
| HYC13 | 366283 | 2576.435 | 1836.196 | 583.097 | 6.625 |
| HYC14 | 156325 | 2787.724 | 2167.203 | 296.7 | 7.897 |
| HYC15 | 303588 | 2649.068 | 2604.131 | 266.812 | 6.257 |
| HYC16 | 46715 | 2619.972 | 2958.004 | 408.797 | 8.8 |
| HYC17 | 366448 | 2502.714 | 1362.124 | 210.482 | 5.22 |
| HYC18 | 11089 | 2458.204 | 1774.79 | 659.012 | 5.755 |
| HYC19 | 53751 | 2385.851 | 2934.734 | 567.414 | 7.83 |
| HYC20 | 206575 | 2377.839 | 2636.9 | 257.81 | 7.094 |
| HYC21 | 368782 | 2339.231 | 1691.896 | 422.581 | 5.421 |
| HYC22 | 282244 | 2356.277 | 2780.186 | 406.3 | 6.96 |
| HYC23 | 235929 | 2280.891 | 959.038 | 203.414 | 5.253 |
| HYC24 | 262563 | 2316.416 | 980.891 | 473.461 | 4.818 |
| HYC25 | 114765 | 2132.278 | 2834.504 | 245.925 | 6.123 |
| HYC26 | 153346 | 2154.928 | 1771.428 | 320.61 | 5.086 |
| HYC27 | 101957 | 1303.062 | 1092.118 | 104.256 | 3.112 |
| HYC28 | 173356 | 2156.132 | 1883.222 | 363.438 | 5.354 |
| HYC29 | 182867 | 2107.616 | 2234.725 | 839.57 | 6.491 |
| HYC30 | 137215 | 2141.476 | 2604.658 | 892.595 | 6.391 |
| HYC31 | 124435 | 2127.035 | 1869.309 | 468.925 | 6.391 |
| HYC32 | 416577 | 1693.692 | 1084.199 | 112.66 | 3.346 |
| HYC33 | 17218 | 1305.766 | 1505.269 | 194.905 | 3.58 |
| HYC34 | 68813 | 1758.495 | 1765.818 | 320.153 | 5.287 |
| HYC35 | 77425 | 1899.261 | 1850.19 | 184.567 | 5.688 |
| HYC36 | 59618 | 2070.393 | 2157.016 | 259.498 | 5.688 |
| HYC37 | 121692 | 2070.541 | 923.05 | 250.074 | 3.446 |
| HYC38 | 176456 | 1981.555 | 1717.192 | 601.205 | 4.885 |
| HYC39 | 79142 | 1973.263 | 850.485 | 257.669 | 4.182 |
| HYC40 | 257002 | 1968.499 | 1130.922 | 299.794 | 4.617 |

**Supplementary Table S9.** Sequence information of aptamer for other experiment.

| ID | Sequence (5′-3′) |
| --- | --- |
| Aptamer-CD63 | 5′-CACCCCACCTCGCTCCCGTGACACTAATGCTA-3′ |
| Random Sequence | GTGACGCAGCAGCTGCCTGTACATGGGCTATCTGGCTGGACAC |
| TBA29 | AGTCCGTGGTAGGGCAGGTTGGGGTGACT |

**Supplementary Table S10.** Characteristics of Selected Aptamers and Their Validated Protein Targets.

| **Aptamer ID** | **Uniport ID** | **Gene name** | **Protein name** | **Membrane connection** | **Liver cancer-related** | **ExoCarta** |
| --- | --- | --- | --- | --- | --- | --- |
| HYC4 | Q9NYZ3 | GTSE1 | G2 and S phase-expressed protein 1 | √ | √ | √ |
| HYC15 | P26045 | PTPN3 | Tyrosine-protein phosphatase non-receptor type 3 | √ | √ | × |
| HYC24 | P35240 | NF2 | Merlin | √ | √ | √ |
| HYC34 | Q8N3C0 | ASCC3 | Activating signal cointegrator 1 complex subunit 3 | √ | √ | √ |
| HYC35 | P20839 | IMPDH1 | Inosine-5'-monophosphate dehydrogenase 1 | √ | √ | √ |
| HYC8 | Q92685 | ALG3 | Dol-P-Man:Man(5)GlcNAc(2)-PP-Dol alpha-1,3-mannosyltransferase | √ | √ | √ |
| HYC40 | Q9BZC7 | ABCA2 | ATP-binding cassette sub-family A member 2 | √ | × | √ |
| HYC5 | Q9Y5X1 | SNX9 | Sorting nexin-9 | √ | × | √ |
| HYC9 | P26038 | MSN | Moesin | √ | √ | √ |
| HYC11 | Q9Y4B5 | MTCL1 | Microtubule cross-linking factor 1 | √ | √ | √ |
